# Inhibitors of the Small Membrane (M) Protein Viroporin Prevent Zika Virus Infection

**DOI:** 10.1101/2021.03.11.435022

**Authors:** Emma Brown, Gemma Swinscoe, Daniella Lefteri, Ravi Singh, Amy Moran, Rebecca Thompson, Daniel Maskell, Hannah Beaumont, Matthew Bentham, Claire Donald, Alain Kohl, Andrew Macdonald, Neil Ranson, Richard Foster, Clive S. McKimmie, Antreas C. Kalli, Stephen Griffin

## Abstract

*Flaviviruses*, including Zika virus (ZIKV), are a significant global health concern, yet no licensed antivirals exist to treat disease. The small Membrane (M) protein plays well-defined roles during viral egress and remains within virion membranes following release and maturation. However, it is unclear whether M plays a functional role in this setting. Here, we show that M forms oligomeric membrane-permeabilising channels *in vitro*, with increased activity at acidic pH and sensitivity to the prototypic channel-blocker, rimantadine. Accordingly, rimantadine blocked an early stage of ZIKV cell culture infection.

Structure-based channel models, comprising hexameric arrangements of two *trans*-membrane domain protomers were shown to comprise more stable assemblages than other oligomers using molecular dynamics (MD) simulations. Models contained a predicted lumenal rimantadine binding site, as well as a second druggable target region on the membrane-exposed periphery. *In silico* screening enriched for repurposed drugs/compounds predicted to bind to either one site or the other. Hits displayed superior potency *in vitro* and in cell culture compared with rimantadine, with efficacy demonstrably linked to virion-resident channels. Finally, rimantadine effectively blocked ZIKV viraemia in preclinical models, supporting that M constitutes a physiologically relevant target. This could be explored by repurposing rimantadine, or development of new M-targeted-therapies.

## Introduction

Zika virus (ZIKV) was first isolated in the eponymous Ugandan forest in 1947 and is a mosquito - borne arbovirus (<u>ar</u>thropod-<u>bo</u>rne), classified within the *Flavivirus* genus of the *Flaviviridae*^1,2^. Historically, ZIKV caused infrequent and relatively mild infections in humans with symptoms comprising fever, rash, headaches, joint/muscle pain and conjunctivitis^3^. However, once confined to Africa and Southeast Asia, a large outbreak in 2007 on Yap Island (Micronesia), followed by further outbreaks in French Polynesia, New Caledonia, and the Cook Islands in 2013/14, signified spread of the virus across the Pacific. The virus then arrived in Brazil in 2015, where a n epidemic affecting >1M people ensued. This resulted in worldwide spread of the virus involving more than eighty countries and signified emergent ZIKV as a global health concern ^4^.

Worryingly, the outbreaks in Yap and thereafter encompassed considerably more serious symptomology, which has been linked to evolution of New World ZIKV strains^5^. This included more serious and frequent neurological complications, increased incidence of Guillian-Barre Syndrome, and a devastating epidemic of microcephaly in new-borns where maternal infection had occurred during the first trimester of pregnancy^6–8^. Eventually, the outbreak was contained by surveillance combined with control of insect vectors, predominantly *Aedes aegypti* and *Aedes albopictus* mosquitos^9^. More than seven years later, a ZIKV vaccine remains elusive, although several are in trials (reviewed in ^10^). Similarly, no licensed antiviral therapies exist to treat ZIKV infection, yet repurposing of nucleotide analogue inhibitors of the NS5 protein shows promise ^11–13^.

ZIKV, like other *Flaviviruses*, possesses a positive sense single-stranded RNA (ssRNA) genome over ten kilobases in length, capped at the 5ʹ end by a methylated dinucleotide (m^7^GpppAmp)^14,15^. The RNA is translated into a polyprotein that is subsequently cleaved by host and viral proteases to yield ten mature gene products^16^; these are spatially organised into structural virion components at the N-terminus, and non-structural replicase proteins at the C-terminus. The mature enveloped virion comprises the viral RNA, the capsid (C) protein and two membrane proteins: envelope (E) and the small membrane (M) protein^17,18^. Virions undergo maturation as they traffic through the cell during egress, with M and the precursor (pr) peptide acting as chaperones for the E protein ^19^. Maturation occurs upon furin-mediated cleavage of pr-M in the Golgi^20,21^, although this has variable efficiency^22,23^. The cleaved pr peptide remains associated with E, preventing exposure of the fusion loop within acidifying secretory vesicles. Release from the cell into a pH-neutral environment leads to the loss of pr and resultant structural rearrangement of E and M ^24^. Whilst E comprises the essential virus attachment and entry protein, the role of M within the mature infectious virion remains unknown^25^.

The presence of dimeric M, bound to E dimers^25^, within the virion raises the question of whether it might play a role during the virus entry process. Unfortunately, cryo -EM structures of acidified *Flavivirus* particles - a proxy for changes likely to occur during endocytosis - do not contain density attributable to the M protein^26^. Moreover, E undergoes a shift from a dimeric to a trimeric form thought to mediate membrane fusion^27–31^. This presumably disrupts E-M dimeric interactions, thereby allowing M an opportunity to also adopt alternative conformations and explore other oligomeric states.

Interestingly, dengue virus (DENV) M peptides formed membrane channels *in vitro*^32^, which would require the formation of higher order oligomers. However, electrophysiology studies in *Xenopus* oocytes failed to detect M mediated proton conductance^33^. Nevertheless, due to its size and hydrophobicity, M has long been speculated to comprise a virus -encoded ion channel, or “viroporin”, typified by the M2 proton channel of influenza A virus (IAV)^34–36^. M2 functions to allow protons to pass into the virion core during virus entry, enabling the uncoating of viral RNA, and is the target of the antiviral drugs amantadine and rimantadine, setting clinical precedent for viroporins as drug targets.

Here, we provide evidence that ZIKV M forms rimantadine-sensitive, oligomeric membrane channels whose activity increases at acidic pH. Rimantadine blocks an early step during the ZIKV life cycle, consistent with a role for M channels during virus entry. Critically, rimantadine also abrogates viraemia within *in vivo* ZIKV preclinical models, supporting that M channel activity is a physiologically relevant target. In-depth molecular dynamics investigations support that M channels are comprised of hexameric complexes, with the channel lumen lined by the C-terminal *trans*-membrane domain (TMD). Ensuing models provided an effective template for *in silico* high throughput screening (HTS) of a library of FDA-approved and other biologically active compounds, targeting two discrete sites on the channel complex. Pleasingly, enrichment of the library led to the identification of much - improved blockers of M channel activity, laying the foundation for potential drug repurposing alongside bespoke antiviral development.

## Results

### ZIKV M peptides form channel-like oligomeric complexes within membrane-mimetic environments

M protein resides as a dimeric form resolvable within cryo -EM structures of mature ZIKV, and other *Flavivirus* particles, in close association with E dimers^25^. Thus, M would be required to form higher order oligomers to form a putative channel across the membrane. This might conceivably occur during endocytosis where acidic pH induces reconfiguration of E into a trimeric fusion complex, presumably also liberating M dimers to move within the virion membrane ^27–31^. Thus, we hypothesised that M peptides should form channel structures within a lipidic environment in the absence of E protein.

A peptide lacking the disordered N-terminal 20 amino acids was synthesised based upon the M protein sequence from a New World ZIKV isolate (PE243)^37^. Peptides therefore comprised the three helical regions of the mature M protein, the second and third of which form TMDs within virion membranes (Figure 1a, S1, S2). Peptides were reconstituted using several non-ionic detergents used frequently as membrane-mimetics in structural and/or biophysical studies^38–41^, as well as in our previous investigations of viroporin oligomerisation^42–45^.

**Figure 1.**
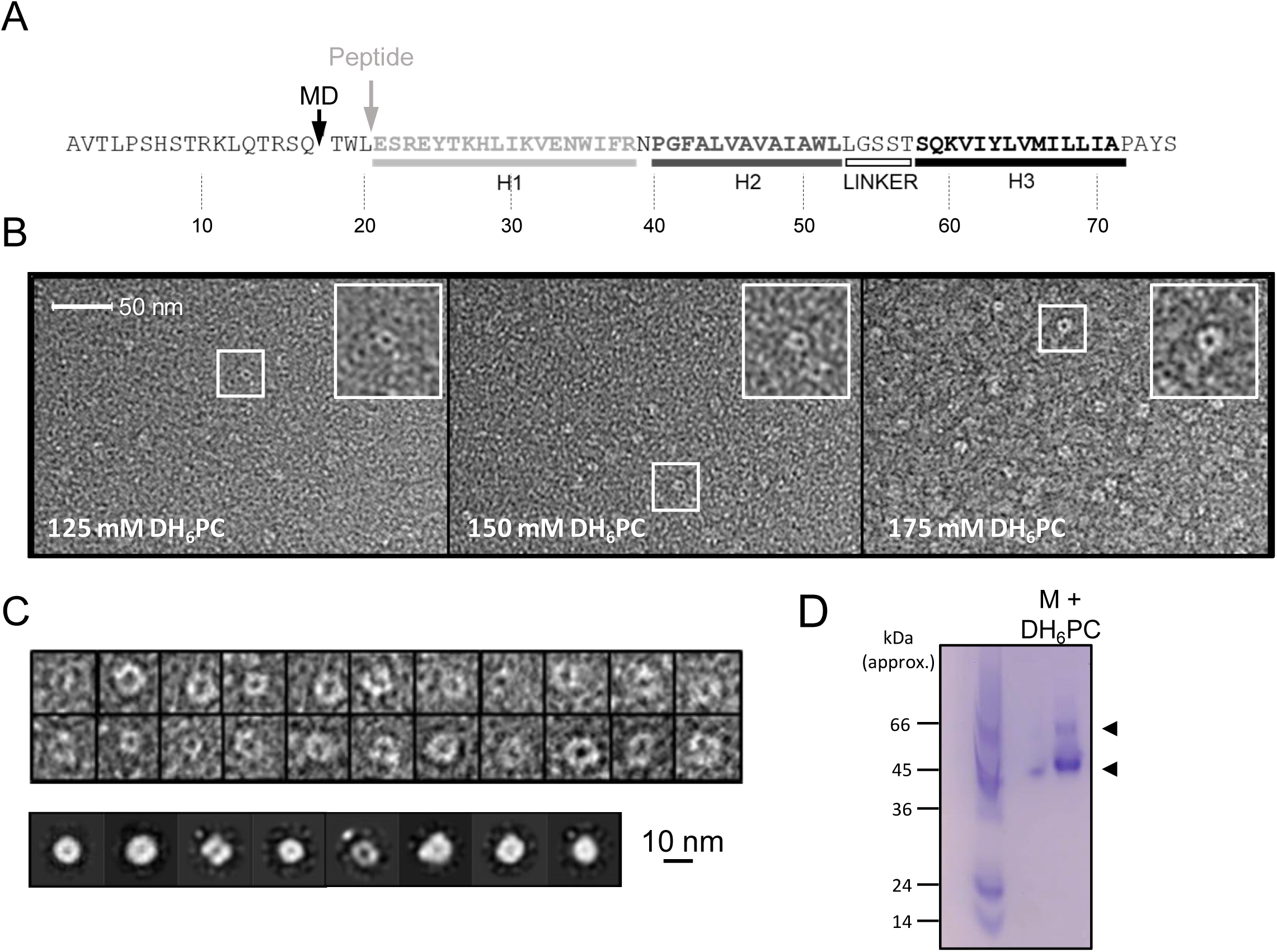
*In vitro characterisation of M*. **A.** M sequence showing helical regions and peptide truncation (*in vitro* synthesis (grey arrow) and MD simulations (black arrow)). **B**. Visualisation of M peptide with increasing concentrations of detergent, stained with uranyl acetate. Fields are representative of multiple images with ∼9000 particles collected in total across all conditions. Insets show zoomed images of particles with accumulation of stain within central cavity, consistent with channel formation. **C.** Top panel – examples of particles visualised at 150 mM illustrating heterogeneity. 9907 particles from 150 mM samples were resolved into 2D class averages with 25 iterations (bottom panel) yet rotational symmetry could not be determined. D. Native PAGE of M peptide (5 μg) reconstituted in DH_(6)_PC (300 mM).

To investigate the nature and stoichiometry of M complexes, DH _6_PC solubilised M peptides were first immobilised upon carbon grids and negatively stained using uranyl acetate over a range of detergent concentrations. Unfortunately, use of 300 mM detergent prevented the efficient adhesion of peptides, yet visualisation of M complexes at lower concentrations revealed the formation of circular structures, the majority of which were oriented approximately in parallel with the grid plane (Figure 1b).

Interestingly, complexes exhibited a significant degree of structural heterogeneity ( Figure 1c, top panel), although separable class subsets were resolvable amongst >9000 complexes analysed at 150 mM detergent (Figure 1c, bottom panel). Unfortunately, the degree of heterogeneity and small size prevented determination of stoichiometry by 2D rotational averaging. Nevertheless, the absence of obvious aggregation and the formation of, albeit heterogeneous, channel-like structures, supports that M can adopt conformations corresponding to membrane-spanning channel complexes.

Visualisation of peptide-detergent complexes using native PAGE revealed that dihexanoyl-phosphatidylcholine (DH_6_PC, 300 mM) induced the formation of higher order oligomers ( Figure 1d). A dominant species of ∼45-50 kDa formed along with another less abundant assemblage in the region of ∼60 kDa, although it is not possible to assign precise molecular weights by this method. However, this was consistent with the potential formation of hexameric and/or heptameric oligomers, confirming that dimers were not the only multimeric conformation adopted by free M protein.

### ZIKV M peptides exhibit acid-enhanced channel activity with sensitivity to rimantadine

Given the formation of higher order oligomers, we next assessed whether M peptides exhibited membrane permeabilisation using an indirect assay for channel formation, based upon the rele ase and unquenching of fluorescent dye from liposomes. This assay has been used to characterise the activity of several other viroporins and can be adapted to both explore aspects of channel biochemistry as well as identifying small molecule inhibitors of channel activity^42–51^.

Dose-dependent dye release occurred using sub-micromolar concentrations of M peptide, consistent with channel forming activity (Figure 2a); 780 nM was chosen as an optimal concentration for ensuing investigations. Interestingly, mildly acidic pH conditions increased M-mediated release of carboxyfluorescein (CF), reminiscent of influenza A virus (IAV) M2 ^48^, hepatitis C virus (HCV) p7^50,52^ and human papillomavirus type 16 (HPV16) E5^44^ in similar experiments (Figure 2b). Such conditions mimic those found within endocytic vesicles, consistent with M channels potentially being activated during virus entry.

**Figure 2.**
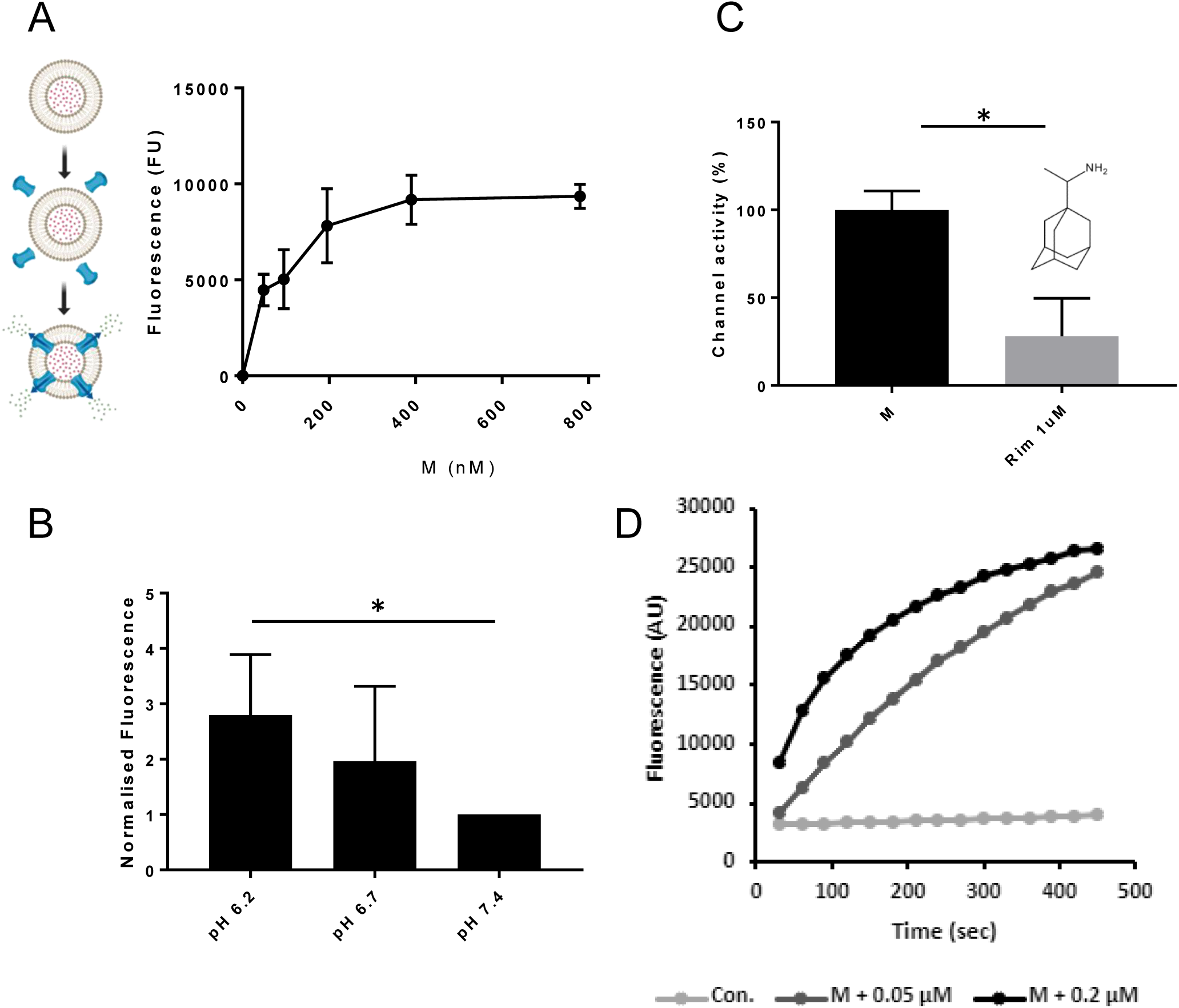
M induces rimantadine sensitive membrane permeability, augmented by reduced pH. **A.** Titration of DMSO-reconstituted M peptide in endpoint liposome CF release assay. Graph represents a single biological repeat representative of at least three others, comprising triplicate technical repeats at each concentration. Error bars show standard deviation. **B.** Modified endpoint CF assay undertaken with altered external buffer pH as indicated. CF content of re -buffered clarified assay supernatants were detected as a single endpoint measurement. Data comprise three biological repeats for each condition, error bars show normalised standard error, * p≤0.05, Student T-test. **C.** Effect of rimantadine (1 μM) upon M activity. Results are from three biological repeats normalised to 100 % activity for solvent control. Error bars show adjusted % error. * p≤0.05, Student T-test. **D**. Real-time CF assay data for M peptides in “virion-sized” liposomes, produced by extrusion through a 0.05 μm, rather than a 0.2 μm filter.

We tested whether M channel activity was sensitive to inhibitory small molecules, indicative of the formation of structurally ordered complexes. Prototypic channel blockers can be used to identify druggable sites within viroporin complexes as a result of their binding promiscuity. Regions so identified are then amenable to the development of improved small molecule inhibitors^46,48,49^. One such prototype, rimantadine (1 μM), caused a significant reduction in M channel activity (Figure 2c), consistent with the presence of a viable binding site within a soluble, folded M channel complex.

Finally, we tested whether M peptides were able to form channels within membranes exhibiting significant curvature, such as that seen within an enveloped virion. Liposomes extruded using a 0.05 μM filter provided a similar level of M-induced dye release, albeit with a slightly reduced kinetic, likely due to the increased molar ratio of resulting liposomes to peptide (Figure 2d).

### Rimantadine inhibits ZIKV infectivity in cell culture

We hypothesised that if M protein-mediated channel activity observed *in vitro* was relevant during the ZIKV life cycle, then rimantadine would exert an antiviral effect. We therefore tested the ability of rimantadine to inhibit ZIKV infection of Vero cells, adding the drug during infection and for the ensuing 24 hr period prior to analysis. Rimantadine exerted a dose-dependent reduction in ZIKV infection, assessed by both the number of cells infected as well as ensuing viral protein production (Figure 3a, b, c). Infected cell numbers underwent statistically significant reductions at rimantadine concentrations of 10 μM and above. Importantly, rimantadine affected neither cell viability, nor the uptake of fluorescently labelled EGF as a proxy marker for clathrin-mediated endocytosis over the concentration range tested (Fig S3).

**Figure 3.**
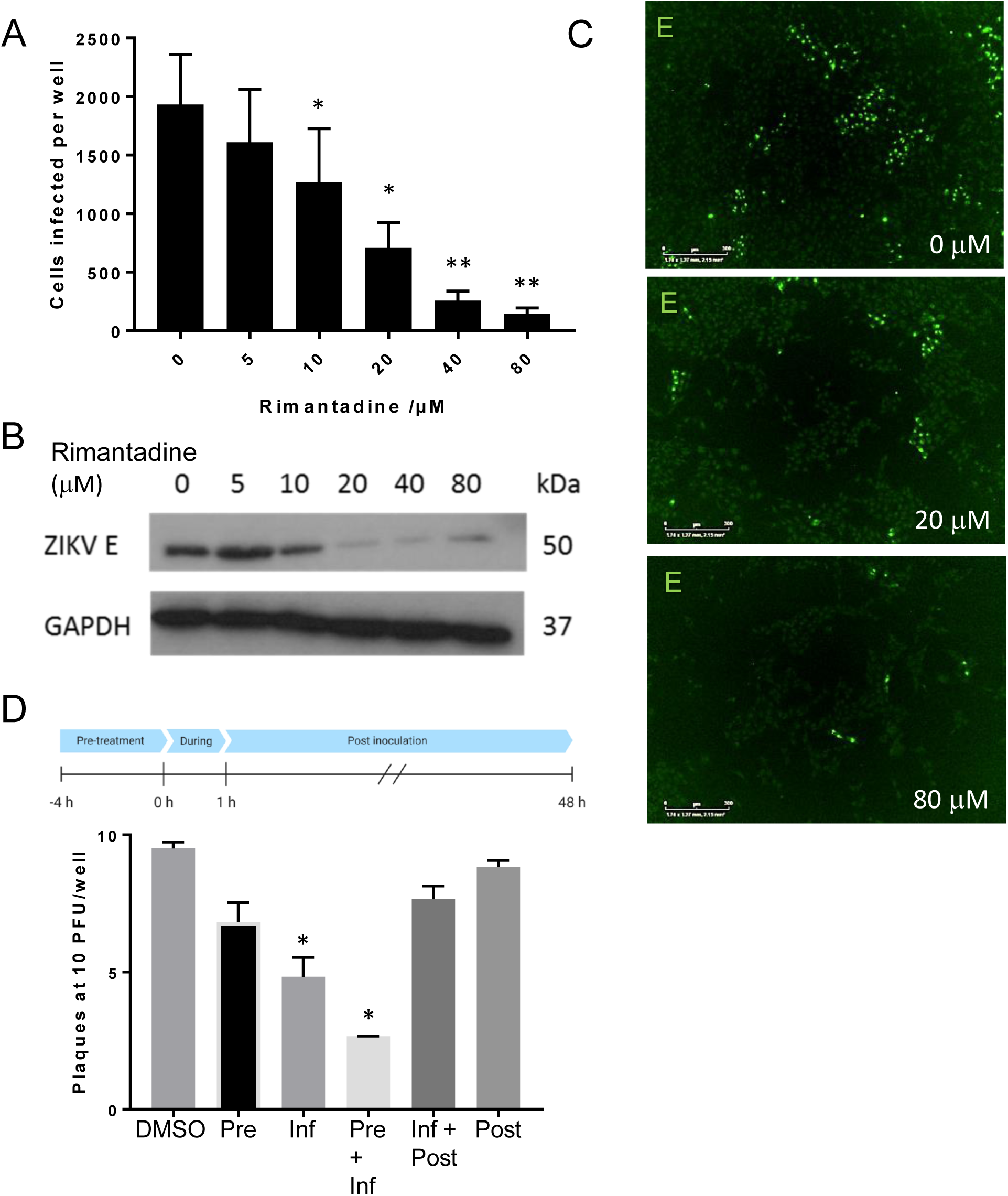
Rimantadine blockade of ZIKV infection. **A.** Vero cells were infected at an MOI of 0.1 pfu/cell with increasing rimantadine concentrations in 96 well plates. Infectious units were quantified 48 hr post-infection using an IncuCyte Zoom to count ZIKV infected cells stained with antibody to ZIKV E and an Alexafluor 488 nm secondary antibody. Counts were averaged over four image panels per well and results are from three biological replicates performed in triplicate. Error bars represent standard error of the mean between experiments. *p≤0.01, ** p≤0.001, paired student T-test relative to solvent control. **B.** Representative anti-E western blot of n = 3 biological repeats of 6-well plate experiments run in parallel to fluorescence-based assays. Infections and timings as for A. **C.** Example representative fluorescence images from IncuCyte analysis shown in A, scale bars represent 500 μm. **D.** Plaque reduction assay using BHK21 cells infected with 10 pfu/well in the presence of 80 μM rimantadine during different stages of infection shown by schematic. N = 2 biological replicates. Error bars represent standard error of the mean between experiments, *p≤0.01, paired student T-test relative to solvent control.

We next added rimantadine (80 μM) during different stages of the infection process, measuring drug effects by a BHK cell plaque-reduction assay, to determine the approximate stage of the ZIKV life cycle affected. Statistically significant reductions of infectivity occurred when rimantadine was present during infection, with more pronounced effects upon pre-incubating cells with the drug (Figure 3d). Measurable, but non-significant reductions also occurred upon adding rimantadine only prior to infection, or during and following infection. Taken together, rimantadine affected an early stage of the virus life cycle, coincident with virus entry.

### Molecular dynamics supports that M protomers adopt a double *trans*-membrane topology

The structure of M within mature ZIKV and other *Flavivirus* virion cryo-EM reconstructions is a dimeric complex of double-TMD protomers, closely associated with the TMDs of adjacent E dimers ^25^. The relatively short TMDs within M cause a pinching of the virion membrane bilayer, reducing its thickness by several nm^21,53,54^.

M is highly conserved between ZIKV isolates (Figure 4a), and regions including the linker region between the first and second alpha helices are shared with other *Flaviviruses* (Figure S2d). Where sequence conservation is absent, secondary structure and amino acid similarity are maintained. However, four different TMD prediction software packages revealed disparate outputs for the region known to form the second and third helices within cryo-EM reconstructions, with two favouring the formation of an extended single TMD (Figure S2a). Interestingly, the MEMSAT-SVM package not only predicted a double TMD topology, but also that helix three displayed properties consistent with being pore lining (Figure S2b).

**Figure 4.**
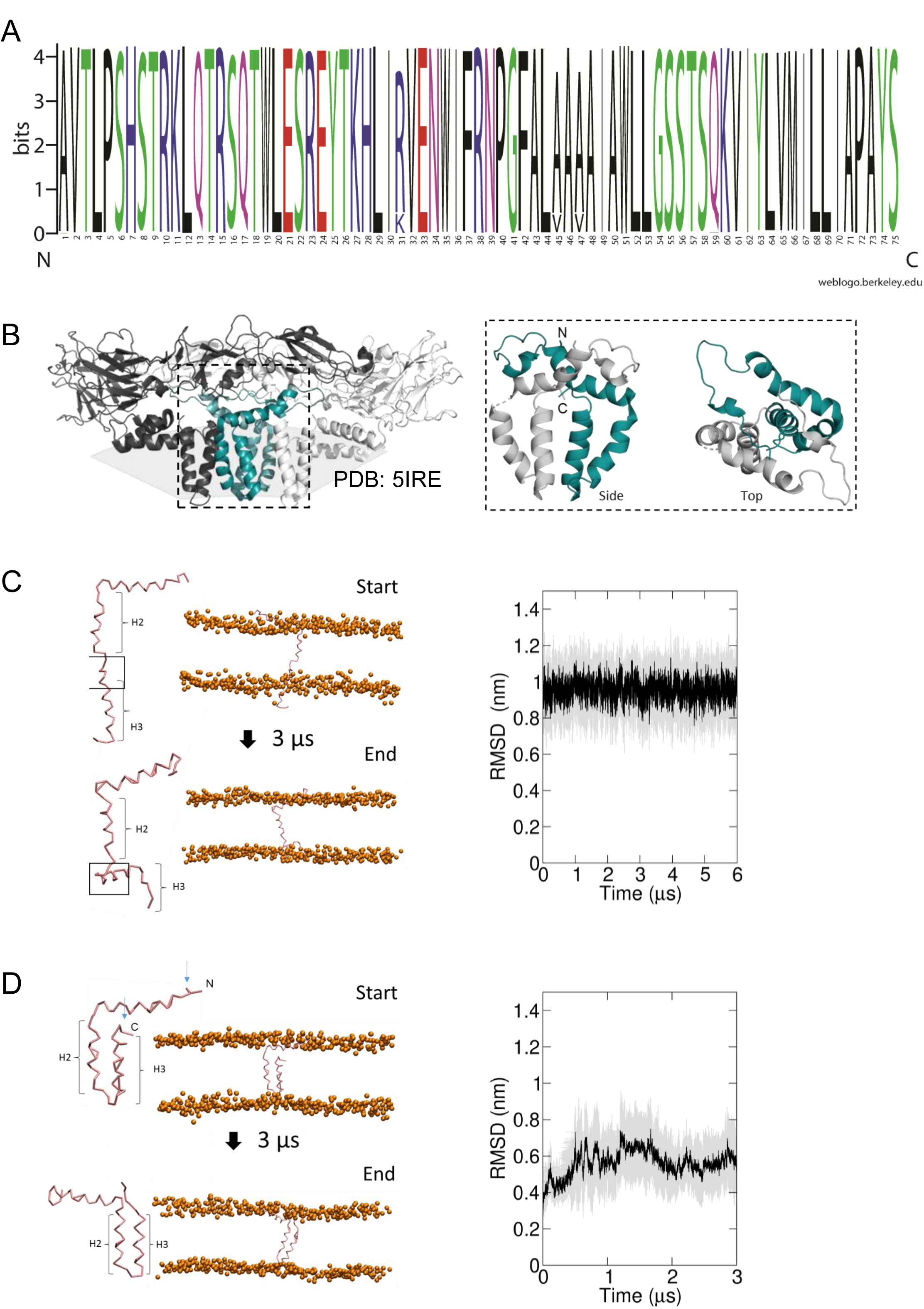
Predicted membrane topology for M protomers. **A.** M sequence “logo” (https://weblogo.berkeley.edu/logo.cgi) showing relative conservation at each amino acid position . 929 sequences were retrieved by a Uniprot search for “Zika virus”, then aligned using CLUSTAL Omega. Alignments were exported to Jalview where removal of partial/irrelevant sequences resulted in analysis of ∼700 sequences. Conservation is high (>95%) across the majority of the protein, with the exception of Arg31, Ala45 and Ala47, but even these remain present within >80% sequences. **B**. Image of an energy minimised M protomer taken from PDB: 5IRE within a simulated hydrated lipid membrane system (see methods). **C.** Straightened, single TM M protomer inserted into a (DOPC) bilayer and subjected to CG simulations for 3 μs, running from start (top) to finished (bottom) pose. RMSD over time (five repeat simulations, standard deviations in grey) showed a high degree of movement and flexibility exhibited by this conformation (∼1 nm throughout), as the protein appears to attempt moving the C-terminal helix (H3) back along the plain of the membrane. Into close proximity of the bilayer. **D**. By comparison, the two TM structure simulated in a model bilayer as for B. This conformation remained stable for the duration of the simulation (five repeat simulations, standard deviations in grey), and the region adjoining the inter-helical loop induced “snorkelling” of the lipid envelope.

Thus, we undertook molecular dynamics simulations to address whether M in the absence of potentially stabilising interactions with E dimers might prefer a double, or a single TMD conformation. Monomeric M protein was placed within model POPC bilayers (Figure 4c, d, S4), starting as either the cryo-EM derived conformer, or one where the two TMDs were straightened into a membrane - spanning helix containing a small unstructured region between helix two and three (Figure 4c, d, S5). Simulations of the two species within a POPC lipid bilayer revealed that the single helix rapidly began to fold back towards the membrane, which may have led to a hairpin-like structure during a longer simulation. Even if not the case, this form of the peptide was clearly unstable compared to the cryo-EM structure (Figure 4d). Thus, we consider it is likely that higher order M oligomers also comprise double-TMD protomers.

### Molecular dynamics simulations favour the formation of compact hexameric channel complexes

Observations from native PAGE and electron microscopy led us to construct hexameric, and heptameric models of M channels with helix 2 or 3 lining the channel lumen (Figure 5, S5, S6, annexe); helix 3 was predicted to display pore-lining properties based upon MEMSAT-SVM predictions (Figure S2b), so this class of models was the principal focus. Two different hexameric arrangements were modelled (Figure 5, S6-8): in the first arrangement, helix one was orientated almost perpendicular to the channel pore with helix three facing towards the pore ( termed “radial”, Figures 5e, S6b, S7). For the second model, the protein was rotated clockwise by 20 degrees to significantly increase the number of inter-protomer contacts within a more compact structure (termed “compact”, Figures 5a-e, S6, S8). The second conformation included interactions between the extra-membranous helix one. Anti-clockwise rotation was not possible due to stearic hindrance. We also generated radial and compact hexamer and heptamer models with helix two lining the pore, ensuring that other potential conformations were explored (see annexe). However, hexamer models where helix two lined the pore collapsed rapidly upon simulation, whereas heptamers lined by either helix showed an inability to close in the majority of simulations (Annexe). Moreover, radial hexamers lined by helix three either closed rapidly within the simulation (6, 40 ns) or not at all (>200 ns). In the two channels that closed, one of the subunits moves to the centre of the channel to occupy the pore (Annexe). This again was inconsistent with the formation of a stable channel complex.

**Figure 5.**
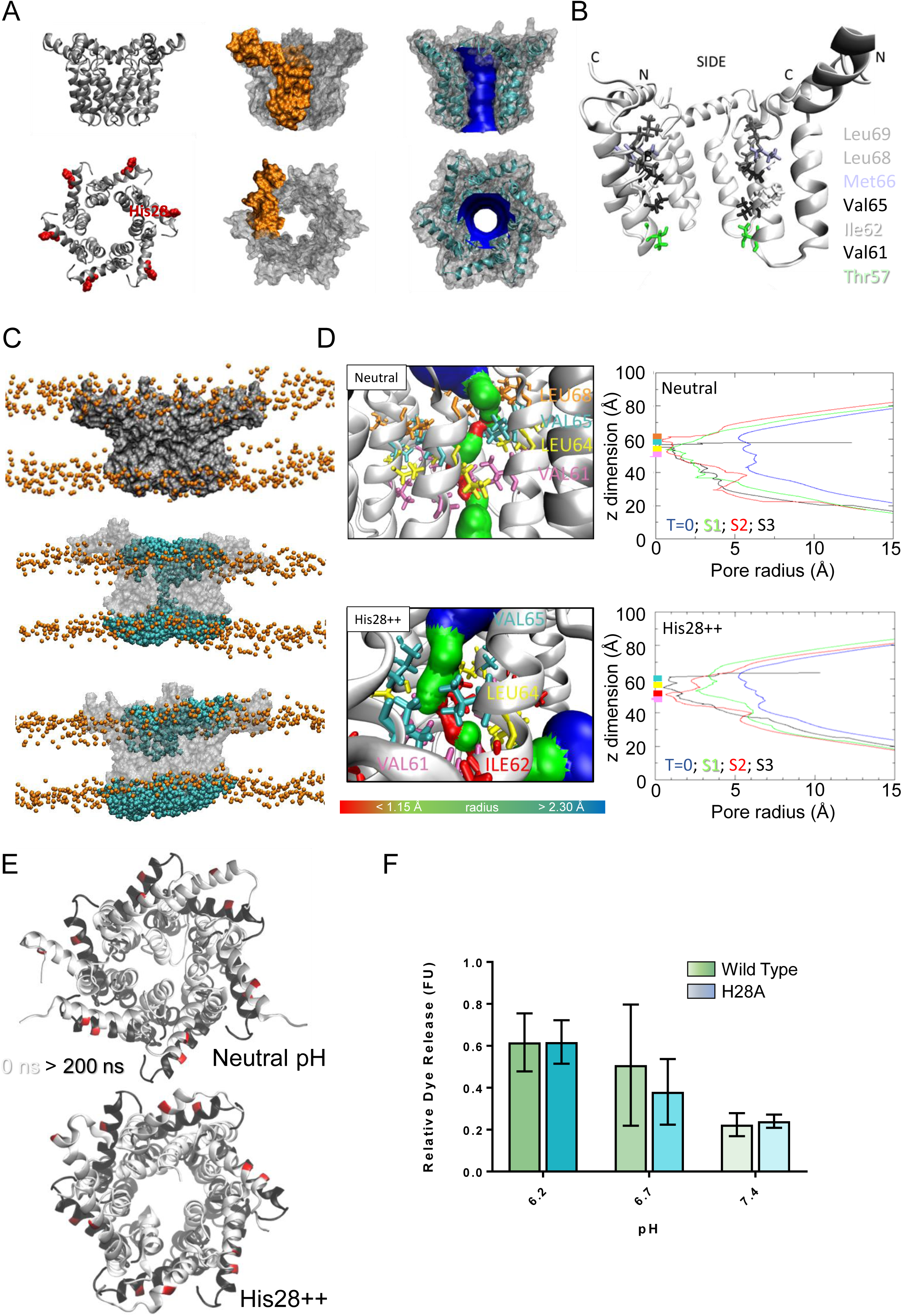
Simulated hexameric M channel complex in a compact formation, lined by helix 3. **A.** Assembly of protomers into compact hexamer model using Maestro. Left – ribbon diagram illustrating position of His28 (red); middle – space filling model showing individual protomer (gold); right – space filling model showing pore diameter using HOLE. **B.** Cutaway of channel viewed from the side showing pore-lining residues predicted for the compact hexamer from the side. **C.** Snapshots of compact hexamers in DOPC bilayers from a representative simulation showing the energy minimised system at t=0 (top), formation of a conductive water column during early times and eventual closing of the channels later on. **D**. HOLE profiles for lumenal aperture showing differences between starting configuration (blue line) and final structures over three separate 100 ns atomistic simulations (green, red, black) at both neutral and acidic external pH based upon dye release assays (mimicked here via dual His28 protonation). Top – neutral pH, bottom – protonated. Illustrations also reveal similar closing mechanisms in effect at both neutral and acidic pH, centred on Leu64. **E.** Neutral (left) versus protonated (right) compact hexameric complexes over a 200 ns simulation overlaying start (white) and endpoint (black) conformations. **F**. Repeat of pH experiment in Figure 2B comparing wild type (green) and His28Ala (blue) peptides. Results suggest that His28 is not in fact responsible for pH activated increases in membrane permeability.

Simulations supported that the compact hexameric model with helix three lining the lumen is likely to comprise the best physiological representation of a channel structure formed by M protein (Figure 5). The compact channel structure retained a lumen with a radius of 5.23 Å at its narrowest point, lined by Thr57, Val61, Ile62, Val65, Met66, Leu68 and Leu69 (Figure 5b). Upon 200 ns atomistic simulation within a model POPC bilayer, channels initially allowed the formation of a water column, but then closed at Leu64 and Leu68, despite the former not facing the lumen at the beginning of the simulation (Figure 5c, d, e). Channel closing first occurred at 70, 80 and 176 ns within three separate simulations, with channels remaining closed for the remainder of the simulation (Figure 5c, d). Thus, despite the starting position of this model possessing a relatively wide lumen com pared with the diameter of a water column (∼1.15 Å), compact M hexamer models lined by helix three displayed spontaneous closure, consistent with a reasonable physiological representation of a membrane channel. Finally, removal of helix one caused the collapse of stable pores within 50 ns in each of three simulations (Figure S6c), demonstrating its likely importance in the formation of a stable channel complex.

### Influence of protonation upon M channel complex

Data from dye release assays and a role early during the ZIKV life cycle were consistent with the hypothesis that acidic pH might activate M channels during endocytosis. The residue most likely to be protonatable at physiological pH within the E peptide sequence was His28, present within helix I. His28 was modified by dual protonation during simulations, mimicking the predicted effect of endosomal acidification (Figure 5d-e, Figure S6d-e). This change had little effect upon the starting structure compared to neutral conditions. His28 protonated channel complexes displayed minor, but appreciable differences in the time taken to close during three 200 ns atomistic simulations compared to the native model: 100, 130 and >200 ns (i.e. non-closure) (Figure 5d, e). The protonated channels closed involving similar residues to native complexes; Val61, Leu64, Val65 and Leu68 occluded the pore, along with Ile62 (Figure 5d, e). Moreover, a peptide containing a His28Ala mutation had no effect upon pH-activated M channel activity in dye release assays (Figure 5f). This supports that, whilst His28 protonation may play a minor role modulating native channel gating, it is more likely that other ionisable residues in the M protein sequence are either directly responsible for the acid enhancement of activity, or, alternatively, may be able to compensate experimentally for the loss of His28 protonation.

### Identification of two potential druggable sites within hexameric M complexes

We hypothesised that MD-tested compact hexamer models might be sufficiently accurate to both characterise and improve upon the inhibitory action of rimantadine. Whilst this drug exerted a genuine antiviral effect, its relatively low potency in cell culture was reminiscent of its promiscuous activity against other viroporins^44,47,48^. Nevertheless, the action of rimantadine both *in vitro* and *in vivo* implied that at least one physiologically relevant druggable binding site exists within the M channel complex.

Docking of rimantadine into the compact M hexamer model revealed an energetically favourable interaction with a lumenal binding site, close in proximity to the region in which channel closure occurred during simulations, termed “L1” (Figure 6a, Figure S9a-b). The adamantyl cage of rimantadine made predicted hydrophobic contacts with Val61, Ile62, Leu64, Val65 and Leu68, leaving the methyl group and amine solvent exposed, projecting into the channel lumen (Figure S9). The same site also underwent predicted interactions with other adamantyl-containing molecules, including amantadine, which adopted a similar predicted binding mode to rimantadine (Figure S9). However, both methyl- and acetyl-rimantadine reversed the orientation of the adamantyl cage such that the polar group interacted with lumenal residues (Figure S9).

**Figure 6.**
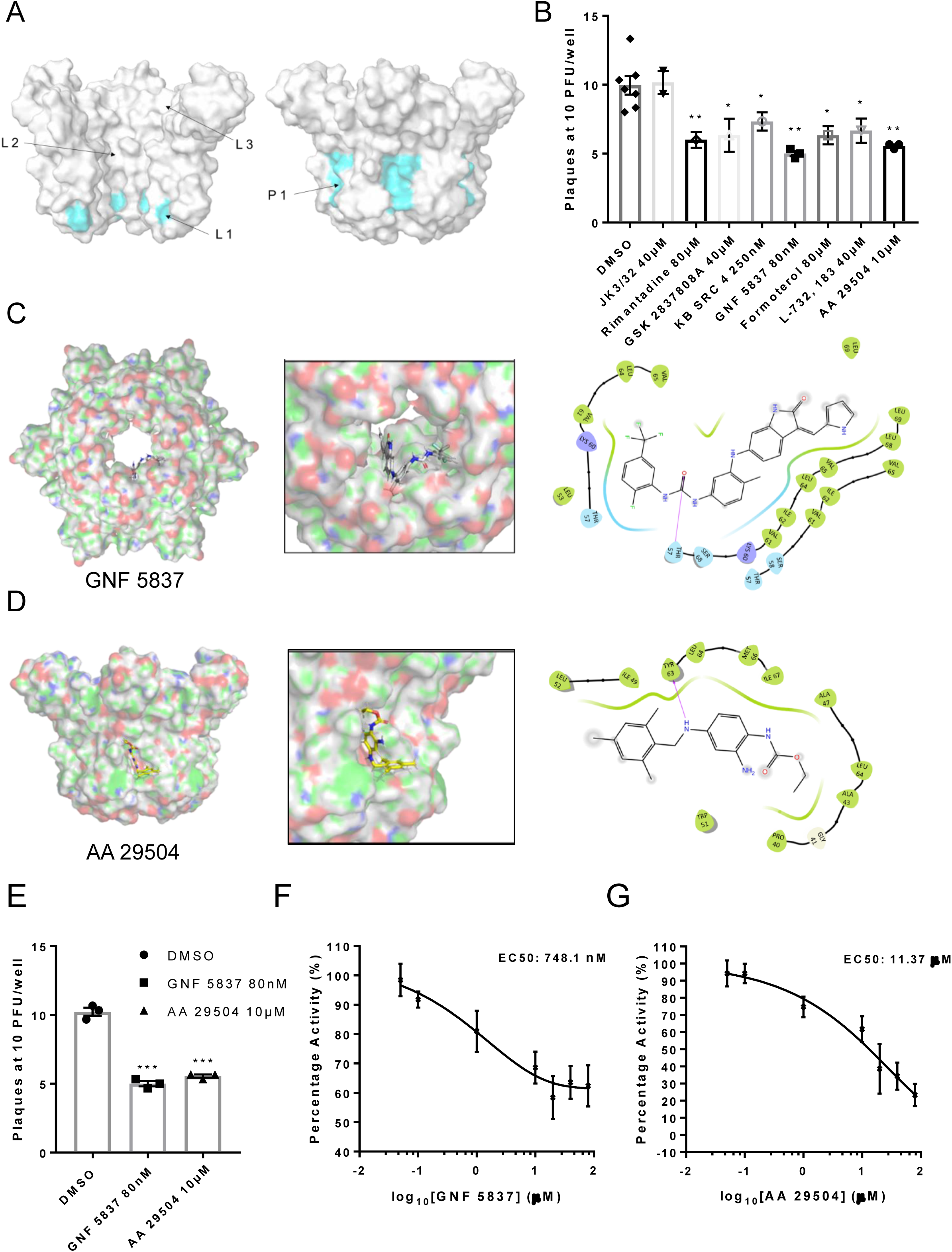
Inhibition of ZIKV replication by in silico enriched repurposed drugs. **A.** Compact M channel complex models were assessed using the SiteMap package within Maestro for cavities corresponding to high druggability scores. Three sites were found within the lumen (L1-L3), with L1 retaining the most favourable score. An additional site upon the membrane-exposed channel periphery was also evident (P1), yet would not score highly in SiteMap due to solvent exposure of compounds during simulated docking. Notably, both L1 and P1 reside within close proximity to the predicted gating region near Leu64. **B**. Compounds selected to bind the L1 and P1 sites *in silico* and screened *in vitro* (Fig S12) were tested for effects in ZIKV culture, using a BHK21 cell plaque reduction assay. Compounds were used at concentrations shown not to evoke cellular toxicity by MTT assays (Fig S13). Experiments are biological triplicates with error bars representing the standard error of the mean. **p≤0.01, *p≤0.05, Student T-test. **C.** Predicted binding pose of GNF 5837 at L1 site generated using Glide (Maestro, Schrodinger) with associated binding site interactions. **D**. As C, but for AA 29504. **E**. Example biological repeat with triplicate technical repeats for the two top compounds targeting the L1 and P1 sites, GNF 5837 (L1) and AA 29504 (P1). **F**. *In vitro* dye release assays showing 8-point log_2_-fold titration of GNF 5837 enabling calculation of 50% effective concentration (EC_50_). **G**. As F, for AA 29504.

Reassuringly, the lumenal rimantadine site (L1) was the most favourable in terms of druggability score for the compact hexamer model, assessed using the SiteMap programme (Maestro, Schrödinger). Two other potential lumenal sites were also identified with lower predicted druggability scores (L2, L3, Figure 6a). However, inspection of the M channel model identified an additional region upon the membrane-exposed periphery that could also undergo interactions with small molecules (P1, Figure 6a). Such binding sites are less likely to be identified computationally due to the high solvent exposure of potential ligands, yet such sites have been successfully targeted for other viroporins, including HCV p7^48,49^.

To explore the potential relevance of the P1 site, we exploited a defined series of compounds developed targeting a similar site upon the HCV p7 channel complex. The planar structure of these compounds is more compatible with the shape of the P1 site compared to L1, as supported by the predicted *in silico* modelling. Encouragingly, two of the seven compounds tested displayed inhibitory effects versus M in dye release assays, with JK3/42 being most active ( Figure S9c). Interestingly, adamantane derivatives with additional R groups were ineffective at targeting M activity (Figure S9d), potentially due to the limited size of the L1 cavity. Thus, we investigated the possibility that more than one distinct druggable binding site existed within M complexes.

### *In silico* screening enriches for improved repurposed M channel inhibitors

We investigated whether *in silico* screening could identify and enrich for compounds predicted to interact with L1 and/or P1, with a view to developing two distinct yet complementary inhibitor series. L1 was defined as the site comprising Val61, Leu64, Val65 and Leu68, and P1 by Tyr63, Leu64, Val65, Met66, Ile67 and Leu68. The two sites effectively represented the internal and external face of a single region within the channel complex, which is also involved during channel closing in simulations and centres upon Leu68. The convergence of these three regions was suggestive of their importance in being able to influence channel gating (Figures 5d, 5e, 6a).

To determine whether compounds with good drug-like properties could target the L1 and P1 sites, we employed a screening library comprised of FDA-approved, generic, and other compounds with proven biological activity. *In silico* screening of 1280 compounds (TocrisScreen https://www.tocris.com/product-type/tocriscreen-compound-libraries) was undertaken using Glide (Schrödinger) and a rank-order list generated for each site following attrition and removal of compounds common to both sites. A short-list of 50 compounds targeting each site was generated for subsequent validation *in vitro* using dye release assays (Table 1, 2). Note, predicted docking scores are not directly comparable between P1 and L1 owing to binding penalties incurred at the P1 site discussed above.

**Table 1.** Top 50 lumenally targeted compounds from the TOCRIS screen library, ranked by glide score.

| "Rank" | Compound Name | Tocris ID | Glide gscore |
| --- | --- | --- | --- |
| 1 | FR 139317 | 1210 | -10.227 |
| 2 | ZCL 278 | 4794 | -9.294 |
| 3 | Elinogrel | 5316 | -8.829 |
| 4 | Taxol | 1097 | -8.711 |
| 5 | UK 356618 | 4187 | -8.401 |
| 7 | TC-1 15 | 4527 | -8.216 |
| 8 | AS 2034178 | 5035 | -8.032 |
| 9 | CP 775146 | 4190 | -7.963 |
| 10 | GW 6471 | 4618 | -9.092 |
| 11 | Pravastatin sodium salt | 2318 | -7.825 |
| 12 | Fluvastatin sodium | 3309 | -7.717 |
| 13 | TC NTR1 17 | 5087 | -7.683 |
| 14 | VER 155008 | 3803 | -7.686 |
| 15 | AMG PERK 44 | 5517 | -7.618 |
| 16 | Glibenclamide | 0911 | -7.537 |
| 17 | Argatroban | 1637 | -7.454 |
| 18 | KB SRC 4 | 4660 | -7.444 |
| 19 | GBR 12909 dihydrochloride | 0421 | -7.965 |
| 20 | (±)-NBI 74330 | 4528 | -7.366 |
| 21 | CU CPT 4a | 4883 | -7.331 |
| 22 | A 887826 | 4249 | -7.315 |
| 23 | SR 2640 hydrochloride | 1804 | -7.296 |
| 24 | NSC 74859 | 4655 | -7.258 |
| 25 | RWJ 67657 | 2999 | -7.217 |
| 26 | Lu AA 47070 | 4783 | -7.659 |
| 27 | Edaglitazone | 4784 | -7.176 |
| 29 | GSK 1562590 hydrochloride | 5110 | -7.11 |
| 30 | Flurizan | 4495 | -7.098 |
| 31 | GW 9508 | 2649 | -7.096 |
| 32 | GSK 269962 | 4009 | -7.051 |
| 33 | AC 5216 | 5281 | -6.986 |
| 34 | DBZ | 4489 | -6.973 |
| 35 | PF 04418948 | 4818 | -6.957 |
| 36 | GSK 2837808A | 5189 | -6.942 |
| 37 | Sal 003 | 3657 | -6.935 |
| 38 | PD 173212 | 3552 | -7.029 |
| 39 | NTRC 824 | 5438 | -9.012 |
| 40 | ONO AE3 208 | 3565 | -6.889 |
| 41 | RS 17053 hydrochloride | 0985 | -6.88 |
| 42 | Pitavastatin calcium | 4942 | -6.864 |
| 43 | L-161,982 | 2514 | -6.858 |
| 44 | AMN 082 dihydrochloride | 2385 | -6.871 |
| 45 | TC-N 1752 | 4435 | -6.815 |
| 46 | PF 431396 | 4278 | -6.799 |
| 47 | GNF 5837 | 4559 | -6.754 |
| 48 | KS 176 | 4169 | -6.731 |
| 49 | Sarpogrelate hydrochloride | 3739 | -6.769 |
| 50 | GKA 50 | 5133 | -6.689 |
| N1 | Mifepristone | 1479 | 0.286 |

**Table 2.** Top 50 peripherally targeted compounds from the TOCRIS screen library, ranked by glide score.

| "Rank" | Compound | Cat No | Glide gscore |
| --- | --- | --- | --- |
| 1 | RWJ 21757 | 2719 | -6.475 |
| 2 | Ferrostatin 1 | 5180 | -6.309 |
| 3 | AA 29504 | 3972 | -6.31 |
| 4 | L-732,138 | 0868 | -6.167 |
| 5 | API-2 | 2151 | -6.158 |
| 6 | 5-BDBD | 3579 | -6.13 |
| 7 | LY 225910 | 1018 | -6.082 |
| 8 | Formoterol hemifumarate | 1448 | -6.065 |
| 9 | TC-S 7006 | 5240 | -5.979 |
| 10 | TCS 2210 | 3877 | -6.005 |
| 11 | Sumatriptan succinate | 3586 | -5.952 |
| 12 | MRS 3777 hemioxalate | 2403 | -6.964 |
| 13 | Thiamet G | 4390 | -6.079 |
| 14 | Abacavir hemisulfate | 4148 | -5.894 |
| 15 | DDR1-IN-1 | 5077 | -5.99 |
| 16 | Fexofenadine hydrochloride | 2429 | -5.859 |
| 17 | 6-Chloromelatonin | 0443 | -5.844 |
| 18 | GSK 0660 | 3433 | -5.864 |
| 19 | PCA 4248 | 0571 | -5.783 |
| 20 | Axitinib | 4350 | -5.765 |
| 21 | DSR 6434 | 4809 | -5.971 |
| 22 | Necrostatin-1 | 2324 | -5.756 |
| 23 | Trifluorothymidine | 4460 | -5.838 |
| 24 | Cilindipine | 2629 | -5.729 |
| 25 | Efondipine hydrochloride<br>monoethanolate | 3733 | -5.693 |
| 26 | ITE | 1803 | -5.69 |
| 27 | L-165,041 | 1856 | -5.689 |
| 28 | EB 47 | 4140 | -6.033 |
| 29 | GSK 2830371 | 5140 | -5.657 |
| 30 | AZD 1480 | 5617 | -5.738 |
| 32 | Amlodipine besylate | 2571 | -5.638 |
| 33 | Melatonin | 3550 | -5.628 |
| 34 | Fludarabine | 3495 | -5.626 |
| 35 | PF 06447475 | 5716 | -5.609 |
| 36 | SU 6668 | 3335 | -5.608 |
| 37 | AZD 5438 | 3968 | -5.836 |
| 38 | SU 11274 | 4101 | -6.145 |
| 39 | FPL 64176 | 1403 | -5.543 |
| 40 | Sunitinib malate | 3768 | -5.543 |
| 41 | YK 4-279 | 4067 | -5.535 |
| 42 | Ralfinamide mesylate | 4029 | -5.678 |
| 43 | ML 298 hydrochloride | 4895 | -5.504 |
| 44 | FH 1 | 5254 | -5.485 |
| 45 | PLX 647 dihydrochloride | 5102 | -6.23 |
| 46 | GPI 688 | 3967 | -5.438 |
| 47 | CP 94253 hydrochloride | 1317 | -5.441 |
| 48 | CGP 57380 | 2731 | -5.44 |
| 49 | BW 723C86 hydrochloride | 1059 | -5.433 |
| 50 | LY 364947 | 2718 | -5.84 |

Dye release assay screens (96-well format) were conducted using compounds at a concentration of one micromolar, with positive hits defined as compounds exerting a 50 +/- 5% decrease in M activity (Figure S10). Twenty-four hits were identified from the L1-targeted short-list, and fifteen for the P1 site.

We selected three compounds from each list, based upon both potency and commercial availability, for validation in plaque-reduction assays. As the components of the repurposing library predominantly target cellular factors, MTT assays were used to determine non-toxic maximal concentrations for each compound (Figure S11). Testing at these concentrations revealed that each of the compounds exerted a statistically significant reduction upon ZIKV infectivity, with the most effective showing sub-micromolar potency (Figure 6b), whereas the inactive JK3/32 compound showed no detectable antiviral effects. Two L1-selected compounds, KB SRC 4 and GNF 5837 displayed antiviral efficacy at 250 and 80 nM, respectively, whereas the most effective P1 -targeted compound, AA 29504, was active at 10 μM. Examination of the predicted binding pose for these compounds at L1/P1 revealed that both molecules underwent additional interactions with regions adjacent to the core binding sites (Figure 6c-d). GNF 5837, an inhibitor of the tropomyosin receptor kinase (TRK) family, was predicted to make extended contacts within the lumen, including with two of the predicted rimantadine binding sites, stabilised by H-bonding between a carbonyl and Thr57 (Figure 6c). These leading hit compounds were re-tested in a second round of plaque reduction assays, confirming efficacy (Figure 6e) and displayed *in vitro* EC_50_ values at similar orders of magnitude as their observed potency in cell culture, namely sub-micromolar for GNF 5837, and approximately 10 micromolar for AA 29504 (Figure 6 f, g).

The considerable enrichment for potency, compared to rimantadine, seen amongst compounds selected *by in silico* screening supports the validity and utility of structure-guided models in their potential use as templates for rational drug development.

### M channel inhibitors reduce the specific infectivity of ZIKV virions

It was important to establish whether M-targeted compounds might exert off-target effects upon virion stability, or other aspects of virus entry. Frustratingly, the role of (pr)M during virion egress precludes the efficient generation of *Flavivirus* envelope pseudotype systems in their absence, a useful tool in this regard. As an alternative, compound treated ZIKV particles were compared to solvent controls following ultracentrifugation and separation using a continuous iodixinol gradient. This not only allows characterisation of specific infectious species, but also sequesters small molecules at the top, lower density range of the gradient, preventing incidental exposure of target cells when inoculating with virus-containing fractions, and so minimising potential cell off-target effects.

Interestingly, ZIKV infectivity profiles contained two peaks of infectivity when samples were diluted 1:10 onto target cells: a low-density peak at ∼1.11 g/mL, and a high-density peak at 1.13-1.14 g/mL. However, detection of infectivity within the low-density peak diminished upon dilution at 1:100 or 1:1000, and this was not the case for the high-density peak (Figure S14). Interestingly, the distribution of the ZIKV E protein was more abundant within lower density fractions compared to the main infectivity peak at higher density (Figure 7a). Drug treatment specifically reduced infectivity (measured at 1:10 supernatant dilution) within the high-density infectious fraction compared to controls; the lower density fractions were unaffected (Figure 7a-b). Importantly, as well as a global loss of infectivity, specific infectivity within the high-density fraction normalised by RT-qPCR for ZIKV RNA (E gene target) was also significantly reduced (Figure 7c), meaning that intact virions were rendered less infectious following exposure to inhibitors rather than being physically disrupted via non-specific “virolysis”. Moreover, based upon previous studies, the concentration of inhibitors added to target cells from high density fractions would have been negligible, excluding artefactual effects upon cell entry.

**Figure 7.**
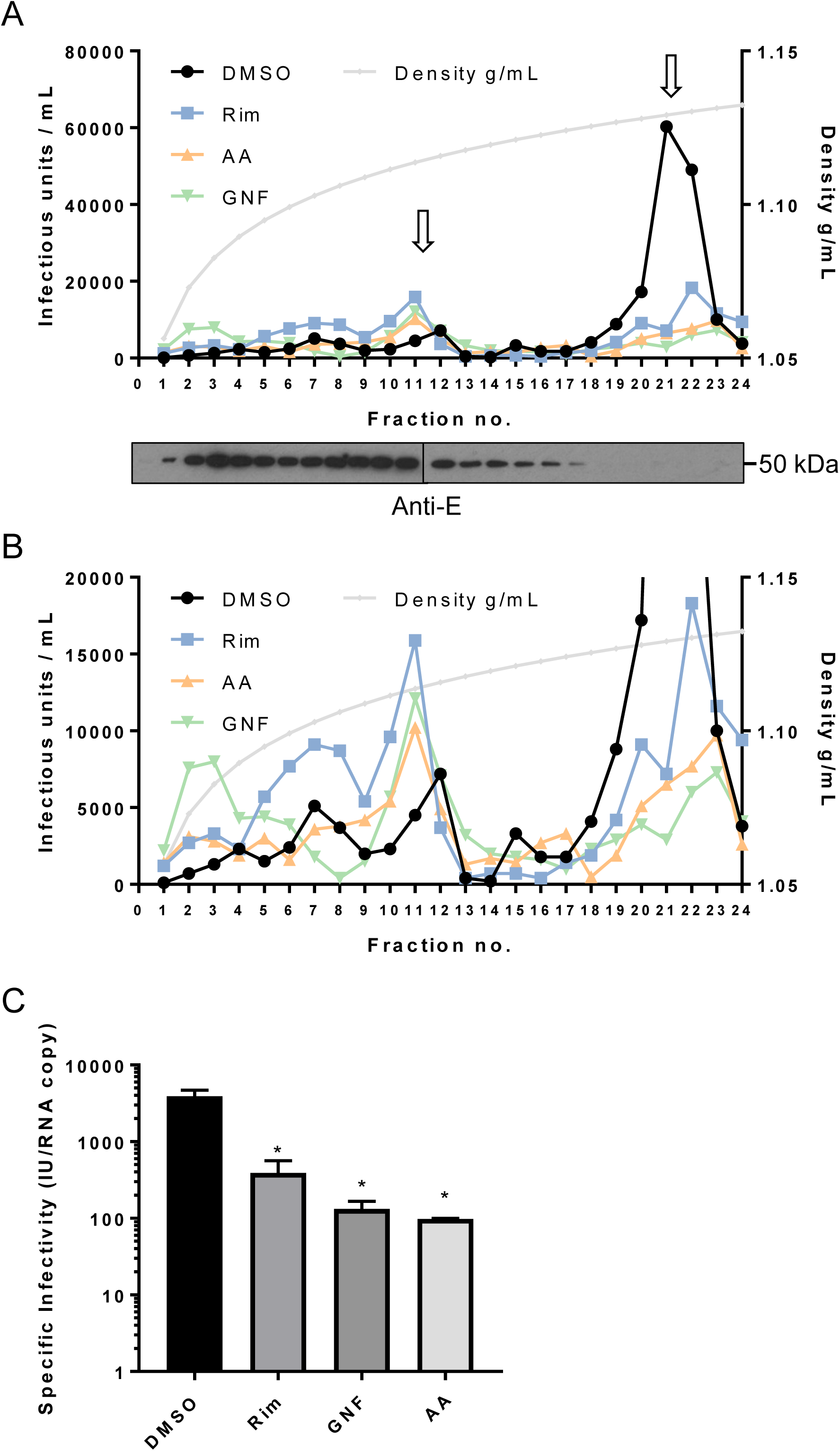
M channel inhibitors lack virus-lytic or cell-targeted effects. To exclude the possibility that early effects upon ZIKV infection were due to a directly damaging effect of compounds upon virion integrity, particles were concentrated and purified from Vero cell supernatants by PEG-precipitation and ultracentrifugation through a sucrose cushion. Virions were then dosed with inhibitors prior to ultracentrifugation through a continuous iodixinol gradient. This not only purifies virions according to buoyant density, but separates them from small molecules, precluding off-target effects upon cells during ensuing infection (see methods for details). **A.** Infectivity profile from a representative gradient experiment where purified virions were treated with rimantadine (40 μM), GNF 5837 (1 μM), or AA 29504 (10 μM) prior to ultracentrifugation (or, DMSO at equivalent concentrations). 10 μl from each fraction was then used to infect naïve Vero cells in a 96-well plate, representing a 10-fold dilution. Infected cells were detected by immunofluorescence and counted as described elsewhere. A further 15 μl was analysed by western blotting for the (E)nv glycoprotein (lower panel). **B.** Zoomed in view of gradient in A, with Y-axis capped at 20, 000 units to delineate infectivity profiles. **C.** Specific infectivity of virions in the presence/absence of inhibitors based upon q-RTPCR normalisation confirms the reduction of infectivity rather than disrupted virions.

### Rimantadine prevents ZIKV viraemia *in vivo*

We assessed rimantadine antiviral effects *in vivo* using a preclinical ZIKV infection model^55^ comprising C57BL/6 mice and transient blockade of the interferon alpha receptor type 1 (IFNAR1). This allows the establishment of robust infection within an immunocompetent host and does not lead to the severe sequelae seen within immunodeficient systems. Moreover, the model incorporates improved physiological relevance via concomitant biting at the inoculation site by female *Aedes aegypti* mosquitos, which enhances the efficiency of infection (Figure 8a).

**Figure 8.**
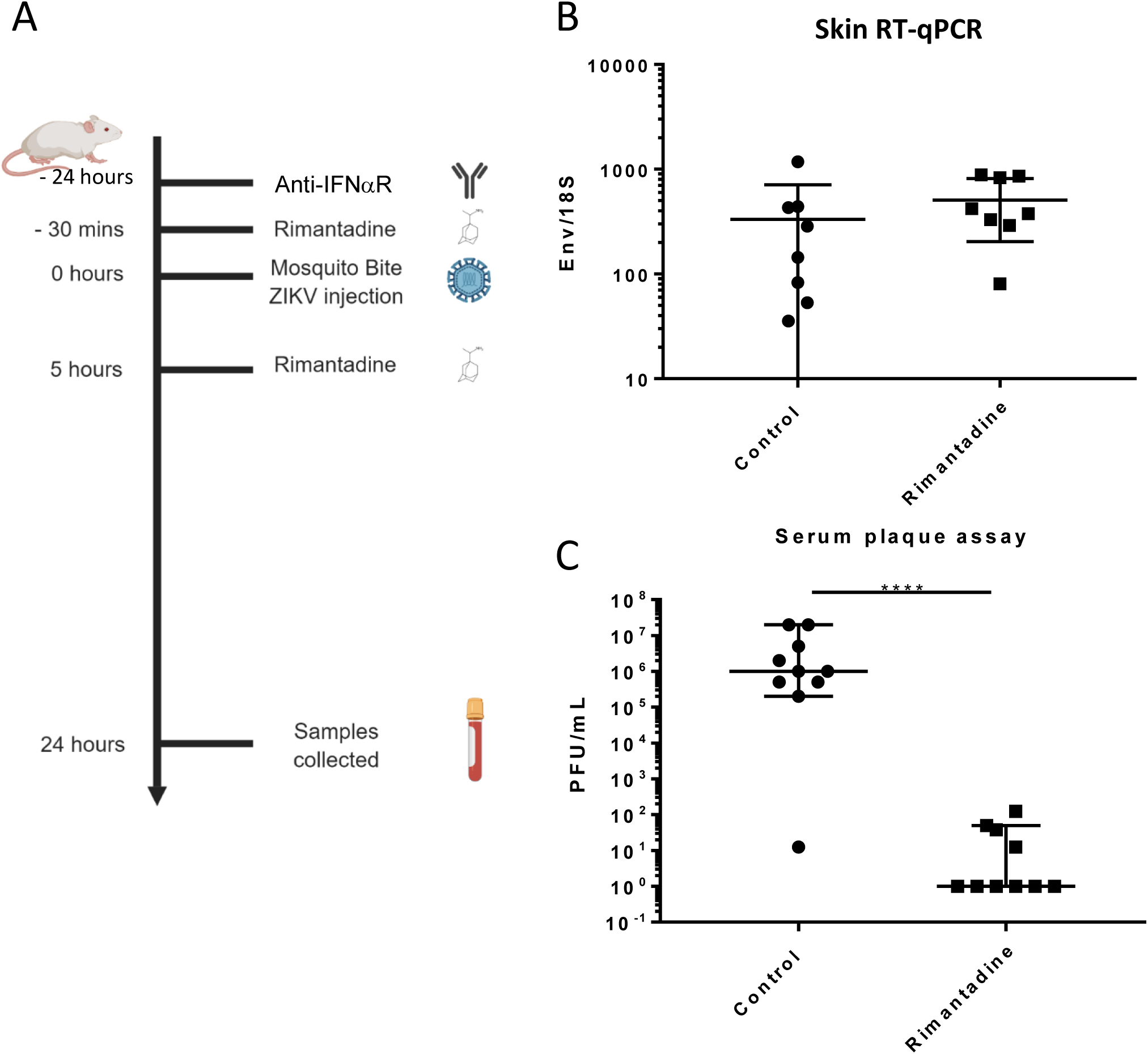
Anti-ZIKV activity in preclinical models. **A.** Schematic of preclinical experiments involving transient IFNAR blockade, augmentation of ZIKV infection through mosquito bites and treatment with rimantadine. **B.** RT-qPCR for ZIKV E relative to 18S RNA from skin tissue at injection site derived from one of two representative experiments. **C.** Infectious titre of serum derived from rimantadine treated or control mice, determined by plaque assay in BHK21 cells. ****p≤0.0001, Student T -test.

Ten C57BL/6 mice received a sub-cutaneous dose of 20 mg/kg rimantadine (or carrier control), 30 min prior to ZIKV infection (10^3^ plaque forming units, pfu) by injection into the dorsal side of one hind foot, immediately following exposure to up to five mosquito bites. A second bolus of rimantadine was administered 5 hr post-infection. Animals were sacrificed and processed 24 hr post-infection, harvesting tissue from the inoculation site as well as blood via cardiac puncture. RT -qPCR analysis confirmed equivalent copies of viral RNA present within tissues at the inoculation site ( Figure 8b), whereas viraemia was dramatically reduced within rimantadine treated animals, measured by BHK cell plaque assay (Figure 8c). Thus, rimantadine exerted an antiviral effect *in vivo* that prevented dissemination of the virus, consistent with cell culture observations, and supporting the druggability of M *in vivo*.

## Discussion

This work supports that the ZIKV M protein mediates membrane permeability *in vitro* through the formation of oligomeric structures, and that small molecules targeting this activity also decrease the infectivity of ZIKV at an early stage of the life cycle in culture. This is reminiscent of the IAV M2 protein, which responds to endosomal pH to allow protonation of the virion core, expediting virion uncoating^56^; we have recently demonstrated that HCV p7 plays a similar role ^49^. Moreover, like M2, the activity of M is sensitive to both prototypic small molecules and rationally targeted ligands, preventing the spread of infection both in cell culture and *in vivo*. Thus, M channels represent a potential new target for antiviral therapies that could both limit disease severity and engender prophylactic use to disrupt transmission.

M resides as E-associated dimers within the mature virion^25^, yet for M channels to form requires the assembly of higher order structures in the membrane to generate an intact aqueous pore. However, the rearrangement and formation of trimeric E complexes during acid-induced fusion provides an opportunity for M to also adopt alternative conformations. Density corresponding to M is absent from cryo-EM reconstructions of acidified mature *Flavivirus* virions, which are themselves inherently unstable and require the binding of E-specific FAb fragments to preserve their architecture^26^. It is possible that asymmetric formation of relatively few channel complexes leads to a loss of signal upon application of icosahedral symmetry, or due to the overwhelming inherent symmetry of the particle. It would be of interest to apply symmetry expansion or focused classification techniques to acidified virions to investigate this possibility.

DENV M protein was previously proposed to function as a viroporin based primarily upon its size and hydrophobicity, yet conclusive evidence for a defined role within *Flavivirus* life cycles has been lacking. C-terminal M derived peptides from DENV type 1 were shown to form cation -selective channels in suspended bilayers^32^, with sensitivity to amantadine (10 μM) and higher concentrations of hexamethylene amiloride (HMA, 100 μM). However, a subsequent study using a series of pr-M protein expression constructs with additional N-terminal signal peptides and myc epitope tags failed to demonstrate pH activated channel activity in *Xenopus laevis* oocytes^33^. Nevertheless, in addition to the difficulties interpreting negative data, it was unclear whether proteins derived from t hese constructs underwent authentic proteolytic processing to enable trafficking of mature M protein to the oocyte membrane. Older studies showed that DENV replication in human peripheral blood leukocytes and Rhesus macaque kidney epithelial cells (LLC-MK2) was suppressed by amantadine and rimantadine, with maximal effect when drugs were administered concomitant with infection ^57^, reminiscent of findings herein.

Consistent with the formation an aqueous pore across the membrane, ZIKV M peptides migrated as higher order structures when reconstituted in the membrane-mimetic detergent, DH_6_PC. These corresponded to potential hexamers or heptamers, albeit estimated by native PAGE, and were confirmed to adopt a channel-like architecture using TEM. However, heterogeneity prevented determination of precise stoichiometry by rotational averaging. Nevertheless, M complexes mediated membrane bilayer permeability using an indirect ion channel assay, based upon fluorescent dye release from liposomes. Dose-dependent dye release was not only increased by acidic pH, but was also sensitive to rimantadine. Importantly, rimantadine sensitivity supports that dye release was mediated by interactions with a folded protein complex rather than via aggregates disrupting membrane integrity, and increased activity at acidic pH is consistent with activation within acidifying endosomes during virus entry. Other pH-gated channels including IAV M2, HCV p7 (genotype and polymorphism-dependent) and HPV E5, are similarly activated within the same assay _system44,48,49,52._

Surprisingly, mutation of His28 to Ala, which is conserved between ZIKV and DENV and represents the most obviously solvent-accessible, ionisable residue present within M protein peptides, did not affect the acid-activation of M channel activity *in vitro*. This contrasted the apparent influence of this residue during MD simulations, whereby dual protonation of His28 extended average channel opening times, albeit to a relatively modest extent. Notably, multiple residues in addition to His28 could theoretically become protonated, albeit most being less likely at physiological pH. It may be that these are able to compensate for the loss of His28 *in vitro*, or, conversely, that another residue or residues are in fact responsible, and this was merely emulated by His28 protonation during simulations.

Rimantadine was able to interrupt ZIKV infection of Vero cells, causing a dose-dependent reduction in the number of cells infected and ensuing virus protein expression. The EC _50_ for rimantadine was between 10-20 μM, which is comparable to drug potency versus susceptible HCV strains ^47^. Rimantadine was most effective when added prior to and simultaneously with infectious innoculae, suggesting that pre-loading of endocytic vesicles with the drug may expedite the inhibition of virion-resident channels. Interestingly, adding rimantadine both during and after infection was consistently less effective than adding it during the infection alone. We are uncertain why this should be the case, yet we speculate that in addition to inhibitory properties, rimantadine may also exert a drug rescue effect upon cells during multi-cycle replication, lessening the cytopathic effects of ZIKV infection and thereby artificially inflating the number of surviving cells remaining at the end of the assay.

However, rimantadine also potently suppressed ZIKV replication in a preclinical model, supporting that M channels constitute a physiologically relevant drug target. Rimantadine treatment prevented viraemia, consistent with an effect upon virus spread linked to drug effects observed in culture. To date only M2, and more recently SARS-CoV2 E, are the only examples where targeting a viroporin has been shown to exert preclinical antiviral efficacy. Thus, our findings strongly support that M channels are also physiologically relevant drug targets.

In lieu of structural information for channel complexes, we employed molecular modelling and molecular dynamics simulations to understand how channels might be formed from either dimeric or monomeric M protein. Importantly, monomer simulations indicated that a single *trans*-membrane spanning topology was unstable resulting in the C-terminus folding back towards the membrane. Hence, we considered this to support that dual-*trans*-membrane topology protomers comprise the more energetically favourable conformation for mature M, as present within the mature virion^25^. This was despite some prediction packages including TMHMM, TOPCONS and Phobius, supporting that M formed single-pass protomers. By contrast, MEMSAT-SVM not only predicted dual-pass conformations, but also predicted that helix three would most likely be pore lining, despite the majority of lumenal residues being hydrophobic in nature.

Compact hexameric models for M channel complexes comprised of dual-spanning protomers displayed consistent channel-like characteristics compared to an array of alternative conformations and oligomeric states. Models showed increased propensity to remain open with His28 protonation as discussed above, and helix one contributed significantly to overall channel stability. Channel closure occurred in proximity to Leu64, closing off the water column. One limitation of these models is that the amino-terminal seventeen amino acids were omitted as this region is predicted to be unstructured making it unlikely to contribute to the overall channel complex architecture. However, this region contains multiple ionisable residues, including His7, which is highly conserved in ZIKV, DENV, WNV and YFV.

Whilst a blunt tool in terms of potency, rimantadine (or other prototypic blockers) binding can signify druggable binding sites within viroporin complexes^46,48,49^. Rimantadine was predicted to bind within a lumenal cavity proximal to Leu64, repeated along the six-fold radial symmetry of the channel model. In addition, the external membrane exposed face of the same part of the channel model also contained a cavity predicted to be conducive to binding small molecules. The presence of two binding sites for small molecules could enable M-targeted drug combinations that might minimise the chance of resistant variants being selected, potentially also achieving synergistic antiviral effects. Thus, a series of chemically distinct molecules was identified using *in silico* HTS targeting each binding site, derived from a drug repurposing library. Hits were shortlisted for predicted binding accounting for hydrophobic exposure induced penalties at the peripheral site, chemical properties (Lipinski), and those predicted to bind both sites were removed from both lists. Screening of short-listed compounds using dye release assays identified multiple hits targeting both sites, several of which were corroborated by testing for antiviral effects in ZIKV infectious assays. (Table 3).

**Table 3.** Properties of hits taken forward into ZIKV cell culture screens, including effects versus canonical targets.

| Drug | M binding site (EC/IC<sub>50</sub>) | Canonical Target (Ki/EC/IC<sub>50</sub>) | MTT effect? |
| --- | --- | --- | --- |
| Rimantadine | Lumen (80 μM) | IAV M2 (variable) | No |
| GNF 5837 | Lumen (~80 nM) | Tropomyosin receptor kinases (TrK) A, B and C (10 nM) | No |
| KB SRC 4 | Lumen (~250 nM) | c-Src (Ki= 44 nM) | Yes, 500 nM |
| GSK 2837808A | Lumen (~40 μM) | Lactate dehydrogenase A and B (LDHA/B), 2.6 / 43 nM | No |
| Formoterol | Periphery (~80 μM) | $\beta_2$-adrenergic receptor agonist (pM range) | Yes, 100 μM |
| L-732, 183 | Periphery (~40 μM) | Tachykinin NK<sub>1</sub> receptor antagonist (IC<sub>50</sub> = 2.3 nM) | No |
| AA 29504 | Periphery (~10 μM) | GABA<sub>A</sub> receptor agonist (IC<sub>50</sub> = 9-13 μM) | Marginal, 20 μM |

**Table 4.**
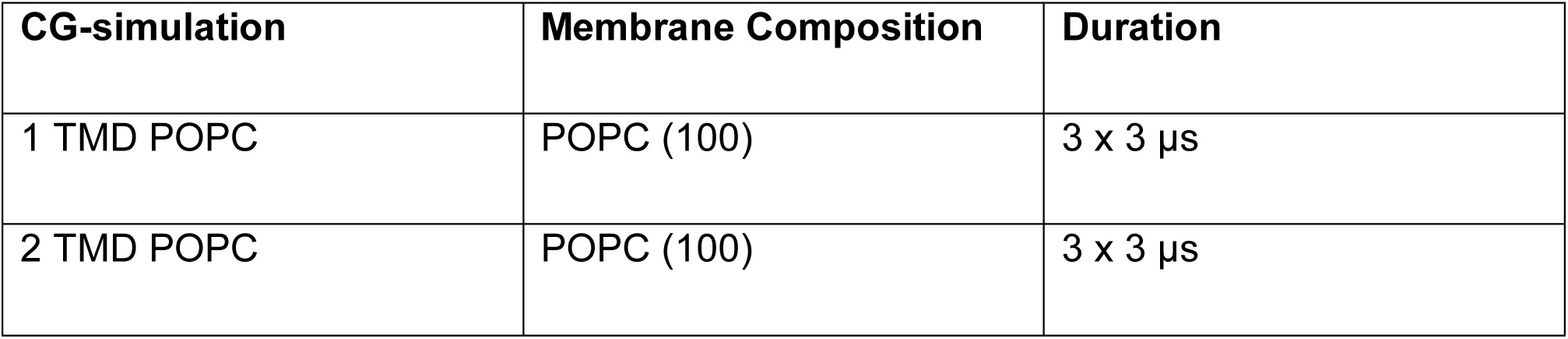
List of Monomer Course-Grain Simulations (martini 2.2 force field)

**Table 5.** List of all-atom simulations (CHARMM36 force field)

| AA-simulation | Membrane Composition | Duration |
| --- | --- | --- |
| **Hexamer** (Lumenal helix; compact vs radial; pH) |  |  |
| H2 Radial | POPC (100) | 2 x 200 ns |
| H2 Radial protonated | POPC (100) | 2 x 200 ns |
| H2 Compact | POPC (100) | 2 x 200 ns |
| H2 Compact Protonated | POPC (100) | 2 x 200 ns |
| H3 Radial | POPC (100) | 3 x 200 ns |
| H3 Radial protonated | POPC (100) | 3 x 200 ns |
| H3 Compact | POPC (100) | 3 x 200 ns |
| H3 Compact Protonated | POPC (100) | 3 x 200 ns |
| **Heptamer** (Luminal helix; compact vs radial; pH) |  |  |
| H2 Radial | POPC (100) | 2 x 200 ns |
| H2 Radial protonated | POPC (100) | 2 x 200 ns |
| H2 Compact | POPC (100) | 2 x 200 ns |
| H2 Compact Protonated | POPC (100) | 2 x 200 ns |
| H3 Radial | POPC (100) | 2 x 200 ns |
| H3 Radial protonated | POPC (100) | 2 x 200 ns |
| H3 Compact | POPC (100) | 2 x 200 ns |
| H3 Compact Protonated | POPC (100) | 2 x 200 ns |

Despite the correlation of anti-M potencies *in vitro* with antiviral effects, it cannot yet be excluded that antiviral efficacy may result from indirect mechanisms linked to the canonical cellular targets of these repurposed ligands. By far the most potent hit against M was GNF 5837, which was predicted to bind the L1 site. GNF 5837 is a potent inhibitor of Tropomyosin receptor kinases (TrK) A, B and C, high affinity receptors for nerve growth factor (NGF) that regulate neuronal survival and differentiation^58^. GNF 5837 has an IC_50_ of ∼ 10 nM against its cognate targets, yet only achieved an antiviral ∼ EC_50_ at 80 nM in BHK cell plaque reduction assays. Importantly, TrK expression is primarily restricted to the CNS and the thyroid, with only very low expression in kidney tissues according to the human protein atlas. Moreover, we could find no reference to TrK expression in either Vero, or BHK cells. Other lumenal hits comprised the c-Src inhibitor, KB SRC 4^59^ (250 nM), and GSK 2837808A^60^ (40 μM), an inhibitor of lactate dehydrogenase A and B (LDHA/B). KB SRC 4 has an *in vitro* Ki of 44 nM versus c-Src (higher against other Src-related kinases), whilst GSK 2837808A has an IC_50_ of 2.6 and 43 nM against LDHA/B, respectively; the former impeded cell metabolic activity at 500 nM, whilst the latter had no effect. We therefore surmise that GSK 2837808A is likely affecting ZIKV replication specifically, due to the considerable increase in effective concentration required, whereas it is conceivable that both GNF 5837 and KB SRC 4 may exert indirect, albeit potent antiviral effects.

The peripheral hits tested in culture comprised Formoterol hemifumarate (80 μM), L-732 183 (40 μM) and AA 29504 (10 μM). Only high concentrations of Formoterol impeded metabolic activity (100 μM). Formoterol is a long-acting β_2_-adrenergic receptor agonist (IC_50_ in pM range), used as a bronchodilator in the treatment of chronic obstructive pulmonary disease (COPD) ^61^. L-732 183 is a potent competitive tachykinin NK_1_ receptor antagonist (IC_50_ = 2.3 nM), the receptor for substance P^62^, whilst AA 29504^63^ is a positive allosteric modulator of GABA_A_ receptors (IC_50_ = 9-13 μM). The considerable difference in potency for both Formoterol and L-732 183 targeting their canonical targets compared to antiviral effects implies that the two are independent phenomena. However, this is less clear for differential potency of AA 29504, although, much like the case for GNF 5837, GABA_A_ receptors, the target of AA 29504, are not expressed in the kidney.

Further evidence that repurposed compounds and rimantadine exerted specific antiviral effects resulted from density gradient separation of ZIKV virions from unbound compounds , as described previously*(49)*. This ensures that target cells are highly unlikely to be exposed to meaningful concentrations of each compound, minimising one possible source of off -target effects. Moreover, virolytic mechanisms were found to be similarly unlikely to explain compound effects as the specific infectivity of virions was diminished by drug treatment, normalised by ZIKV RNA content. Interestingly, compounds specifically reduced infectivity in the major high-density peak, but not in a lower density region of the gradient where lower levels of infectivity were susceptible to relatively low serial dilutions; this was not the case for the high density peak. Interestingly, the only published study we identified relating to multiple particle types in *Flaviviruses* was a study assessing the effects of pr-M cleavage site mutations in the morphogenesis and secretion of tick-borne encephalitis virus (TBEV) subviral particles (SVPs), generated via pr-M-E expression alone^64^. Preventing pr-M cleavage by furin increased the proportion of larger, faster sedimenting SVPs corresponding in size to virions, and which harboured altered E protein glycosylation. Whilst the link between altered particle species, M processing and, potentially, M channel function is intriguing, it currently remains unclear whether these aspects are linked mechanistically.

The compounds identified herein represent only the first steps towards the design of bespoke M inhibitors, which could be enhanced through the exploration of chemical space around potent repurposed ligands; ideally, this will enable the continued emphasis on developing small molecules with superior qualities compared to rimantadine, or other prototypic viroporin inhibitors. Nevertheless, given its activity *in vivo* and in cell culture, there may be value in considering the immediate clinical repurposing of rimantadine to combat severe ZIKV infection and/or *trans*-placental transmission via early-stage clinical trials. The epidemic in South America and ensuing worldwide spread illustrated the speed with which more pathogenic ZIKV can be transmitted, and outbreaks continue in Africa and the Indian subcontinent. ZIKV is also unique amongst *Flaviviruses* as it can also spread from person to person via sexual contact ^65^, as well as tragically infecting the unborn foetus causing microcephaly^66^.

It is possible that emerging ZIKV replicase-targeted drugs such as Galidesivir will not be safe to use during pregnancy due to potential teratogenic effects, although recent studies in Rhesus macaques did not indicate any issues^13^. The FDA classes rimantadine as a class C drug during pregnancy, *i*.*e*. administered if the potential benefit might outweigh the risk. Whilst rimantadine will cross mouse placentas, eleven times the maximum recommended human dose (MRHD) is necessary to become embryotoxic. However, 3.4 and 6.8 times the MRHD did increase pup mortality up to four days post - partum, yet this still by far exceeds the dose used during influenza prophylaxis in the past (Product Information. Flumadine (rimantadine). Forest Pharmaceuticals, St. Louis, MO. 2007). No controlled studies have been undertaken during human pregnancy, but faced with the alternative of microcephaly, the balance of risk may be favourable. The generic status of rimantadine also means that it could be deployed rapidly at minimal cost to areas of endemic ZIKV infection or during outbreaks, which overwhelmingly tend to occur in lower/middle income countries (LMIC).

In summary, this work supports the formation of membrane channels by the ZIKV M protein, consistent with viroporin activity. Blocking this activity with small molecules reveals an important role during the early stages of ZIKV replication, consistent with virus entry and/or uncoating, and this also translates to antiviral efficacy a preclinical model. Rimantadine may constitute an easy-to-access generic drug for the treatment and/or prophylaxis against ZIKV in LMIC. Moreover, the selection of new repurposed ligands with anti-M activity demonstrates potential for developing novel therapies targeting this protein function, which in turn may also apply to other *Flaviviruses*. In addition, the relative enrichment of effective ligands versus both a lumenal and a peripheral binding site, combined with improved potency compared to rimantadine, further demonstrates the value of molecular models as templates for future drug development. Confidence in such models will be further enhanced via iterative drug development to generate a structure-activity relationship (SAR) for inhibitors targeting each site. In the longer term, structural information on M channel structures, ideally located within virion membranes, will be essential to both developing novel therapeutics, and then in turn, using these to investigate the precise roles for M channel function during ZIKV infection.

## Materials and Methods

### Cell culture

Vero (African Green Monkey kidney) and BHK-21 (baby hamster kidney) cells were cultured in Dulbecco’s modified essential cell culture media, supplemented with 10 % FCS and 100 units/ml penicillin and 0.1 mg/ml streptomycin, at 37 °C in 5 % CO2 in a humidified culture incubator. Cells were passaged every 2-3 days using Trypsin/EDTA (SIGMA), sub-dividing cultures using ratios between 1:5 and 1:10, depending upon confluency. C6/36 cells (derived from *Aedes albopictus* mosquitoes) were cultured in L-15 media, supplemented with 10 % TPB, 10 % FCS and 100 units/ml penicillin and 0.1 mg/ml streptomycin, at 28 °C with no added CO_2_.

### Virus stocks

ZIKV/H. sapiens/Brazil/PE243/2015 (PE243) Zika virus^37^ was obtained from a patient in Recife. It has been sequenced and was supplied (AK laboratory) as a frozen viral stock. This was grown once in BHK-21 cells then passaged once in C6/36 cells, titrated, and stored frozen (-80 °C) at 6 x10^6^ PFU/mL. New stocks for use in cell culture were generated in house at 1.6 x10^6 PFU/mL in Vero cells (cell culture assays). C6/36 derived stocks were used for *in vivo* assays. Briefly, ∼6 x 10^6^ Vero cells were seeded into a T75 (Corning) and left to settle over 4 hr. Cells were then washed once in PBS, prior to addition of PE243 virus in complete DMEM media +10 mM HEPES (Gibco), at a multiplicity of infection (MOI) of 0.001 PFU/cell. Infections were allowed to proceed for 1 hr at 37°C, 5 % CO2, then supernatants were replaced with fresh media. Development of ∼60 % cytopathic effect (CPE) led to harvesting and clarification of supernatants (3184 x *g*, 20 min, 4 °C, in an Eppendorf 5810 R centrifuge). Aliquots were then titred by plaque assay (see below), snap frozen , and stored at -80 °C.

### Determination of virus titre and inhibitor assays

Plaque assays for titration (Vero) or plaque reduction (BHK21) assays were performed upon cells at 80 % confluency in 12-well plates. For titration,10-fold serial dilutions of viral supernatants were made in 0.75 % PBSA (PBS containing 0.75 % bovine serum albumin) and 200 μL added to wells for 1 hr with rocking every 15 min. 2 mL overlay media were then added (2X MEM medium (Gibco), 4 % FCS (Gibco), 200 units/mL penicillin and 0.2 mg/ mL streptomycin, mixed with viscous 1.2 % Avicel (FMC Biopolymer)) and cells incubated for 5 (Vero) or 3 (BHK21) days at 5 % CO_2_ and 37 °C. Supernatants were removed and cells fixed in PBS/10 % paraformaldehyde (PFA) for one hr at 4 °C prior to staining with 0.1 % Toludine Blue (Sigma) for 30 min. Plaque reduction assays comprised duplicate wells per condition with 10 PFU added in the presence of inhibitor or DMSO solvent control (maximum of 0.1 % v/v).

Single cell infectious assays were conducted upon Vero cells seeded at 2000 cells per well in a 96 wells cell culture dish (Greiner Bio-one). After settling for 4 hr, cells were incubated for one hour with virus stock diluted in complete medium at an MOI of 1 PFU/cell. Following infection virus containing media was removed, cells were washed with PBS and replaced with fresh cel l culture media, prior to incubating at 37 °C and 5 % CO2 for 48 hr in a humidified incubator. Cells were then washed three times in PBS, fixed using 4 % PFA for 10 min at RT, and stained for ZIKV E. Briefly, cells were permeabilised using 0.1 % TX-100 in PBS at RT for 10 min, then stained using a mouse monoclonal anti-ZIKV E antibody (Aalto Bio Reagents #AZ1176) diluted 1:500 in 10 % v/v FCS in PBS overnight at 4 °C. Cells were washed three times in PBS prior to adding goat anti-mouse Alexa Fluor 488 nm -conjugated secondary antibody, diluted 1:500 in 10 % v/v FCS in PBS at RT for 1 hr. Cells were washed another three times in PBS and left in PBS for imaging. Infectious units (FFU) were determined using an IncuCyte Zoom (Essen Bioscience) to determine numbers of fluorescently labelled cells as well as confluency, as described previously ^67^. The 10 x objective was used to take four images covering each well, with positive and negative control wells allowing modification of the processing definition parameters for optimal detection.

### Preclinical ZIKV model

*In vivo* animal models were approved by the University of Leeds local ethics review committee. Procedures were carried out in accordance with the United Kingdom Home Office regulations under the authority of the appropriate project and personal license (awarded to CSM, and CSM/DL respectively). Wild type C57BL/6j mice were bred in-house and maintained under specific pathogen-free conditions. Mice were age and sex matched in all *in vivo* experiments, and used between 4 and 12 weeks of age.

Mice were dosed with 1.5 mg InVivoMAb anti-mouse IFNAR-1 antibody 24 hr prior to virus inoculation, then a sub-cutaneous dose of rimantadine hydrochloride at 20 mg/kg (or carrier control) 30 min prior to ZIKV infection. Mice were then anaesthetised using 0.1 mL/10 g of Sedator/Ketavet via intraperitoneal (I.P.) injection, then placed on top of mosquito cages using foil to expose only the dorsal side of one hind foot. No more than 5 mosquitoes were allowed to feed on each mouse. 2000 PFU of C6/36 derived ZIKV was injected directly into the bite site using a 5 μL 75N syringe, 26ga (Hamilton) using small RN ga33/25mm needles (Hamilton). A second bolus of rimantadine was administered 5 hr post-infection. Mice were observed four times throughout the 24-hr experiment, weighed once, and culled 24 hr post-infection.

Mice were culled via a schedule 1 method. Skin from the bitten foot was dissected and placed in 0.5 mL RNAlater (Sigma Aldrich, USA) in 1.5 mL tubes. Blood was collected from the ventricles by cardiac puncture, and then centrifuged to isolate serum and stored at -80 °C.

### Mosquito rearing

*Aedes aegypti* (Liverpool strain) mosquitoes were reared at 28 °C with 80 % humidity conditions with a 12 h light/dark cycle. Eggs were hatched overnight from filter papers in trays containing approximately 1.5 cm depth of water. Larvae were fed Go-cat cat food until pupation. Pupae were placed in water-filled containers inside BugDorm mosquito cages where they were left to emerge. A 10 % w/v sucrose solution was fed to adult mosquitoes. Mosquitoes were ready for biting experiments from 21 days post-hatching.

### RNA purification and quantification

Skin tissue was lysed in 1mL Trizol (Invitrogen/Thermo) by shaking with 7 mm stainless steel beads at 50 Hz for 10 min on a Tissue Lyser. 200 μL chloroform was added, and samples mixed by inversion, prior to centrifugation in a microcentrifuge at 12 000 x *g* for 15 min at 4 °C and removal of the top aqueous phase to a fresh tube containing an equal volume of 70 % (v/v) EtOH. RNA was then extracted using a PureLink RNA Mini kit (Life Technologies) according to manufacturer’s instructions. cDNA was synthesised from 1μg of RNA using the “Applied Biosystems High-Capacity RNA to cDNA” kit, according to manufacturer’s instructions.

cDNA was diluted 1 in 5 in RNAse free water, then introduced into a master mix comprising cDNA, primers, water and SYBRÒ green mix. A non-template control (NTC) comprising RNAse free water in place of cDNA was also included. Triplicate technical replicates were made for each biological replicate and a standard curve generated by 10-fold serial dilution of the 10^-2^ PCR standard. PCR plates were run on an Applied Biosystems Quantstudio 7 flex machine. Ct value was calculated automatically using the Quantstudio software, which detects the logarithmic phase of the PCR reaction. Samples were quantified according to their position on the standard curve, which was required to have close to 100 % efficiency, indicated by R^2^≥0.998 and a slope of 3.3. Melt curves were conducted to control for primer specificity. Analysis of qPCR data was done with Microsoft Excel by the use of the median of the technical replicates and normalizing them to the median of the technical replicates of the housekeeping genes. GraphPad Prism software was used to generate graphs and the non-parametric Mann-Whitney test was used for comparisons between two groups.

### Western blotting

6-well plates seeded with 1 x 10^5^ Vero cells were infected with ZIKV at an MOI of 0.1 PFU/cell as above, then media applied containing increasing concentrations of rimantadine or a DMSO solvent control (0.1 % v/v). Cells were harvested at 48 hr post infection by washing three times in 1 mL PBS and scraping into a 1.5 mL Eppendorf tube. Cells were pelleted at 10 000 x *g* in a microcentrifuge, then whole cell lysates were made in 200 μL Enriched Broth Culture (EBC) lysis buffer [50 mM Tris HCl, pH 8.0, 140 mM NaCl, 100 mM NaF, 200 μM Na_3_VO_4_, 0.1 % (v/v) SDS, 1 % Triton X100, Roche complete ULTRA protease inhibitor cocktail]. Lysates were normalised for protein concentration using a Pierce BCA Protein Assay Kit (Thermo Fisher Scientific), then diluted with an equal volume of 2x Laemmli Buffer (100 mM Tris HCl pH 6.8, 4 % (v/v) SDS, 20 % (v/v) Glycerol, 10 mM DTT, 0.025 % (w/v) Bromophenol Blue). Lysates were denatured by heating for 10 min at 95 ^0^C prior to separation on hand-cast Tris-Glycine polyacrylamide gels. Proteins were resolved by SDS-PAGE (25 mM Tris-Cl, pH 8.0, 250 mM Glycine, 0.1 % SDS) then transferred to a polyvinylide fluoride (PVDF, Immunoblot-FL Merck Millipore) membrane, pre-activated in 100 % MeOH, using a Hoeffer semi-dry transfer rig, sandwiched in blotting paper soaked in transfer buffer (25 mM Tris-Cl, 250 mM Glycine, pH 8.3, 20 % v/v MeOH). Transfers proceeded for 1-2 hr at 120-240 mA constant current, depending on the number of gels. Membranes were blocked in 5 % w/v fat-free milk in TBS-T (Tris-buffered saline (50 mM Tris-Cl, pH 7.5, 150 mM NaCl) with 0.1 % v/v Tween 20, Sigma-Aldrich) for 4 hr at RT with gentle shaking, prior to incubation with primary antibodies diluted in 5 % (w/v) BSA in TBS-T at 4 °C overnight with gentle shaking (Anti-E 1:10000, mouse monoclonal, Aalto Bio Reagents #AZ1176). Membranes were then washed three times for 10 min in TBS-T, prior to incubation with secondary antibody (1/10000 goat anti-mouse IgG-Horseradish peroxidase (HRP) conjugate, Sigma #A4416). Blots were visualised using ECL prime western blotting detection reagent (GE Healthcare Life Sciences) by exposure to X-ray film, with protein sizes determined by comparison with pre-stained molecular weight markers (Seeblue® Plus2, Invitrogen).

### Purification and density gradient separation of ZIKV virions

This was undertaken using similar methods to our previous studies of HCV(49). Briefly, 10 mL of infectious ZIKV culture Vero cell supernatants were clarified and concentrated by the addition of PEG-8000/PBS to a final concentration of 10 % w/v, mixing by several inversions, and incubation overnight at 4 ^0^C. Virion concentrates were pelleted at 2000 rpm in a benchtop centrifuge and then resuspended in 0.5 mL PBS. This was then layered over a 1 mL 20 % w/v in PBS sucrose cushion and virions pelleted at 100K x g in a Beckmann TLS-55 rotor at 4 ^0^C for three hours. Pellets were left to resuspend for 1 hr at 4 ^0^C in 200 μL PBS, then layered over a 10-40 % v/v iodixinol/PBS gradient. These were spun in the same conditions as above but for 2 rather than 3 hr. Gradients were fractionated from the top into 24 equal samples by careful pipetting, then 10 μL diluted into media to assess infectivity in Vero cells seeded into 96-well plates as described above. Virion RNA was harvested from samples and quantified as described above. Western blotting was performed upon 15 μL of gradient sample with 5 μL 4x Laemmli sample buffer added followed by boiling for 10 minutes prior to SDS -PAGE and immunoblotting for E protein as described above.

### Native PAGE

M peptide was solubilised at 37 °C for 10 min in 300 mM detergent (1,2 -dihexanoyl-sn-glycero-3-phosphocholine (DH(6)PC), 1,2-diheptanoyl-sn-glycero-3-phosphocholine (DH(7)PC), 1-palmitoyl-2-hydroxy-sn-glycero-3-phospho-(1’-rac-glycerol) (LPPG) and 1-myristoyl-2-hydroxy-sn-glycero-3-phospho-(1’-rac-glycerol) (LMPG)) in Liposome Assay Buffer (10 mM HEPES, pH 7.4, 107 mM NaCl). Native-PAGE loading dye (150 mM Tris-Cl pH 7.0, 30 % (v/v) Glycerol, 0.05 % (w/v) bromophenol blue) was added to samples, which were loaded onto gradient precast gels (4 -20%) (TGX, Bio-Rad) and run using Native-PAGE running buffer (250 mM Tris-Cl, pH 8.5, 192 mM Glycine) at 140 V for 1 hr. Gels were stained with Coomassie Brilliant Blue (0.1 % (w/v) Coomassie Blue, 10 % acetic acid, 50 % MeOH). Unstained SDS-free molecular weight markers (Sigma-Aldrich) were used to estimate protein size.

### Synthetic M peptide

M peptide with an N-terminal truncation (residues 18-75) was manufactured by Alta Bioscience, provided as a lyophilised powder at >95 % purity based upon HPLC. Peptides contained the sequence:

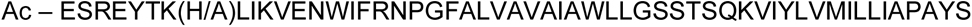

### Transmission Electron Microscopy

5 μg of M peptide was incubated in 10 mM HEPES, 107 mM NaCl and varying concentrations of DH_(6)_PC for 10 min at RT. Sample was added to copper grids with a continuous amorphous carbon film, before washing with water, then stained with 1 % uranyl acetate. Grids were examined using a Tecnai F20 at 120 kV on a FEI CETA camera with a sampling of 4.18 Å per pixel. Particles were manually picked and 2D class averaging were carried out using RELION 3 software^68^.

### Liposome dye release assay

L-α-phosphatidic acid (α-PA), L-α-phosphatidyl choline (α-PC) and L-α-phosphatidyl ethanolamine with lissamine rhodamine b labelled head groups (α -PE) from chicken eggs, supplied at 10 mg/ml in chloroform, were purchased from Avanti Polar Lipids. Aliquots were made using Hamilton glass syringes in glass vials, then stored at -80 °C. Lipids were combined into a glass tube on ice under non-oxygen gas (Nitrogen), as a total of 1 mg (50 μL PA, 50 μL PC & 5 μL PE). Lipids were dried initially under Nitrogen, then overnight using a vacuum dessicator at RT. Lipids were then rehydrated in carboxyfluorescein (CF)-containing buffer (50 mM CF 10 mM HEPES-NaOH pH 7.4 107 mM NaCl) to 2 mg/mL, with vigorous shaking overnight at RT). The following day, liposomes were generated via extrusion (15 passes) using an Avanti mini extruder housing a 0.4 μM filter (Whatmann), at 37 °C. Resulting unilamellar liposomes were washed at least three times in liposome assay buffer (10 mM HEPES pH 7.4, 107 mM NaCl) to remove unincorporated CF, pelleting liposomes at 100 000 x *g* for 15 min at RT using a MLS-50 rotor in a Beckman Coulter TLX ultracentrifuge. Final pellets were resuspended in assay buffer (500 μL) and concentration determined by comparing rhodamine absorbance (OD_570 nm_) in pre-extrusion (diluted 1:20) and final samples:

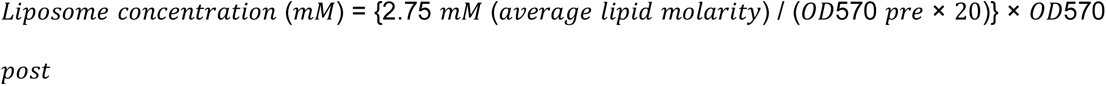

Dye release assays were conducted using up to 800 nM peptide (480 nM standard) in DMSO (maximum of 5 % v/v DMSO per well), with 50 μM liposomes in a total reaction volume of 100 μL. Inhibitors were pre-incubated with peptide for 5 min in flat-bottomed, black 96-well plates (Greiner Bio One) at RT, prior to the rapid addition of chilled liposome suspensions in assay buffer. CF release compared to solvent controls and liposomes alone was measured over 30 min at 37 °C with initial mixing, using a FLUOstar Galaxy Optima plate-reader (BMG Labtech, λ_ex_ 485 nm / λ_em_ 520 nm. Gain adjustment at 90 % total fluorescence was set using a pre-measured sample containing 0.5 % v/v Triton X-100 to lyse all liposomes present. End-point values from three technical repeats were calculated for each averaged biological repeat and significance between the latter determined using multiple paired student T-tests. Assays assessing pH dependence utilised assay buffers at stated pH, with endpoint supernatants re-buffered using 2.5 μL 1M Tris-Cl [pH7.5] prior to clarifying by ultracentrifugation as above and determining fluorescence.

### *In silico* analysis of M protein

Alignment of *Flavivirus* M sequences was performed using Clustal Omega (https://www.ebi.ac.uk/Tools/msa/clustalo/), with outputs visualised using BOXSHADE 3.21 (https://embnet.vital-it.ch/software/BOX_form.html), and curated in Jalview. Conservation outputs were fed into an online tool (https://weblogo.berkeley.edu/) to generate a protein sequence LOGO to illustrate the high degree of conservation between >700 isolates.

M protein tertiary structures were taken from PDB: 5IRE. Online servers used to determine probability of *trans*-membrane regions comprised TMHMM v2.0 (http://www.cbs.dtu.dk/services/TMHMM/), TOPCONS (https://topcons.cbr.su.se/), Phobius (https://phobius.sbc.su.se/), SPLIT v4.0 (http://splitbioinf.pmfst.hr/split/4/) and MEMSAT-SVM (https://bio.tools/memsat-svm). Amino acid properties were determined using EMBOSS Pepinfo (https://www.ebi.ac.uk/Tools/seqstats/emboss_pepinfo/).

### Coarse-grained Molecular Dynamics simulations

Coarse-grained (CG) Molecular Dynamic simulations of monomeric M proteins were performed using Martini v2.2 force field^69^ and GROMACS^70^. The cryo-EM structure of ZIKV M protein structure (PDB: 5IRE)^25^ was converted to a coarse-grained resolution using the “martinize” script. An elastic network model was used in the monomer simulations to maintain the secondary structure, yet not the tertiary structure. The cryo-EM structure was used to simulate the TMD model, whereas “Modeller” ^71^ was used to generate the second model, straightening the linker region between the two TMDs into a longer TMD. Five repeat simulations of 3 μs each were run for each system.

A POPC bilayer was built using INSANE (INSert membrANE) CG tool ^72^. Systems were solvated with CG water particles and ions were added to neutralise the system to a final concentration of 150 mM NaCl. Prior to simulation, systems were energy minimised using the steepest descent algorithm for 500 steps in GROMACS and equilibrated for 10 ns with the protein backbone restrained. The temperature was set at 323 K and controlled by V-rescale thermostat (coupling constant of 1.0)^73^. Pressure was controlled by Parrinello-Rahman barostat (coupling constant of 1.0 and a reference pressure of 1 bar)^74^. Integration step was 20 fs.

### Atomistic Molecular Dynamics Simulations

The all-atom hexameric and heptameric M protein oligomers were first energy minimised prior to conversion into CG using the Martini 2.2 forcefield and as above inserted into the bilayer system using INSANE. The systems were then equilibrated in CG restraining the protein. The systems were then converted back into atomistic resolution using the Martini backward tool^75^. Simulations were then energy minimised, equilibrated for 20 ns with the protein Cα atoms restrained and run using the CHARMM36 force field^76^. Three repeat simulations were run for 200 ns for each system. Temperature and pressure were controlled using the v-rescale thermostat^73^ and Parrinello-Rahman barostat^74^ respectively. Bond lengths were kept constant using the LINCS algorithm^77^. The time-step was 2 fs and the temperature set to 323 K.

### Design of Hexameric and Heptameric M channel structures

A python script was used to calculate the co-ordinates of each monomer within the oligomeric structure, a radii of 1.3 nm was used for hexamer:

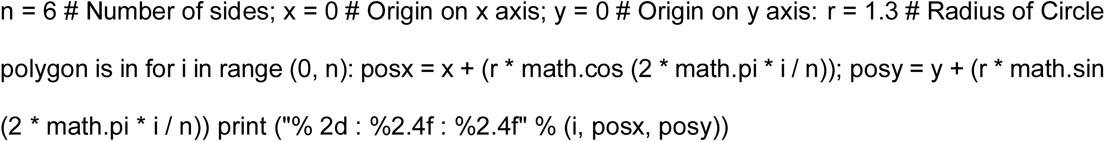

### *In silico* docking and virtual HTS

Maestro (Schrödinger) was used for assessing ligand interactions with the compact hexameric M channel structure. This was minimised in a lipid membrane environment using an Optimised Potentials for Liquid Simulations (OPLS) force field and used for docking of unbias ed compound analogues. The Maestro SiteMap function was used to assess the druggability score for three potential lumenal binding sites (L1-3), and another on the membrane-exposed periphery was identified manually.

For unbiased screening, a library of 1280 FDA-approved molecules (TOCRIS Screen, Biotechne. https://www.ebi.ac.uk/Tools/seqstats/emboss_pepinfo/) underwent the LigPrep function using Maestro (Schrödinger) software. The in-built SiteMap function was then used on the M Protein structure to generate a Glide grid for the docking of Rimantadine in the lumenal site (under aqueous conditions) using extra precise (XP) Glide setting. This was followed by docking in the same site mimicking membrane conditions. Rimantadine was then docked into the peripheral site under both aqueous and membrane conditions to generate 32 poses in each of the four scenarios. The prepared Tocris library was also docked into the two sites on the M protein structure using the four scenarios and Glide settings established using Rimantadine (lumenal-aqueous, lumenal-membrane, peripheral-aqueous, and peripheral-membrane). Duplicate poses were removed, and a final short-list of 50 molecules was generated at each for *in vitro* testing.

## Supporting information

Annexe detailing additional Channel MD simulations

## Acknowledgments

Schematics in figures 1e, 3d, 4a and S2b were generated using Biorender software (https://biorender.com) with a paid subscription. Simulations were undertaken on ARC3 and ARC4, part of the High-Performance Computing facilities at the University of Leeds. Work was supported by Institute PhD Scholarships from the Leeds Institute of Medical Research, University of Leeds, (EB and DF: awarded to SG, CM, AK, and RF), Medical Research Council grant G0700124 (S.G.), and Medical Research Council grants (MC_UU_12014/8 and MR/N017552/1) (A.K.).

## Supplementary Figures

**Figure S1.**
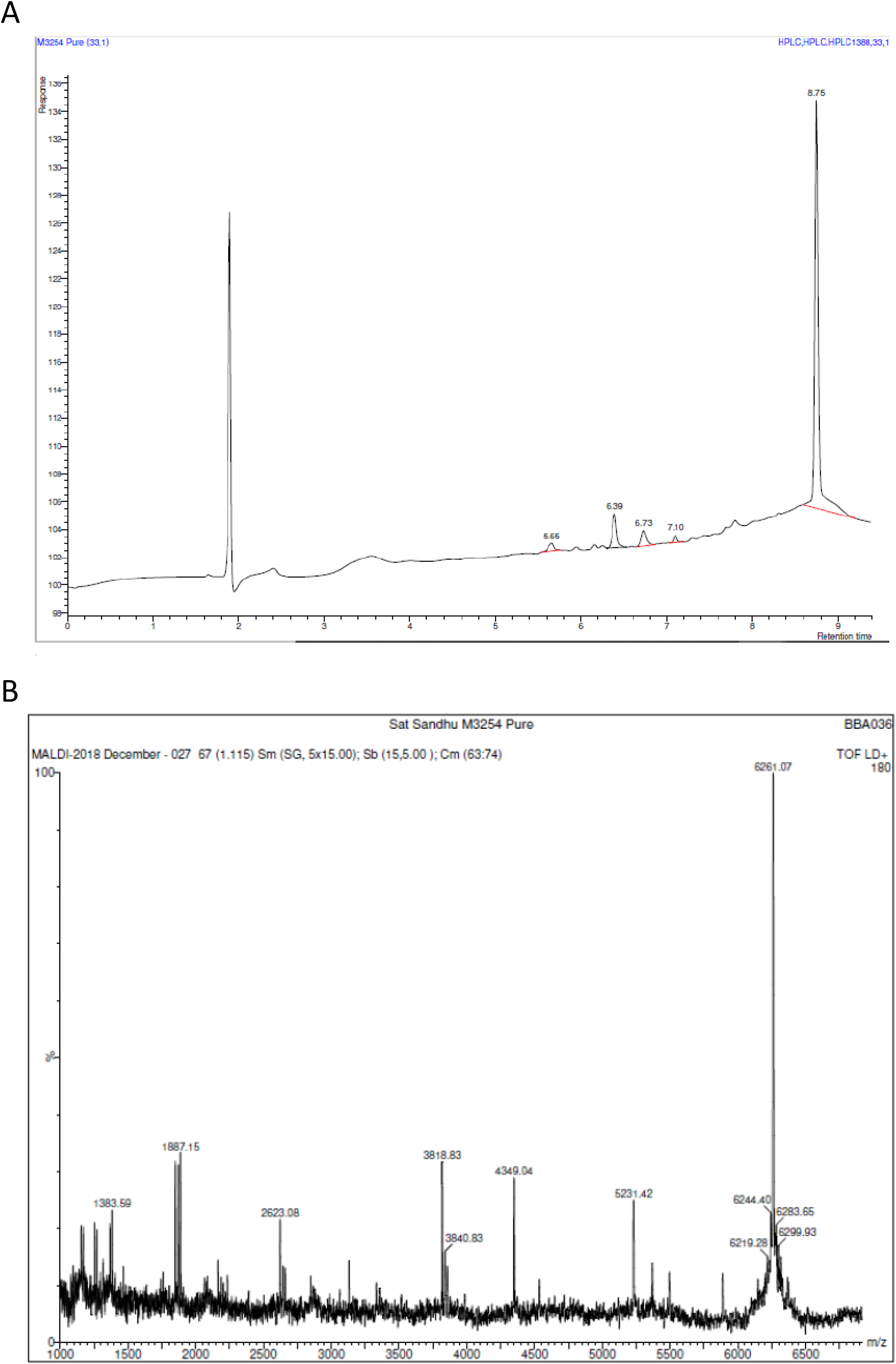
Example HPLC and mass spectrometry analysis of synthetic M peptides supplied by Alta Biosciences.

**Figure S2.**
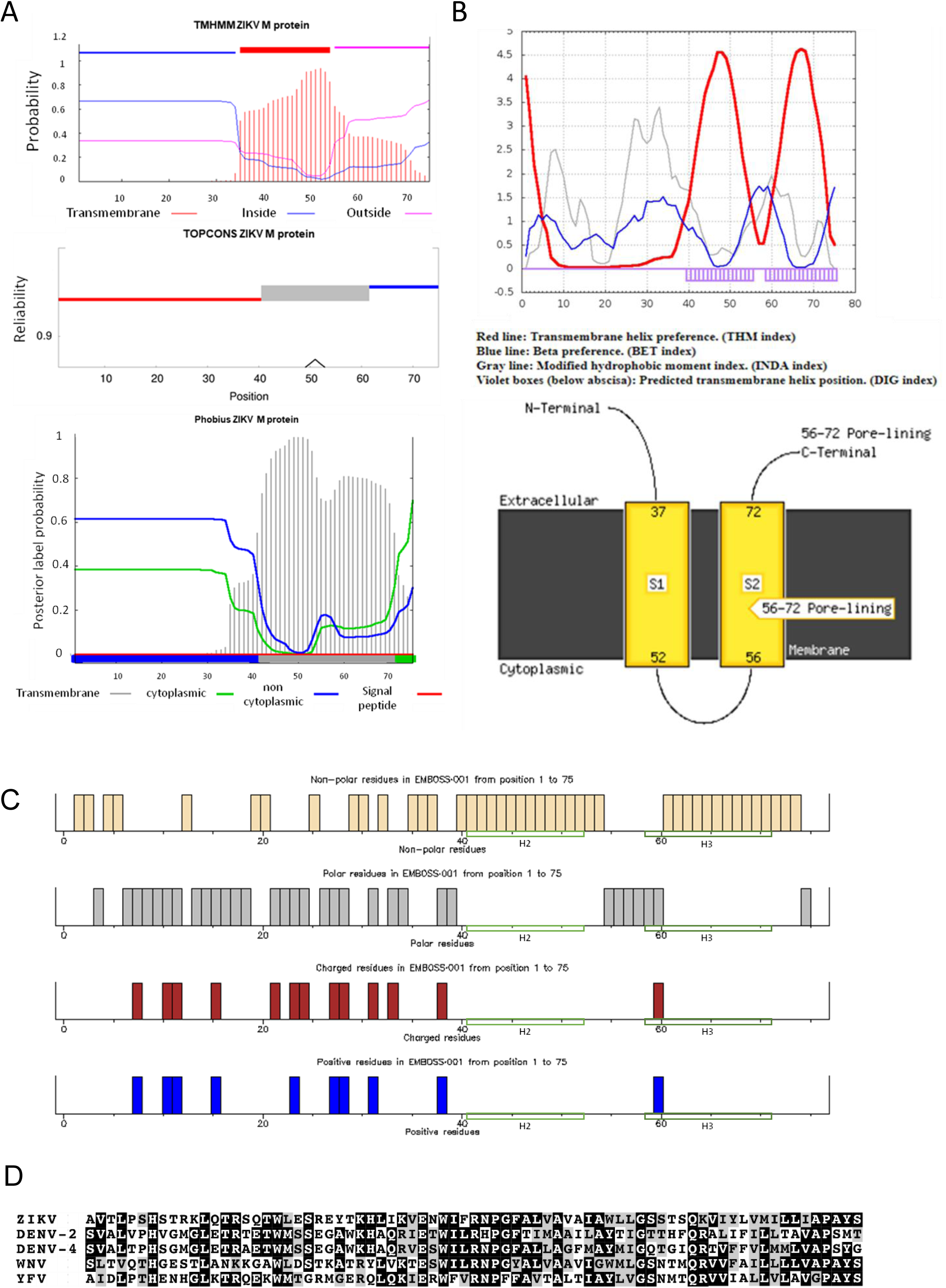
**A**. Prediction of M protein *trans*-membrane regions, topology and hydrophobic profiles generated in different online server packages where single TMD topology was supported: TMHMM, TOPCONS, and Phobius. **B.** As for A, but dual TMD topology: SPLIT v4.0 and MEMSAT-SVM. **C.** M sequence amino acid characteristics visualised using EMBOSS Pepinfo. **D.** Alignment of consensus *Flavivirus* M protein sequences, black – 100% conserved, grey – conserved amino acid properties.

**Figure S3.**
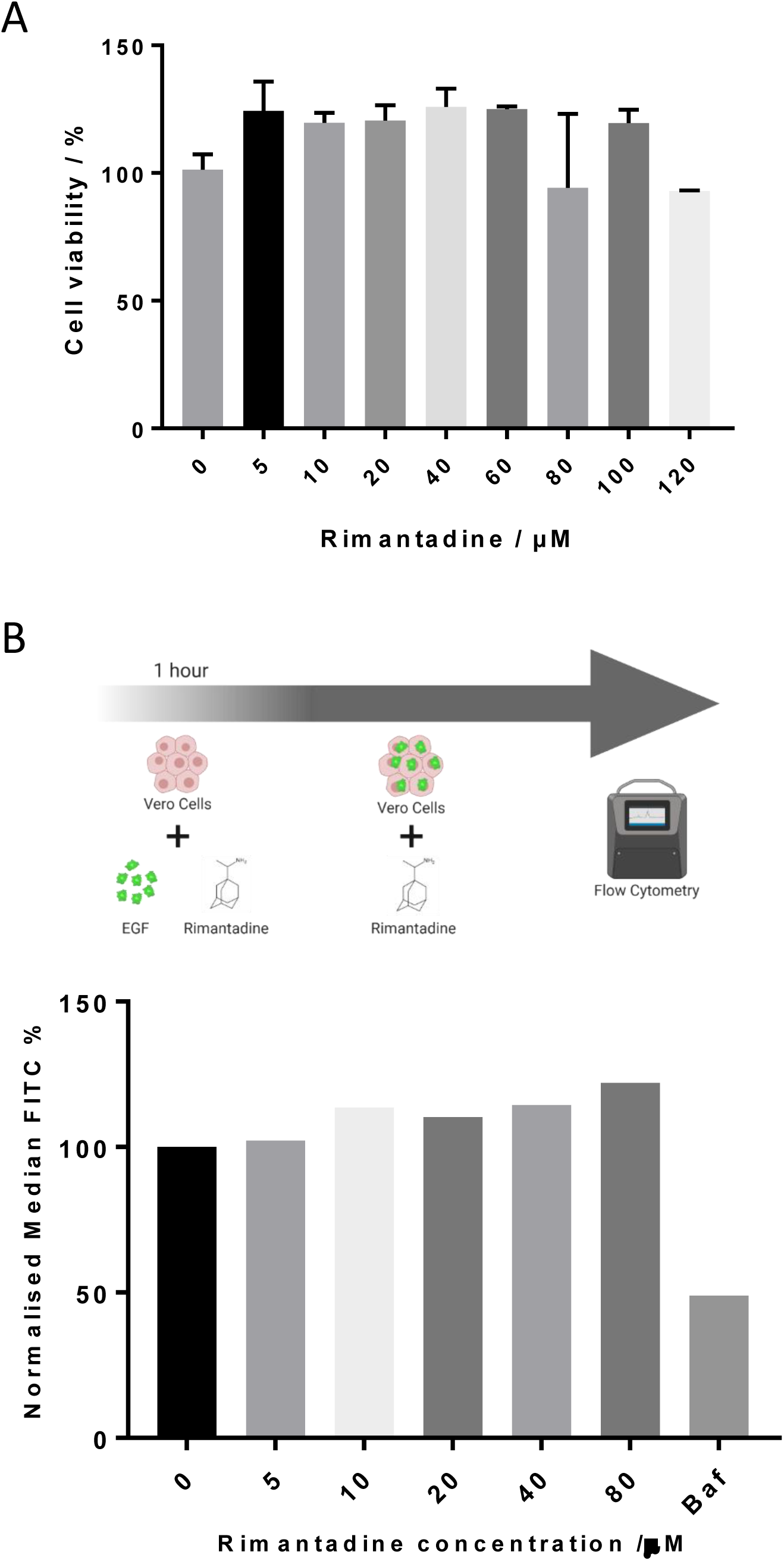
Controls for off-target effects of rimantadine during infectious ZIKV culture. **A.** Viability of Vero cells measured using MTT assay after 48 hr incubation with rimantadine or DMSO control for 48 hr. Graph summarises a single representative biological repeat with three technical repeats. Error bars represent one standard deviation. B. Lack of effect of rimantadine upon uptake of fluorescently labelled EGF by endocytosis. Top – experimental schematic. Bottom – endocytic uptake of fluorescently labelled (FITC) EGF in Vero cells pre-treated with 5 to 80 μM rimantadine. Uptake was measured using flow cytometry and quantified in 25000 cells per condition. Data is expressed as median FITC, displayed as a % of maximum fluorescent EGF uptake at 0 μM rimantadine. Bafilomycin A1 was used at 1 μM as a positive control for blocking endocytic uptake.

**Figure S4.**
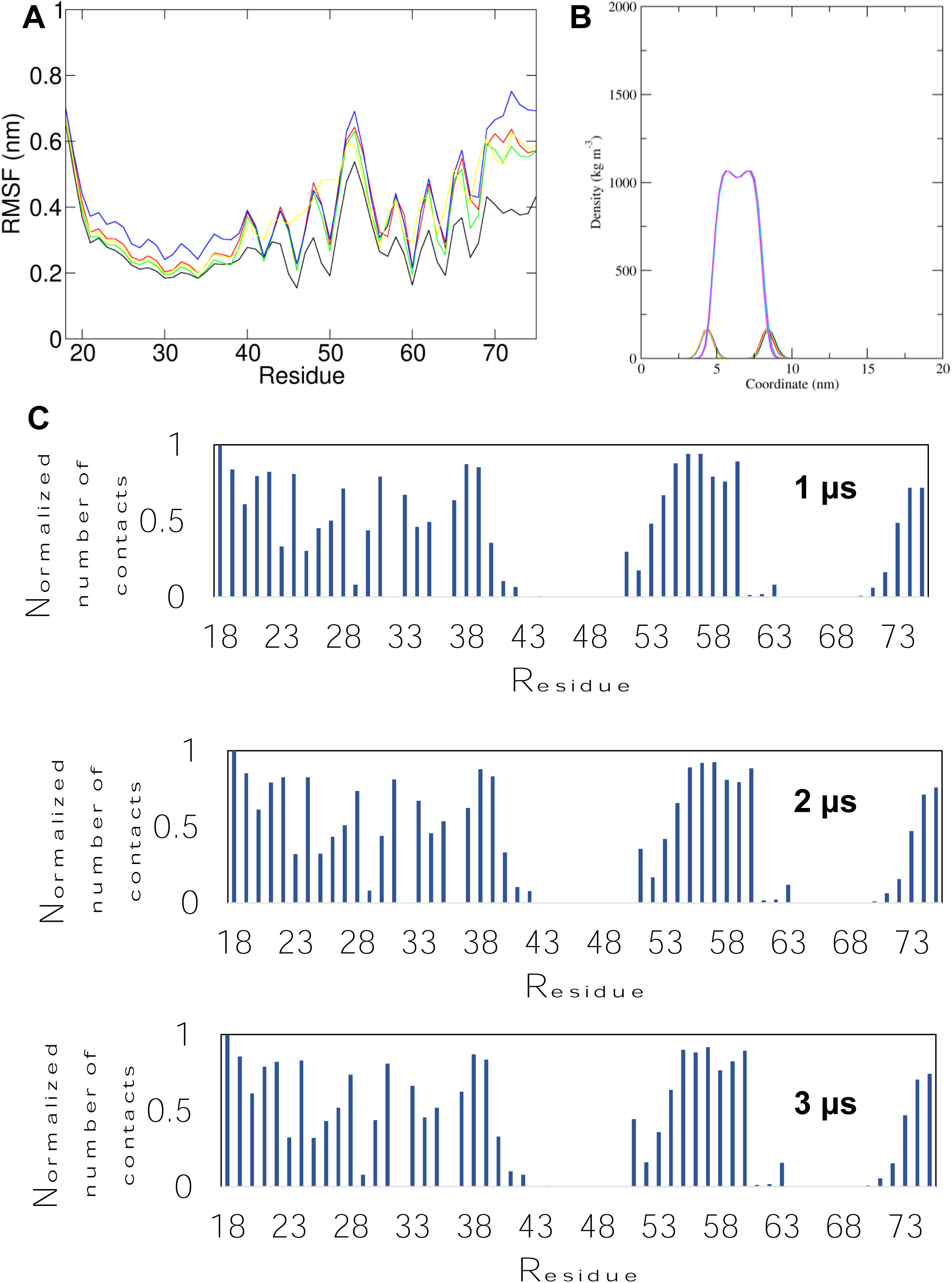
A. Simulation system box for M protein hexamers in hydrated DOPC bilayers, visualised from the side (protein: black; lipids: green and silver; water red and silver), or the top (protein: cyan; lipid: grey). **B.** Overlay of compact (white) vs radial (black) hexamer conformation at neutral pH. **C.** Simulation of M complexes where the N-terminal helix was removed, leading to rapid (< 50 ns) dissociation of the channel complex. **D.** Starting conformations for His28^++^ compact hexamer with lumenal helix 3, showing high similarity to non-protonated forms. **E.** Structures of protonated channels at 100 ns including HOLE profile of channel lumen.

**Figure S5.**
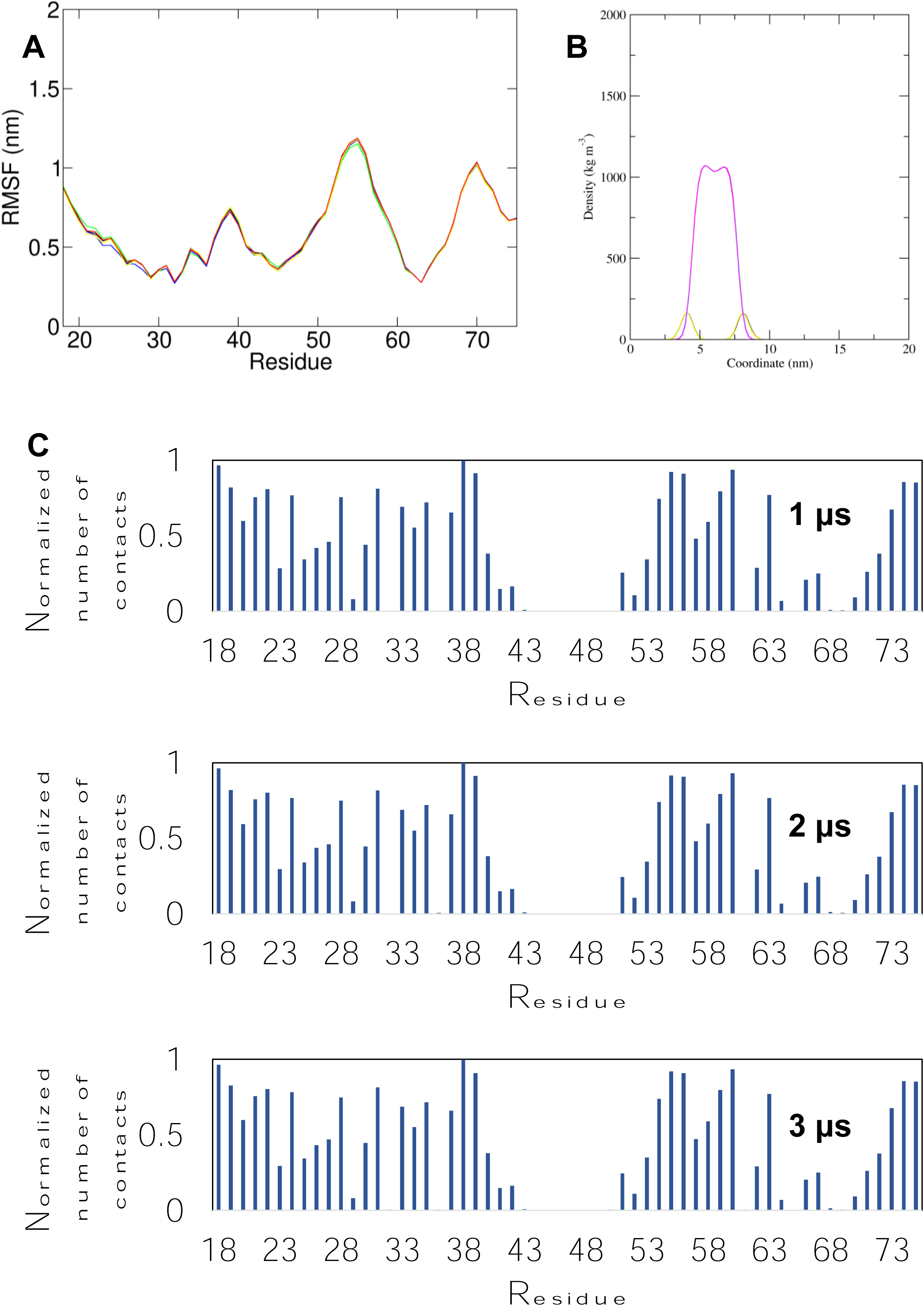
A. Root mean square fluctuation (RMSF) per residue for the simulations in which the cryo-EM derived conformer (i.e. double TMD monomeric M from PDB: 5IRE) was inserted in a POPC membrane. The different colours represent different simulations. **B.** Density profiles along the membrane normal of the POPC head group and the POPC tails for the same simulations. The different colours represent different simulations. **C.** Normalized average number of contacts between the protein and the POPC lipid headgroup across all repeat simulations of the simulation in which the cryo-EM derived conformer was inserted in a POPC membrane shown for 1 μs of the simulation ensemble, 2 μs of the simulation ensemble and for all the simulation ensemble (3 μs). For the normalization the number of contacts of each residue with POPC headgroup was divided by the largest number of contacts.

**Figure S6.**
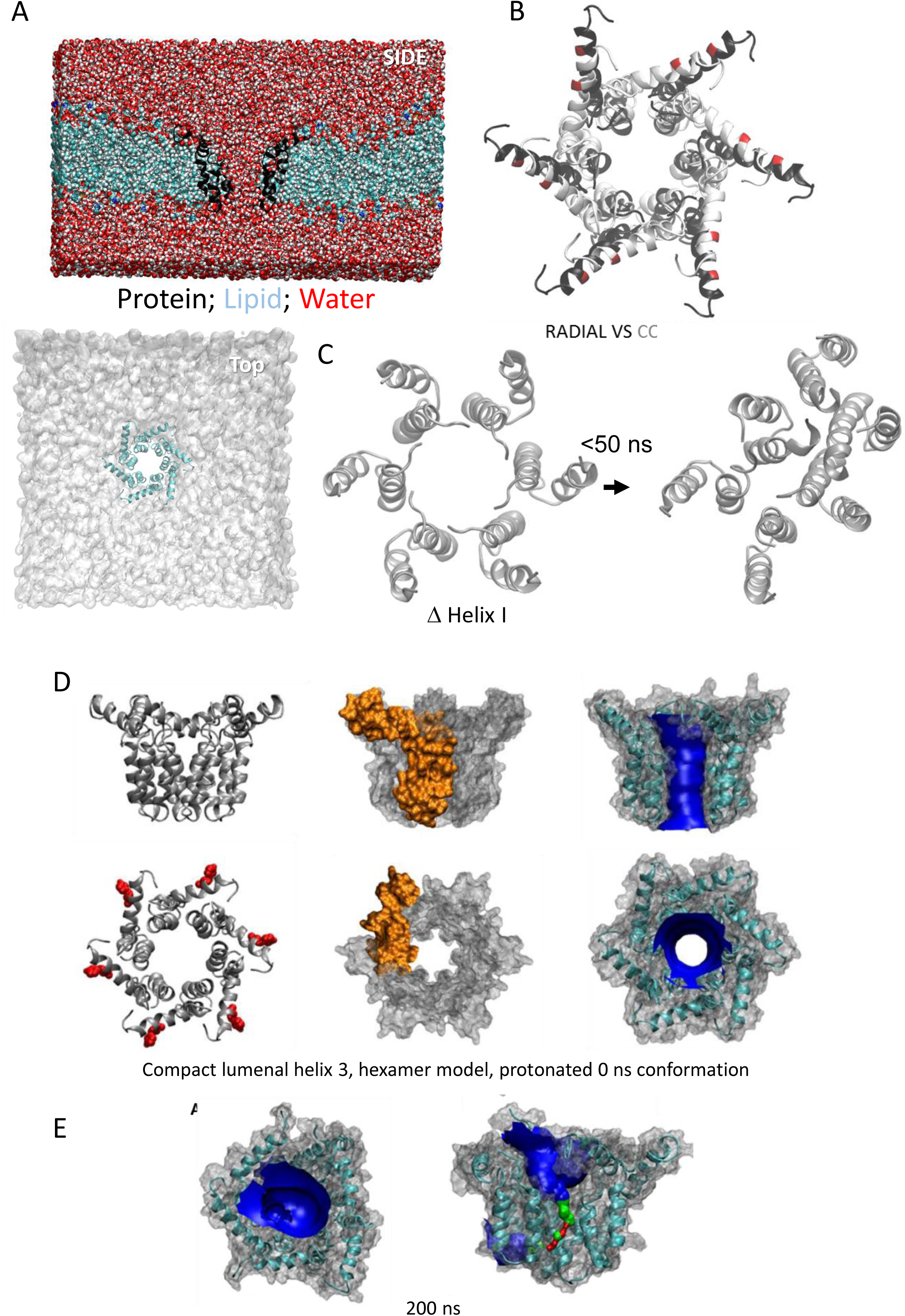
A. Root mean square fluctuation (RMSF) per residue for simulations in which the two transmembrane helices were straightened into a membrane-spanning helix containing a small unstructured region between helix two and three, prior to insertion into a POPC membrane; this was designed to emulate secondary structure predictions where a single TMD was predicted . The different colours represent different simulations. **B.** Density profiles along the membrane normal of the POPC head group and the POPC tails for the same simulations. The different colours represent different simulations. **C.** Normalized average number of contacts between the protein and the POPC lipid headgroup across all repeat simulations of the simulation in which the cryo -EM derived conformer was inserted in a POPC membrane shown for 1 μs of the simulation ensemble, 2 μs of the simulation ensemble and for all the simulation ensemble (3 μs). For the normalization the number of contacts of each residue with POPC headgroup was divided by the largest number of contacts.

**Figure S7.**
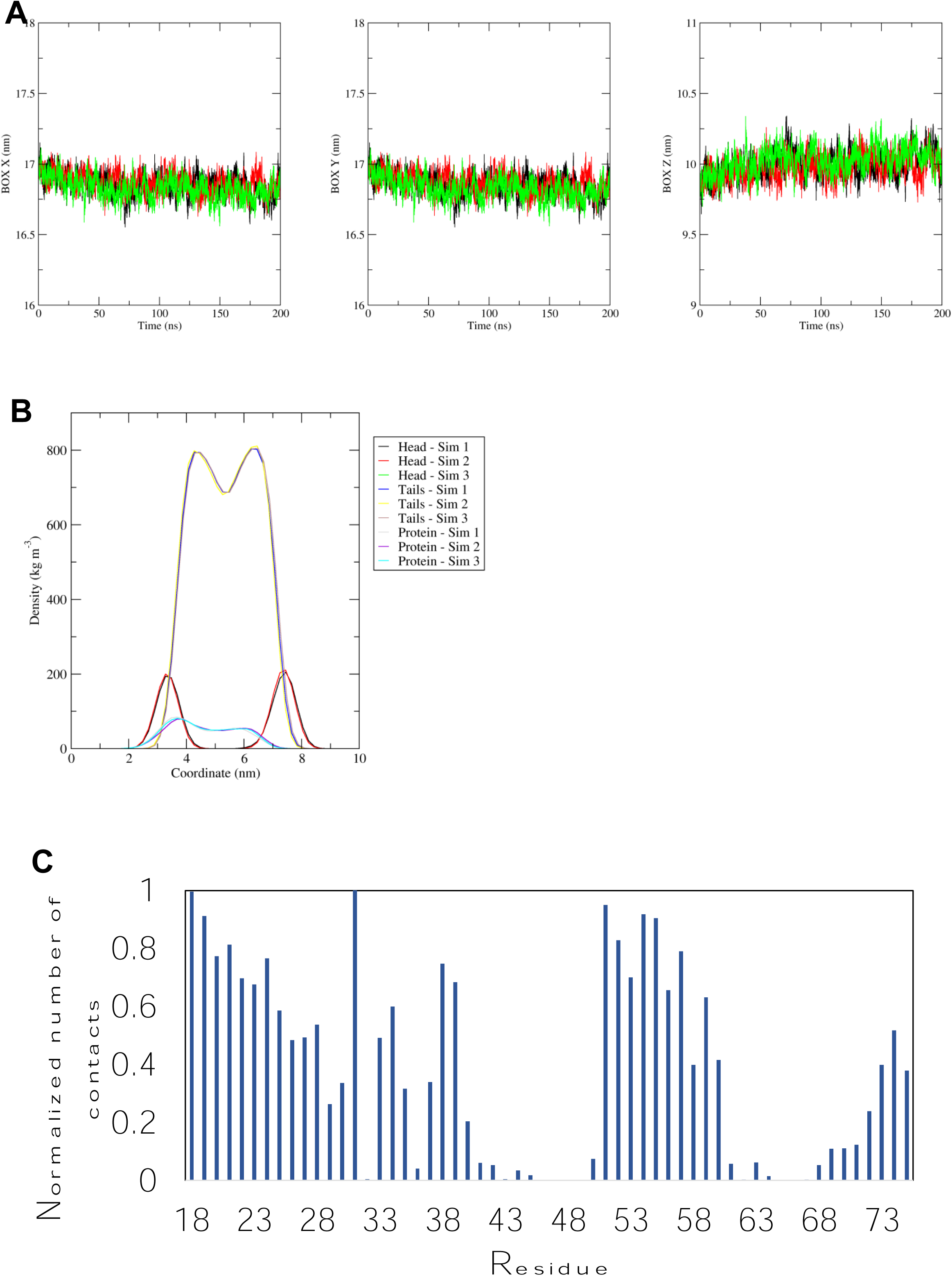
A. Size (in nm) of the X, Y, and Z site of the simulation box for the simulations in which helix one was orientated almost perpendicular to the channel pore with helix three facing towards the pore (radial) of the hexameric channel. **B.** Density profiles along the membrane normal of the POPC head group, the POPC tails, and of the proteins for the same simulations. In A and B the different colours represent different simulations. **C.** Normalized average number of contacts between the protein and the POPC lipid headgroup across all repeat simulations of the same simulation. For the normalization the number of contacts of each residue with POPC headgroup was divided by the largest number of contacts.

**Figure S8.**
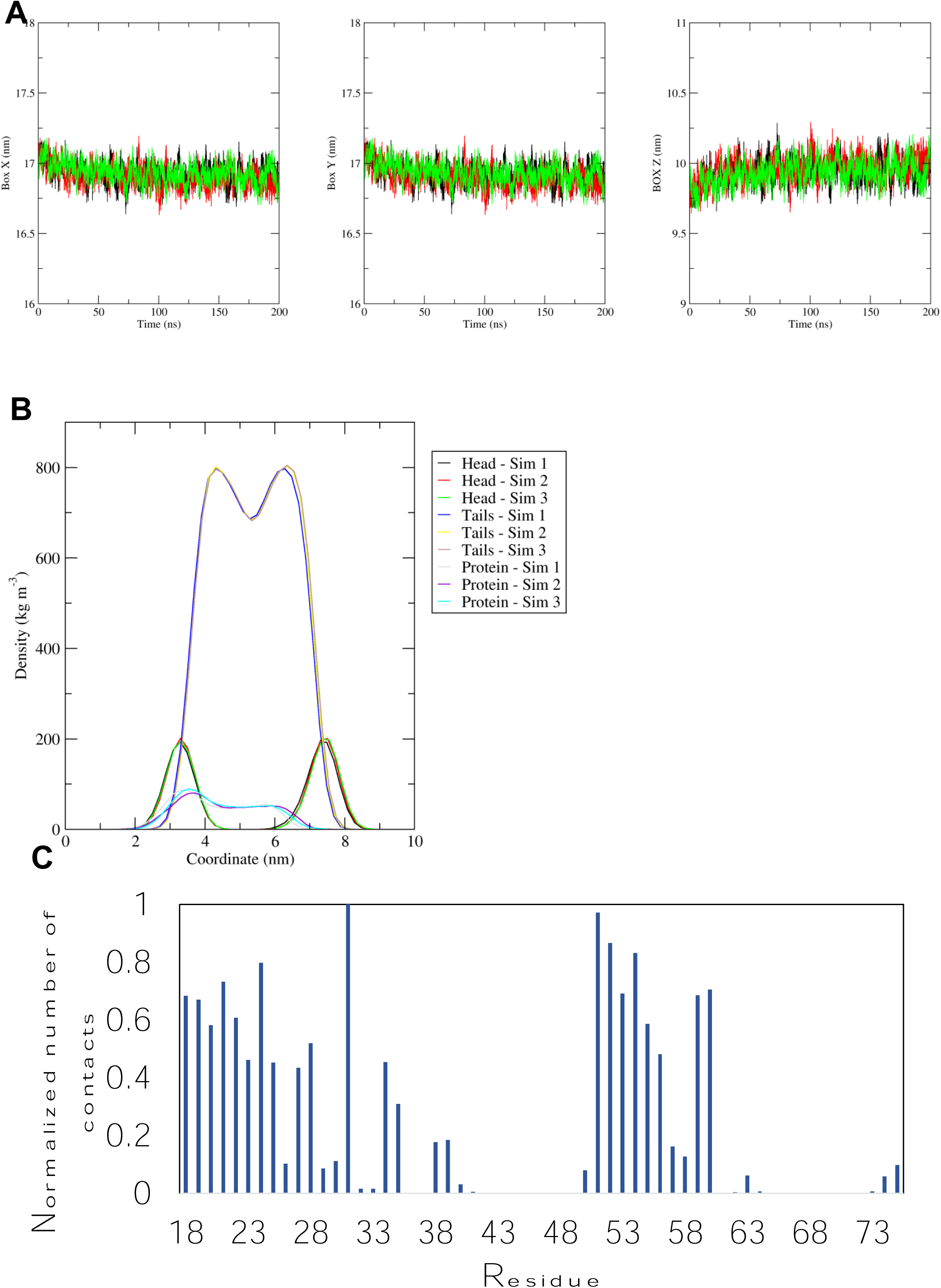
A. Size (in nm) of the X, Y, and Z site of the simulation box for the simulations with the more compact structure (compact) of the hexameric channel. **B.** Density profiles along the membrane normal of the POPC head group, the POPC tails, and of the proteins fo r the same simulations. In A and B the different colours represent different simulations. **C.** Normalized average number of contacts between the protein and the POPC lipid headgroup across all repeat simulations of the same simulation. For the normalization the number of contacts of each residue with POPC headgroup was divided by the largest number of contacts.

**Figure S9.**
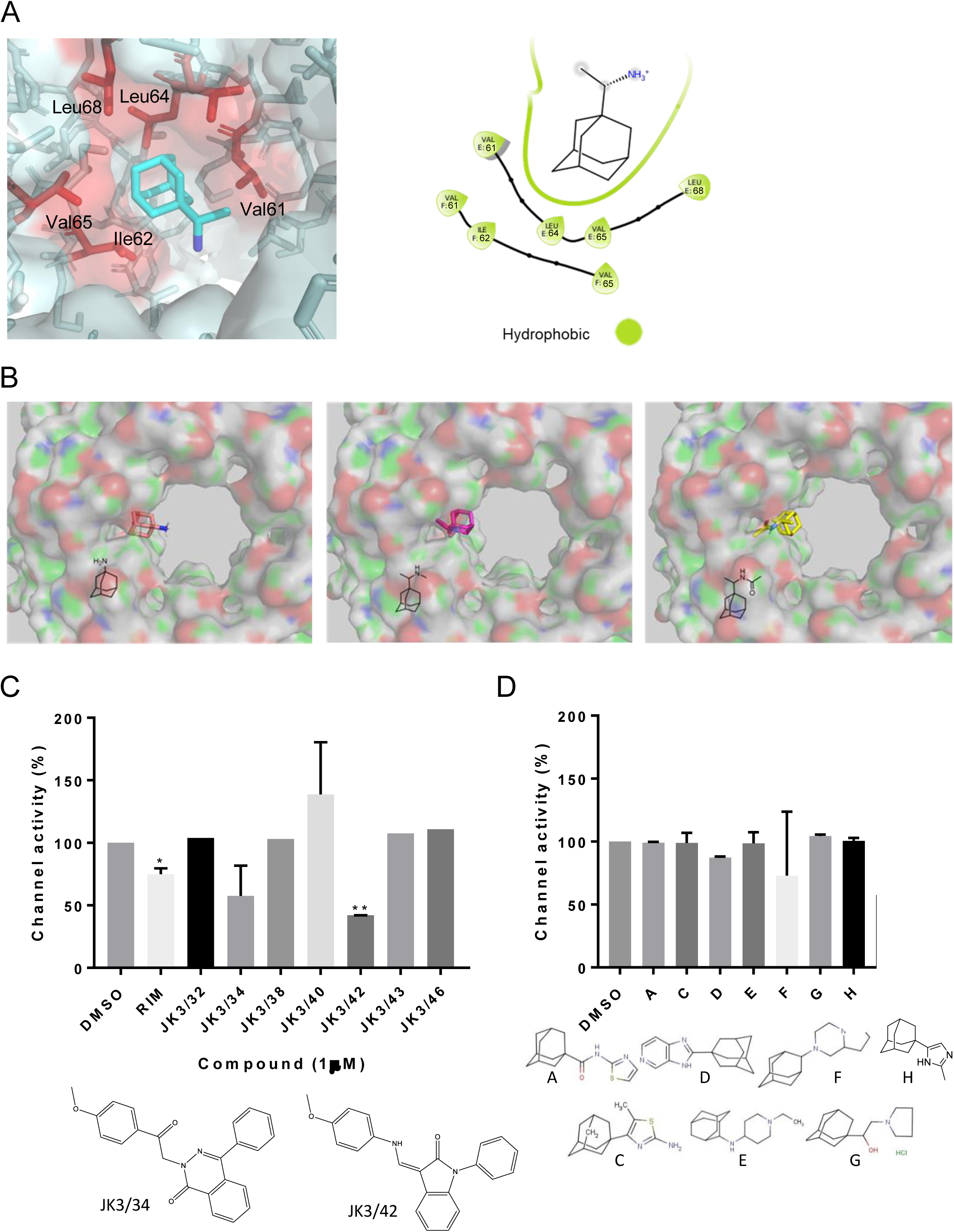
Lack of inhibitory activity for other adamantane derivatives versus M in CF release assays. **A, B.** Rimantadine and derivatives showed overlapping binding modes at the L1 binding site, but derivatives adopted opposite orientations such that the hydrophobic adamantyl cage was predicted to project into the lumen. **C.** Compounds targeting a peripheral site upon the HCV p7 channel complex were tested using CF release assays to ascertain the potential relevance of the ostensibly similar P1 site and would be unlikely to occupy the lumenal sites as a result; these have a more planar structure compared with rimantadine. Two compounds (JK3/34 and JK3/42) were effective at inhibiting M channel activity, distinct from the HCV series lead (JK3/32), which had no effect. **D.** In contrast, a series of adamantane derivatives shown previously to have variable efficacy against both M2 and p7 had no effect upon M channels in dye release assays.

**Figure S10.**
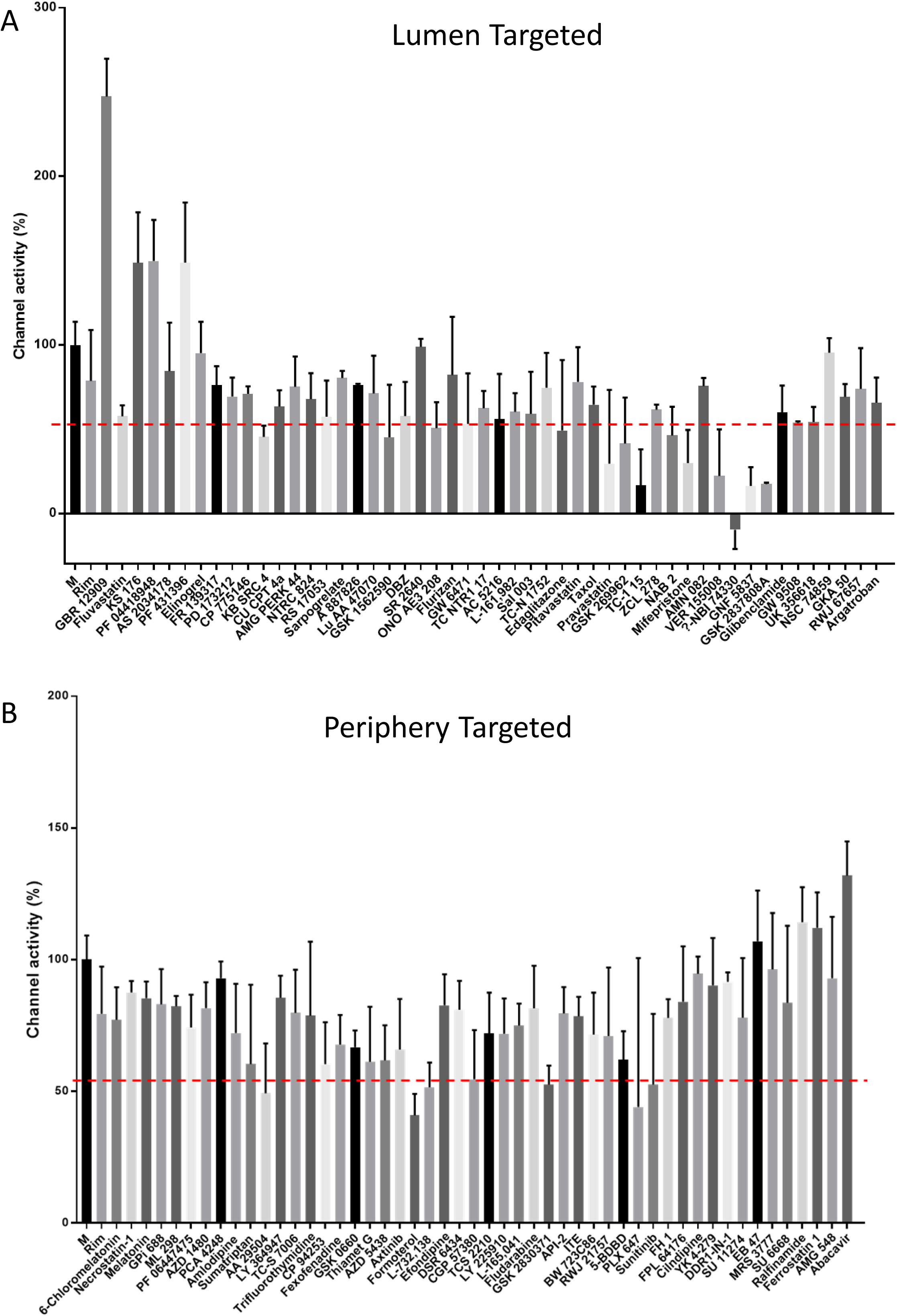
Screening data for lumenally (**A**) and peripherally targeted (**B**) M binding compounds, enriched using *in silico* HTS. The top 50 compounds based upon predicted binding scores and drug-like properties derived from the TOCRIS repurposing library were tested versus M at 1 μM using CF release assays. Results show the average of three biological repeats, with error bars representing the standard error of the mean.

**Figure S11.**
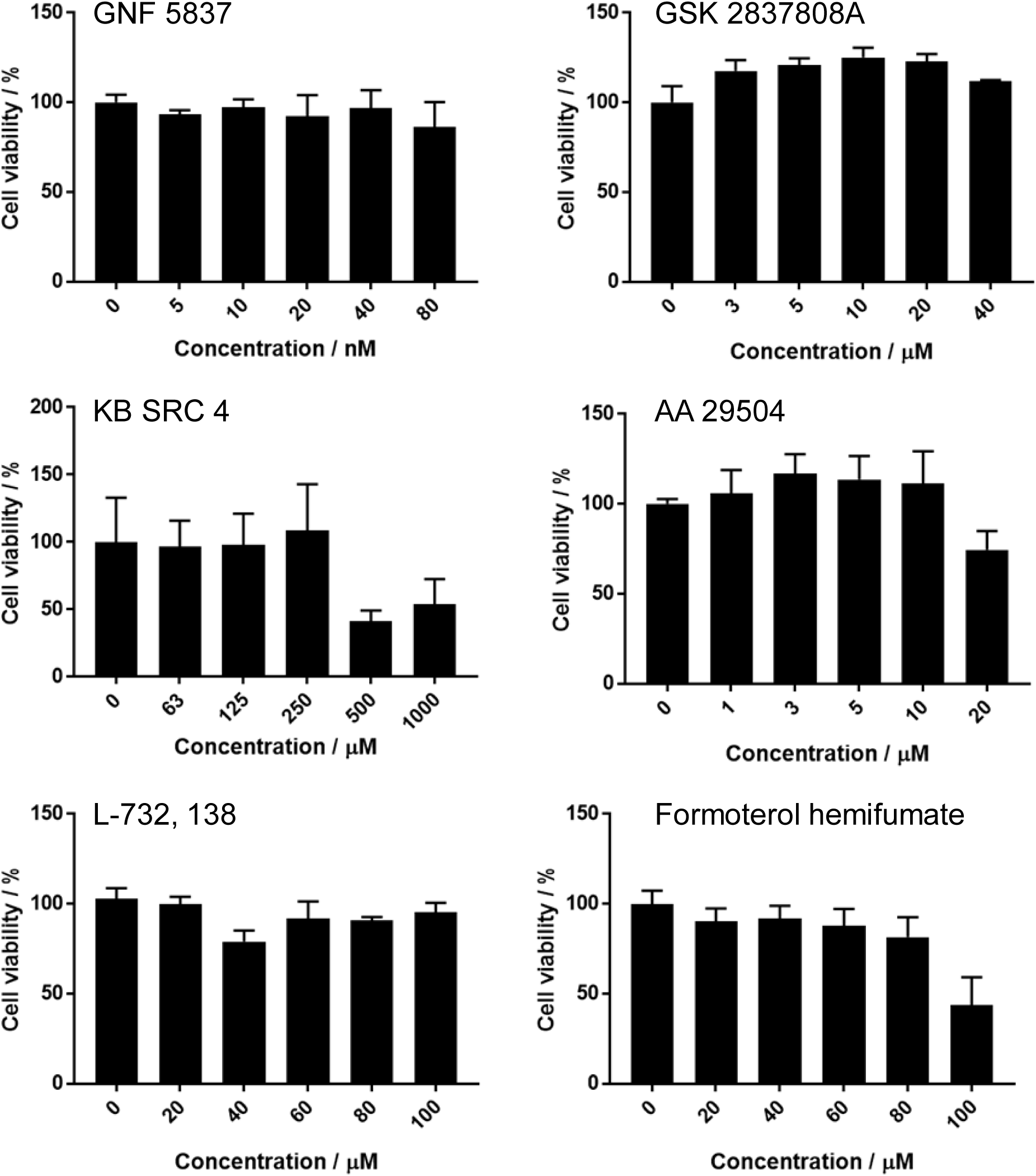
Effects of screening hits upon cell metabolic activity measured by MTT assay in Vero cells. Compound titrations were performed according to remain below cytotoxic doses described in the literature. Results show the average of three biological repeats, with error bars representing the standard error of the mean.

**Figure S12.**
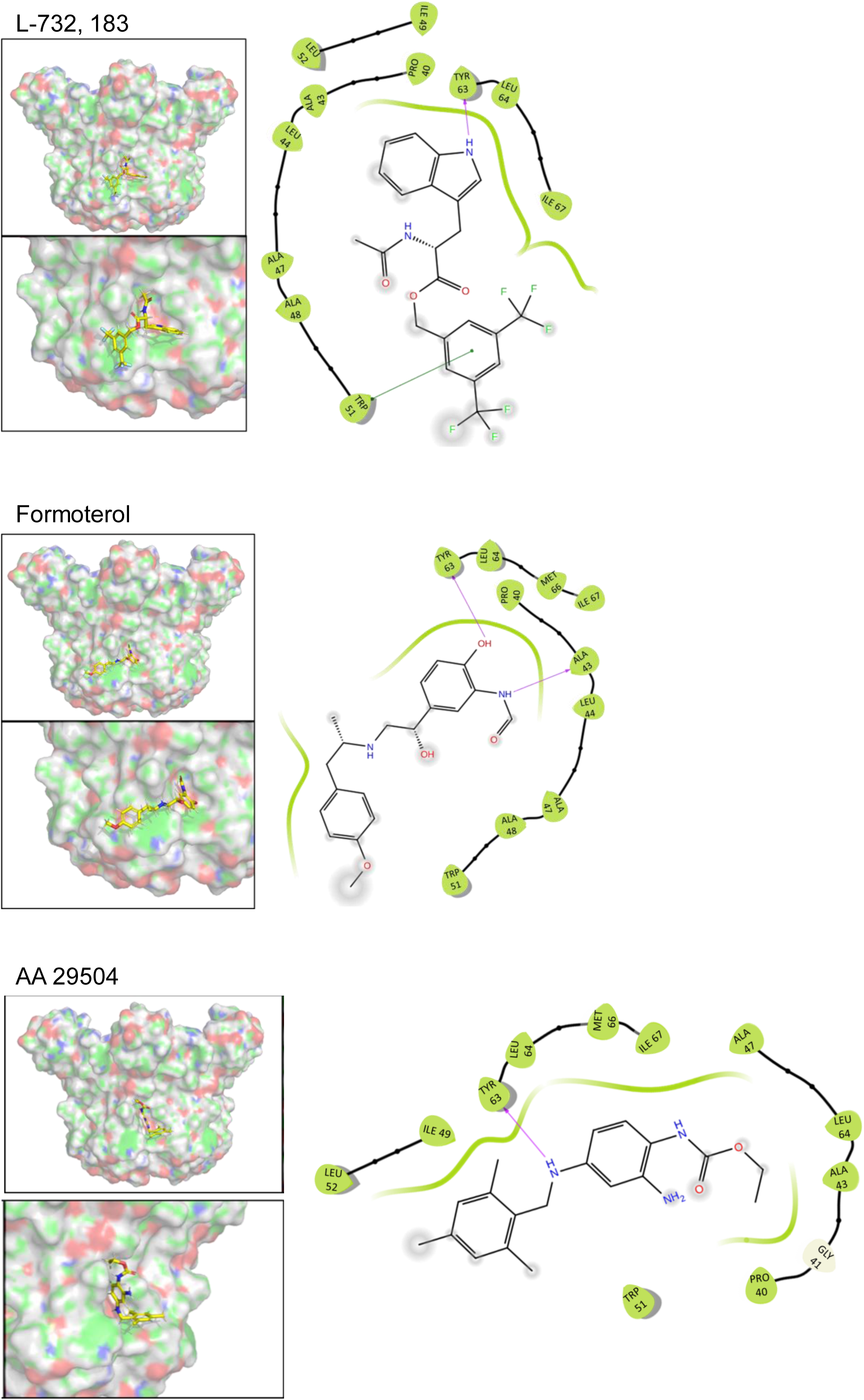
Predicted binding poses and residue interactions for peripherally targeted hit compounds.

**Figure S13.**
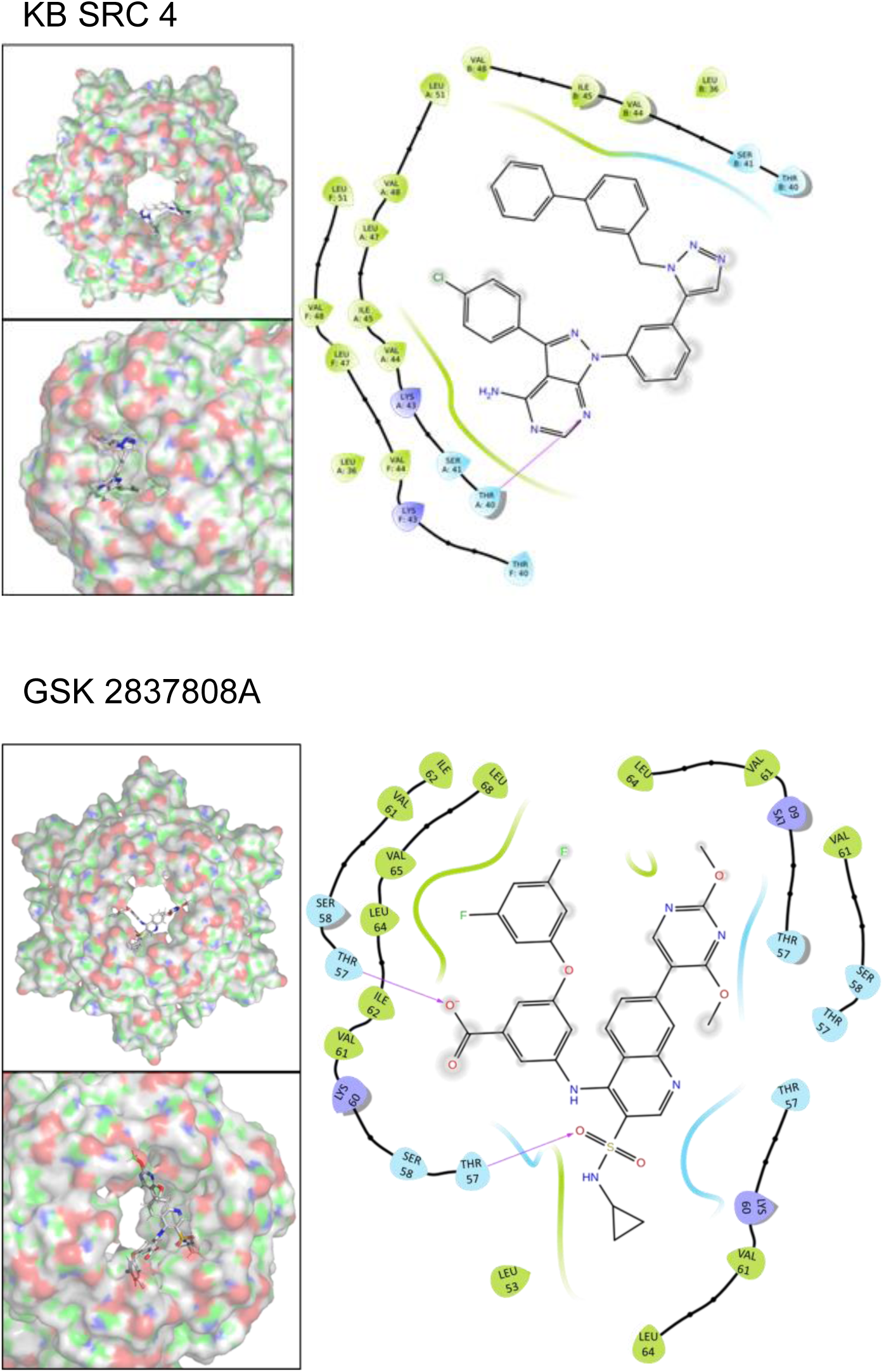
Predicted binding poses and residue interactions for lumenally targeted hit compounds.

**Figure S14.**
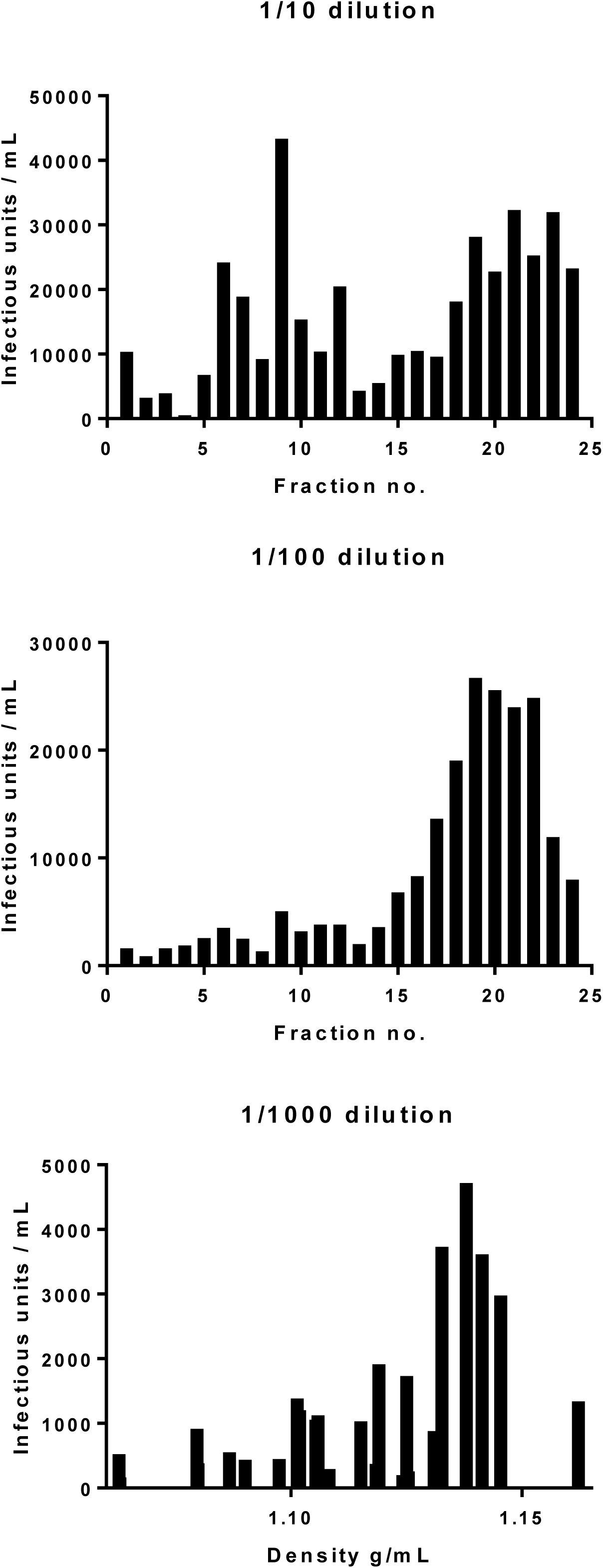
Effect of dilution upon measured titre for ZIKV purified through iodixinol gradients, as described in Fig 7. Identical gradients were titred upon Vero cells across a 10-fold dilution series. More concentrated innoculae (1:10) resulted in a detectable lower density peak, whereas further dilution (1:100, 1:1000) led to this diminishing whilst the higher density peak infectivity remained. Investigations into the composition of the low density peak are ongoing.

Annexe. Additional modelling solutions for oligomers with properties deemed less favourable compared to the compact hexameric complex lined by helix 3. This includes standard and protonated (His28^++^) simulations for radial arrangements of the complex, complexes lined by helix 2 as well as heptameric assemblages across all conditions.

Each model and simulation is depicted as overall structure (**A**), cut-away and top-down images of complexes highlighting lumenal regions (neutral only, **B**), representative endpoint conformation following simulations (**C**) and HOLE profiles describing channel lumenal apertures for start (blue) and final (red, green, black) conformations (**D**).

