## Supplementary material for "Inhibitors of the Small Membrane (M) Protein Viroporin Prevent Zika Virus Infection": Annexe detailing additional Channel MD simulations

### Annexe – alternative simulations for M channel complexes

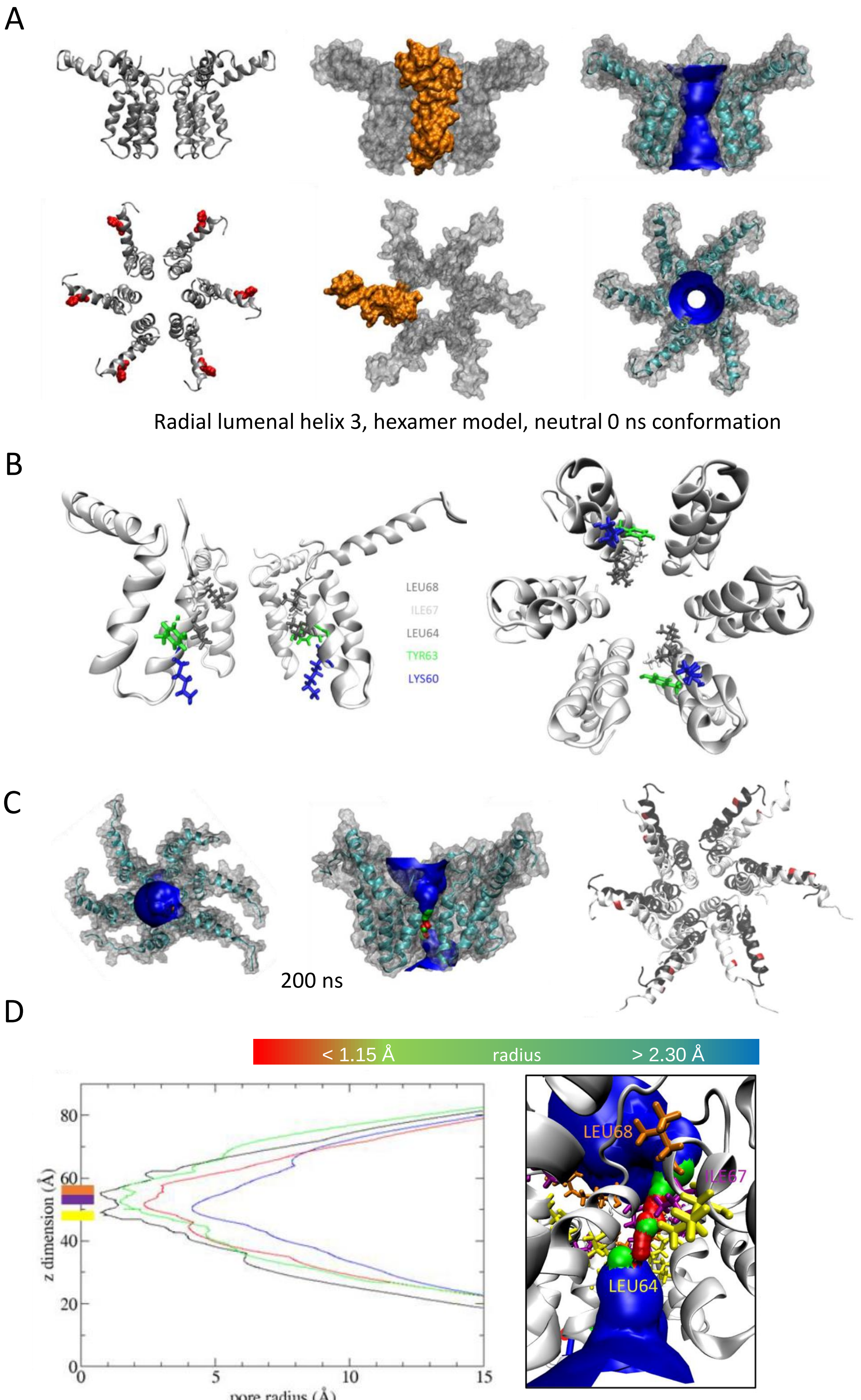


200 ns hole profile (neutral)

A

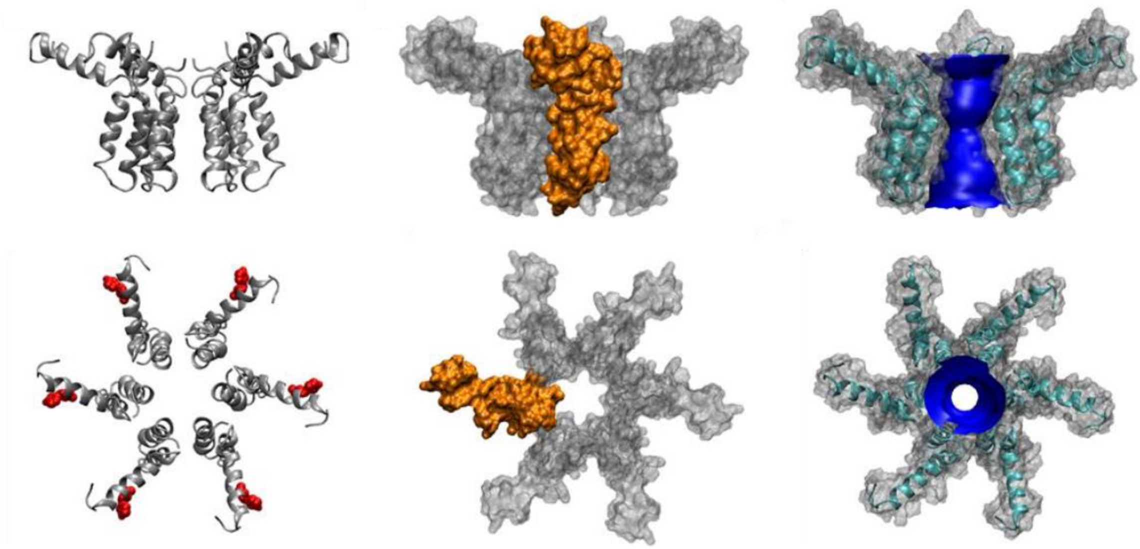

Radial lumenal helix 3, hexamer model, protonated 0 ns conformation

B

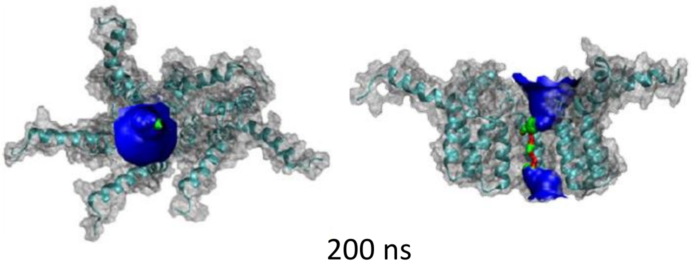

200 ns

C

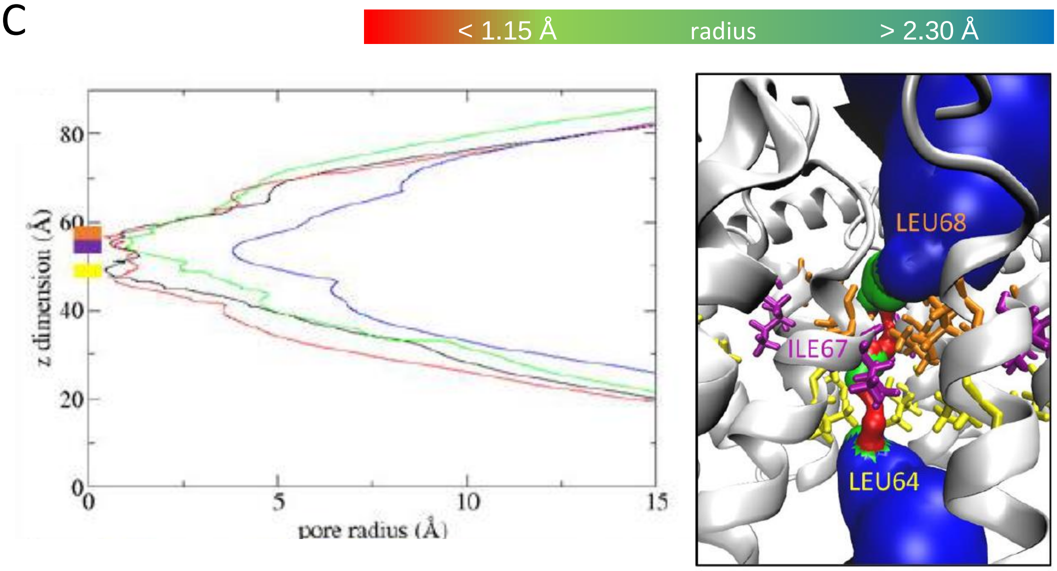


200 ns hole profile (protonated)

A

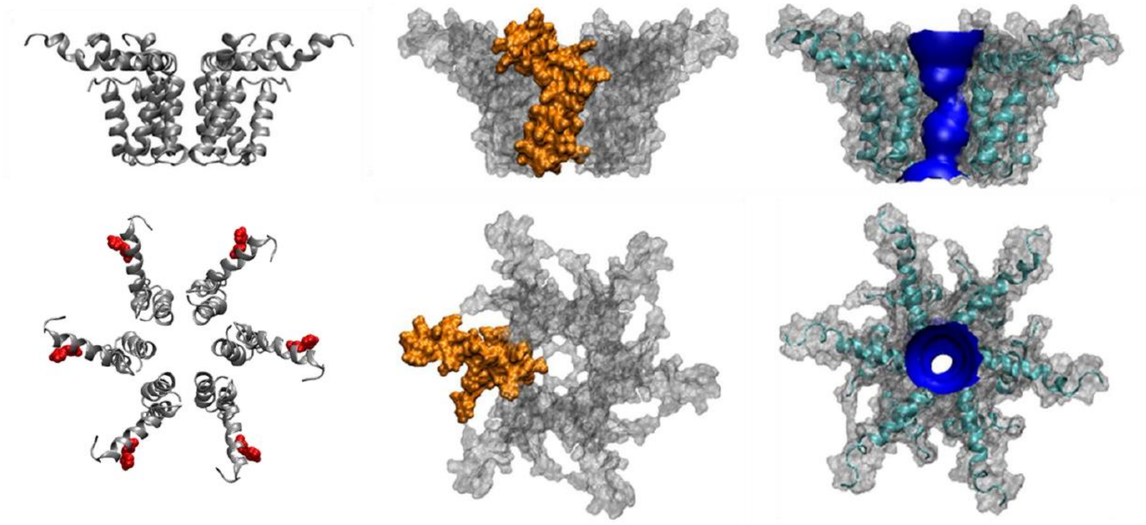

Radial lumenal helix 2, hexamer model, neutral 0 ns conformation

B

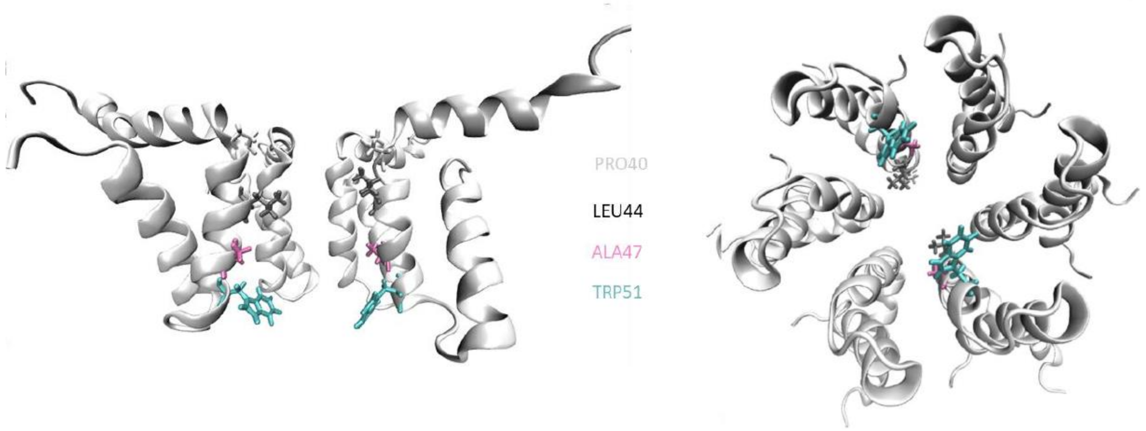


C

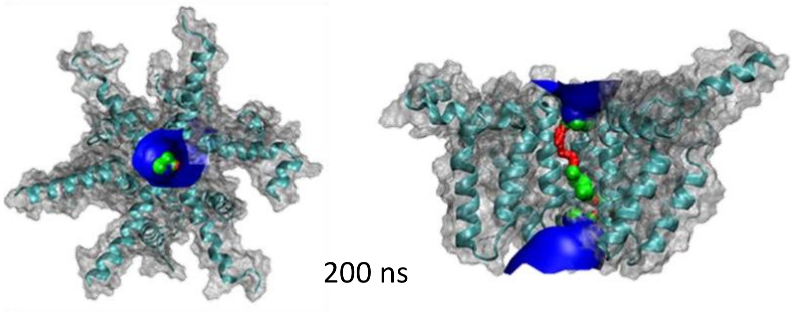


D

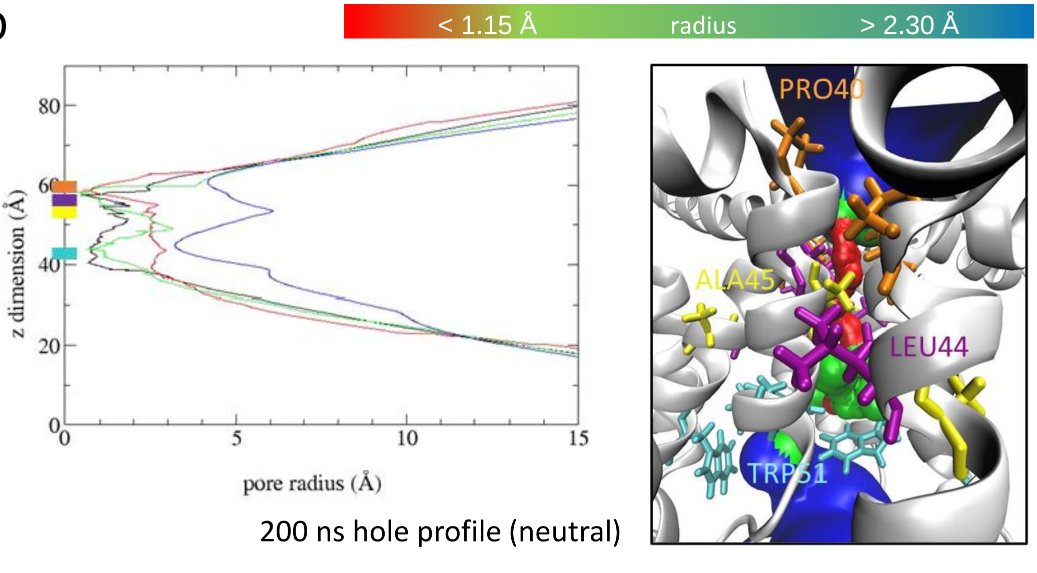


200 ns hole profile (neutral)

A

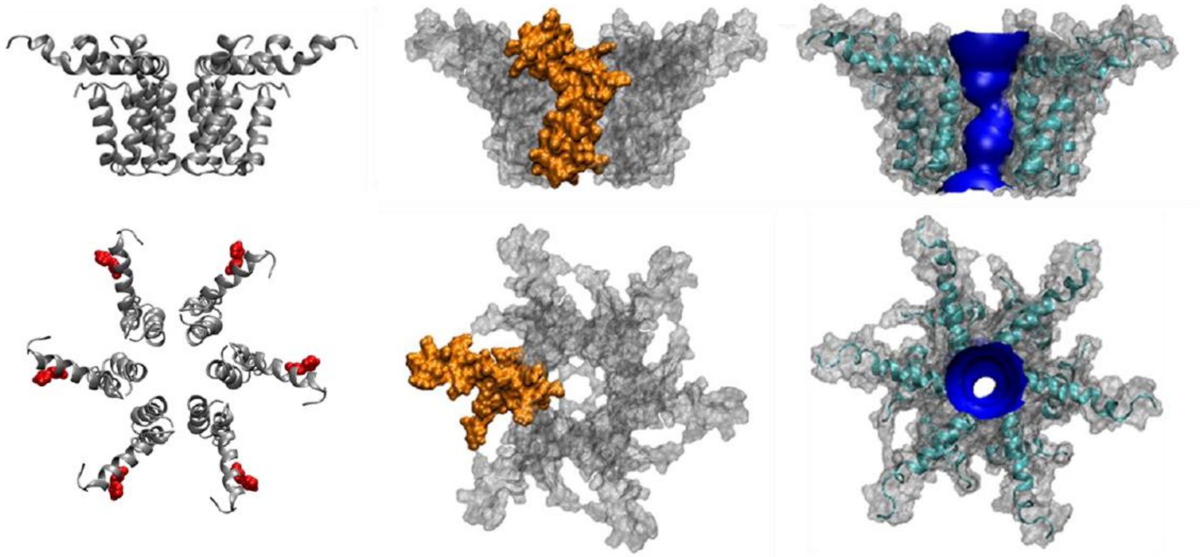

Radial lumenal helix 2, hexamer model, protonated 0 ns conformation

B

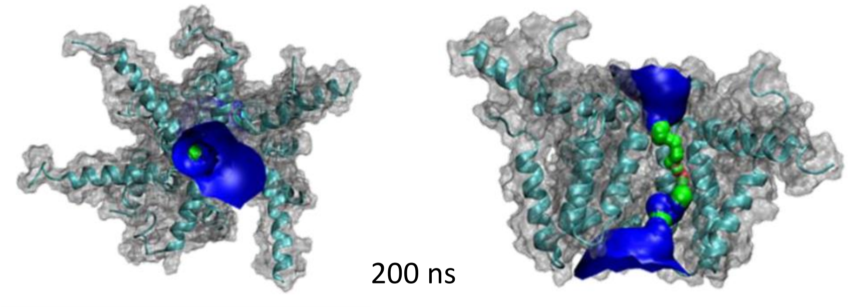

200 ns

C

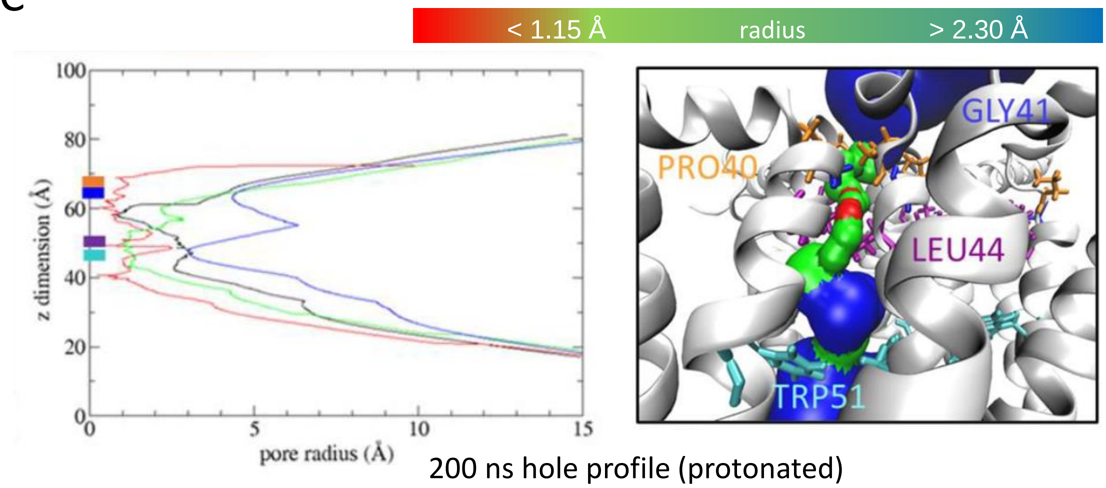


200 ns hole profile (protonated)

A

Annexe – 5

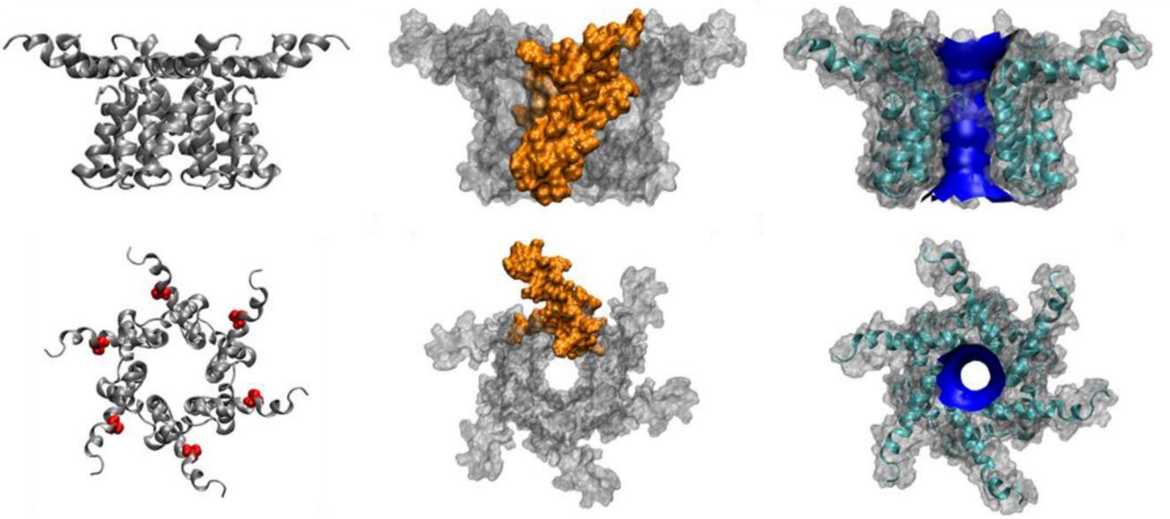

Compact lumenal helix 2, hexamer model, neutral 0 ns conformation

B

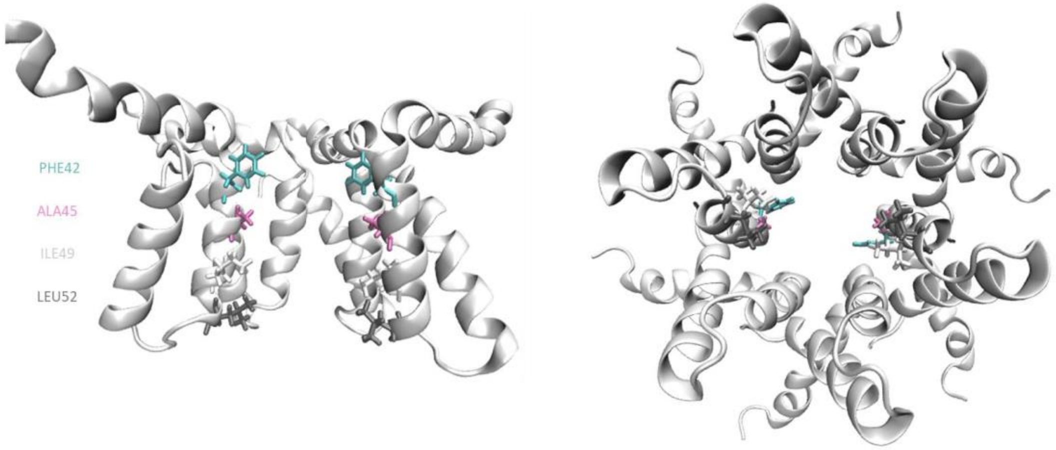


C

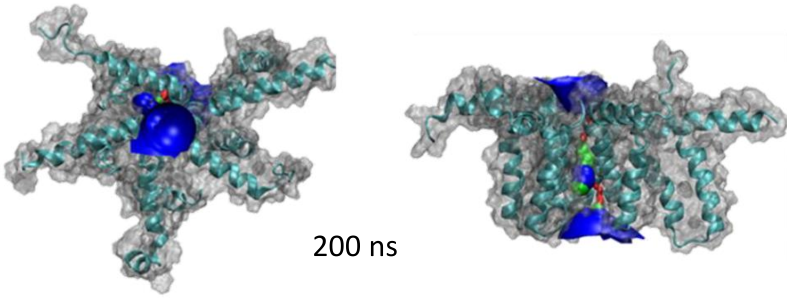


D

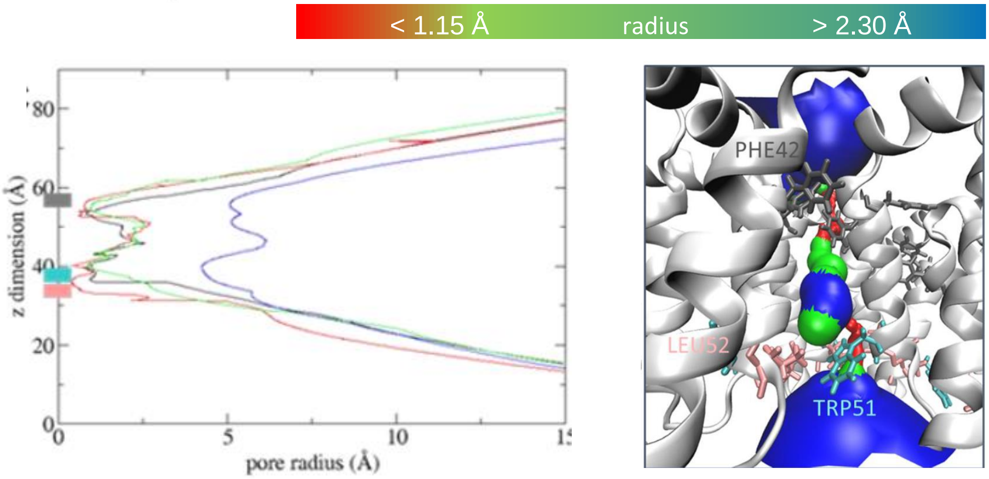


200 ns hole profile (neutral)

A

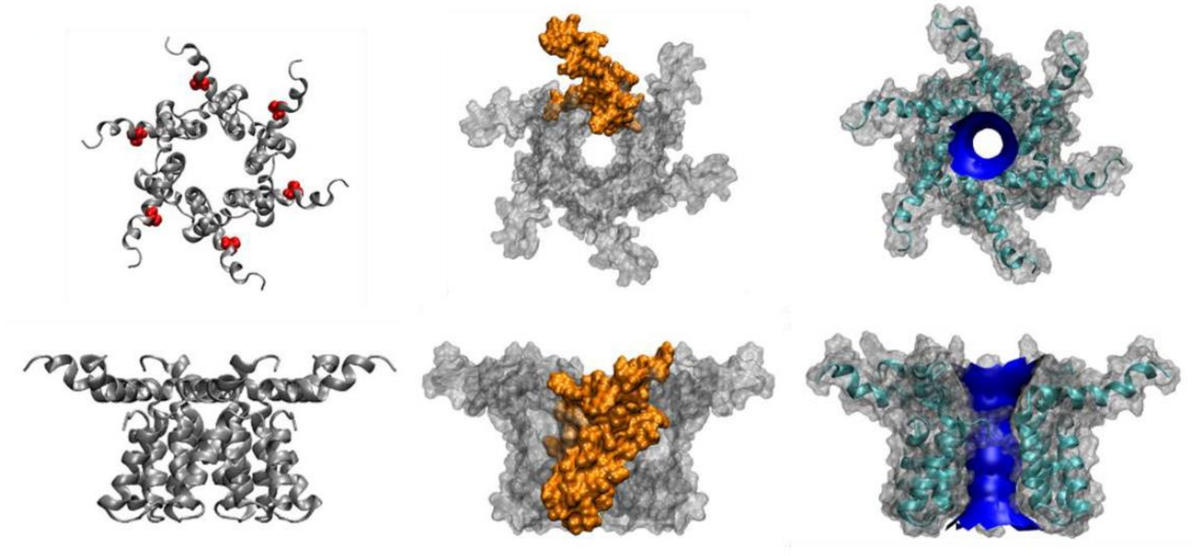

Compact lumenal helix 2, hexamer model, protonated 0 ns conformation

B

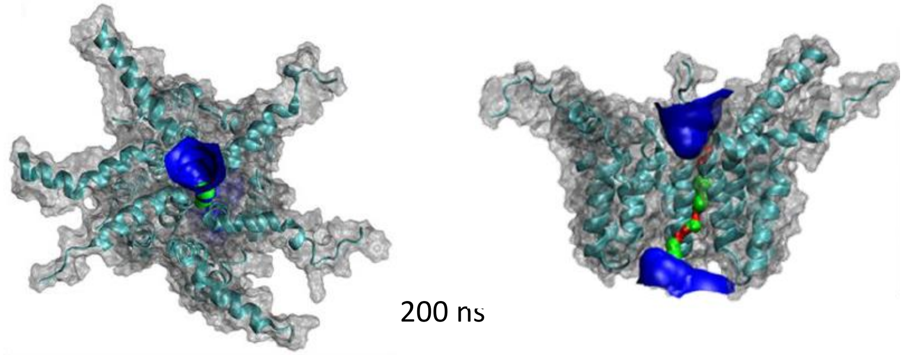


C

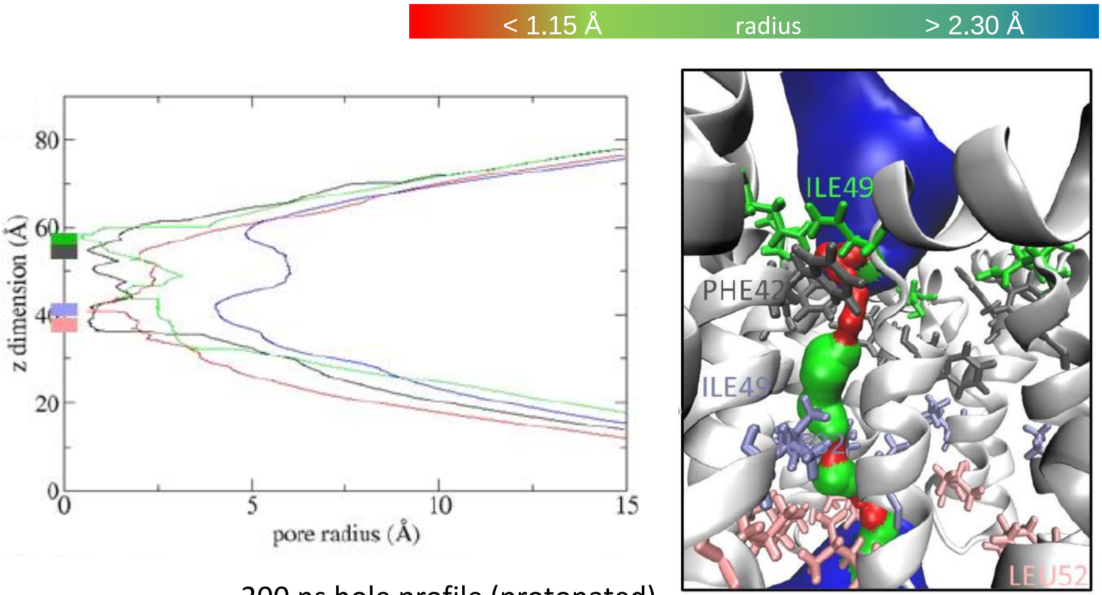


200 ns hole profile (protonated)

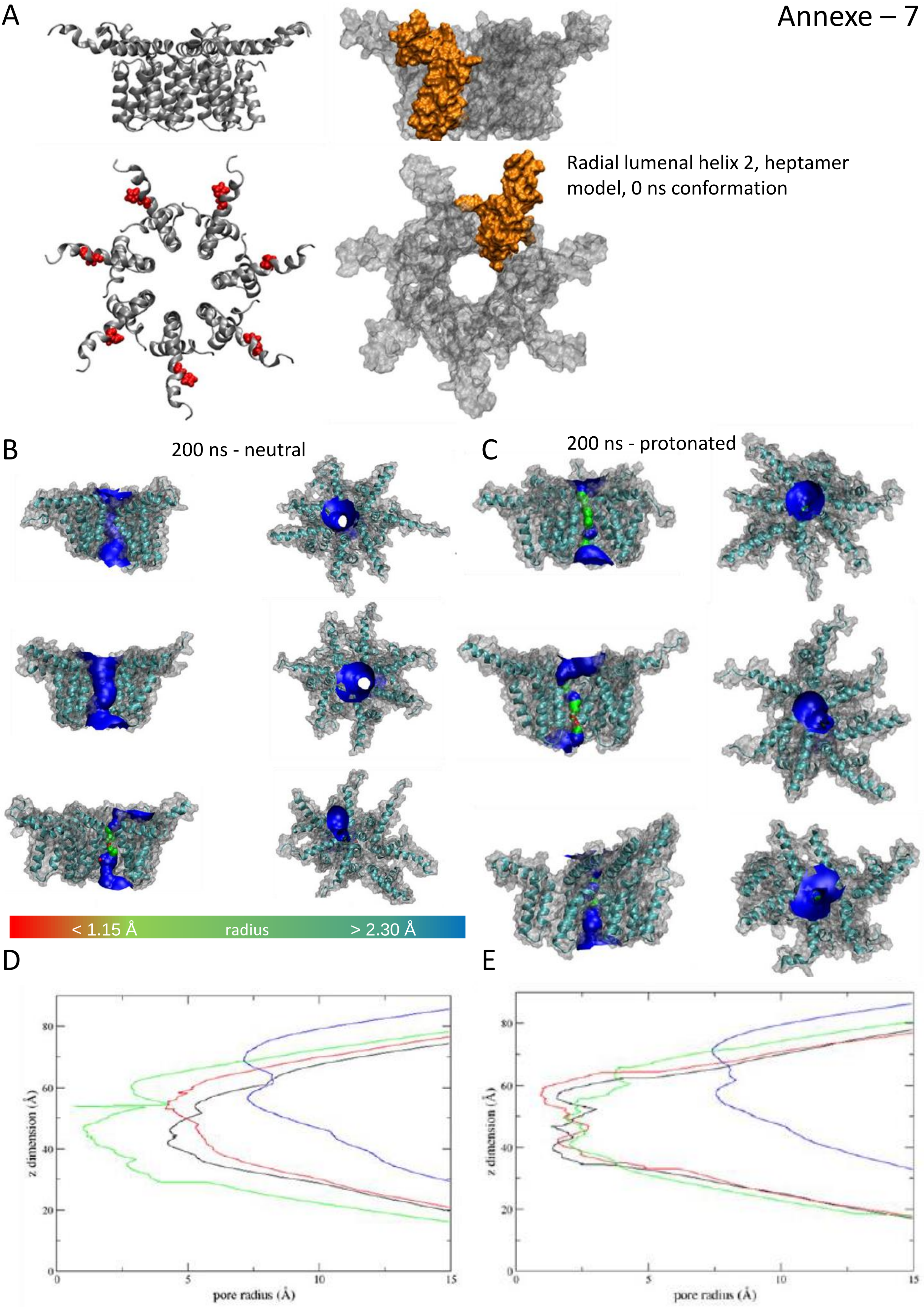


200 ns hole profile (neutral)

200 ns hole profile (protonated)

A

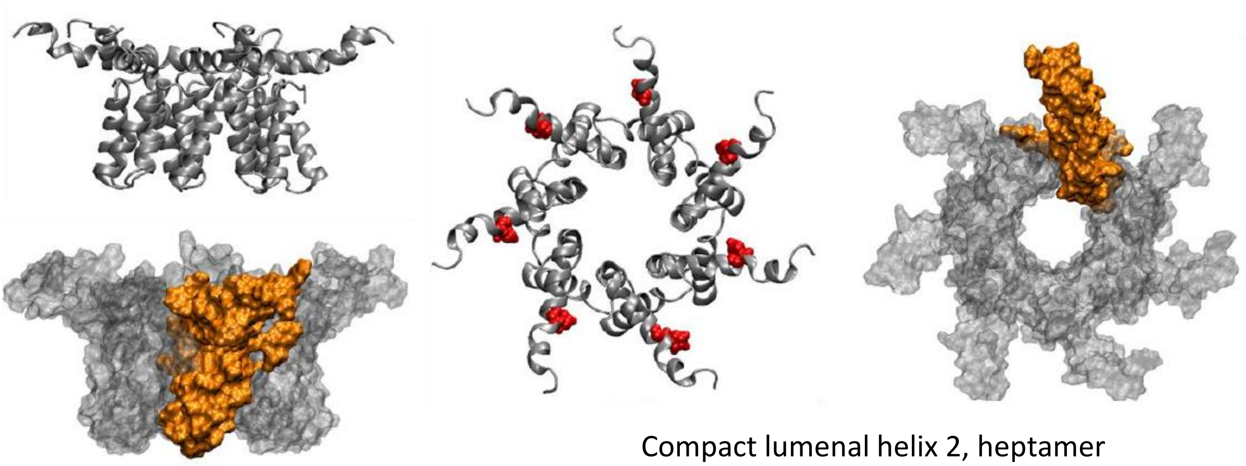

Compact lumenal helix 2, heptamer model, 0 ns conformation

B

200 ns - neutral

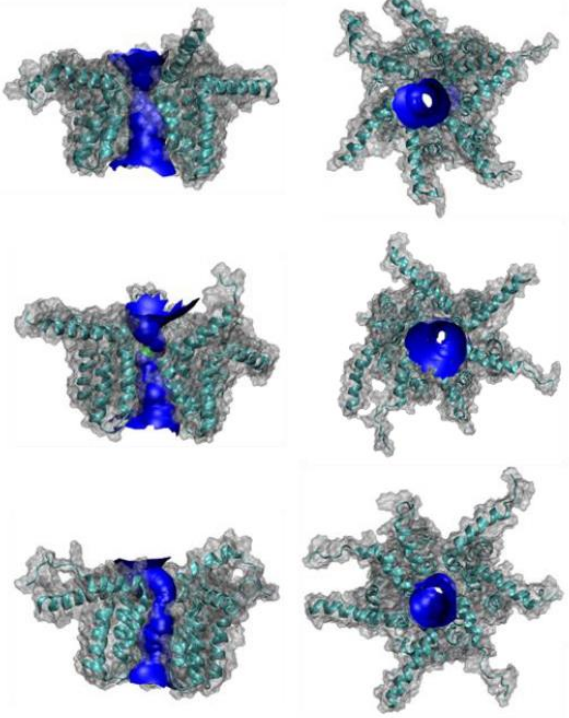

C

200 ns - protonated

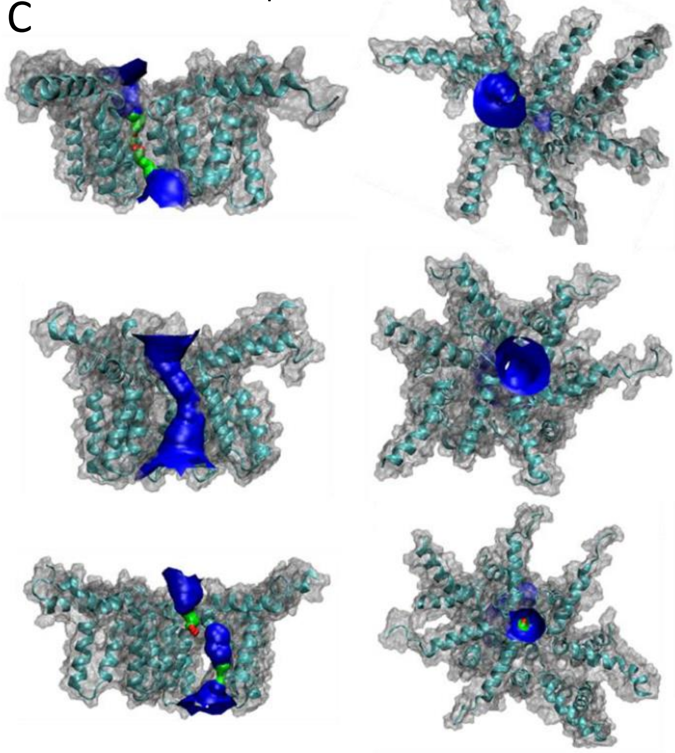

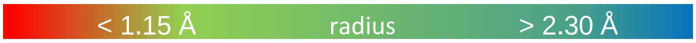


D

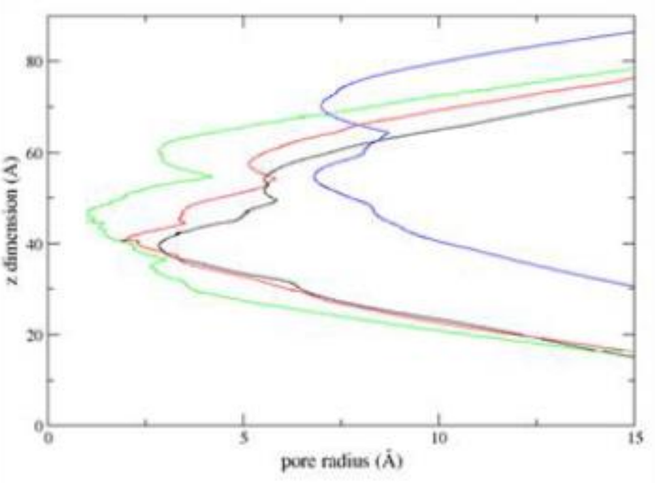


200 ns hole profile (neutral)

E

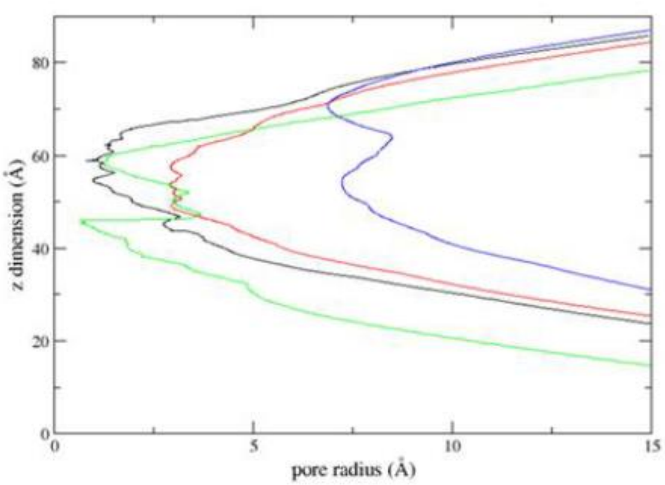


200 ns hole profile (protonated)

A

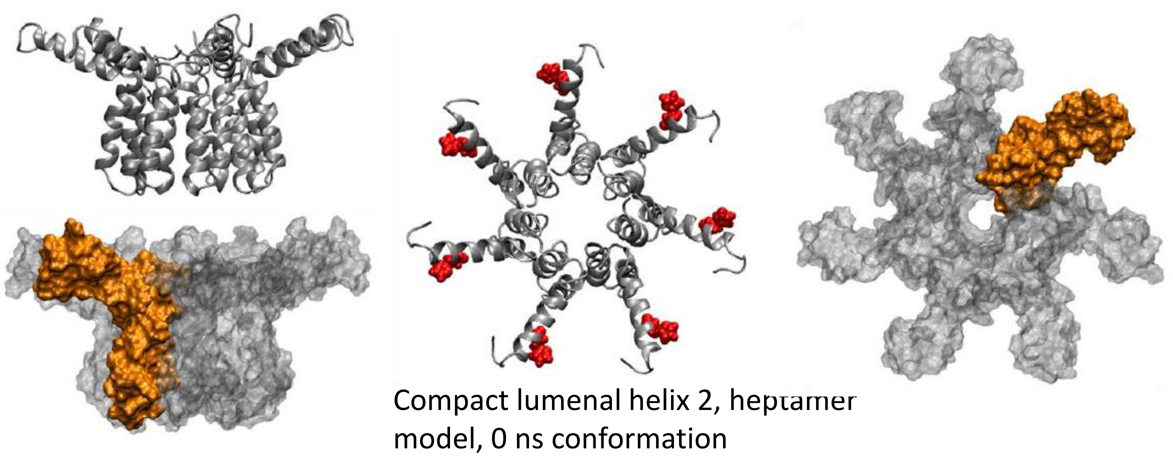

Compact lumenal helix 2, heptamer model, 0 ns conformation

B

200 ns - neutral

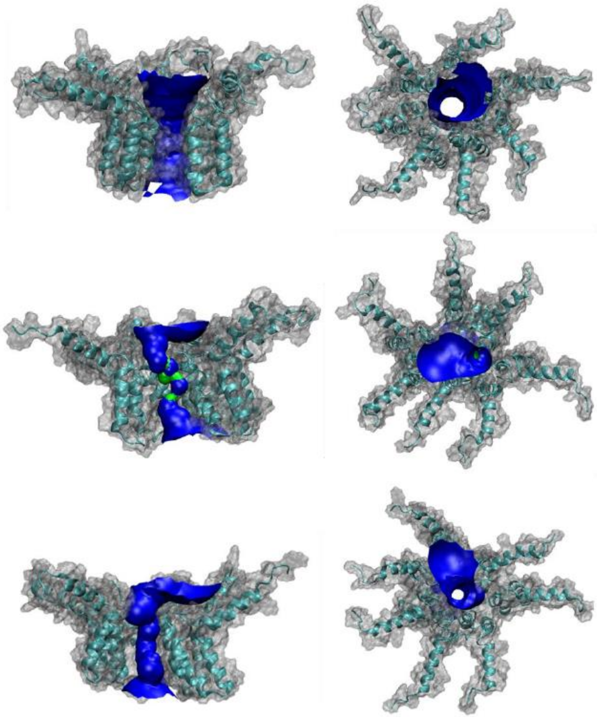

C

200 ns - protonated

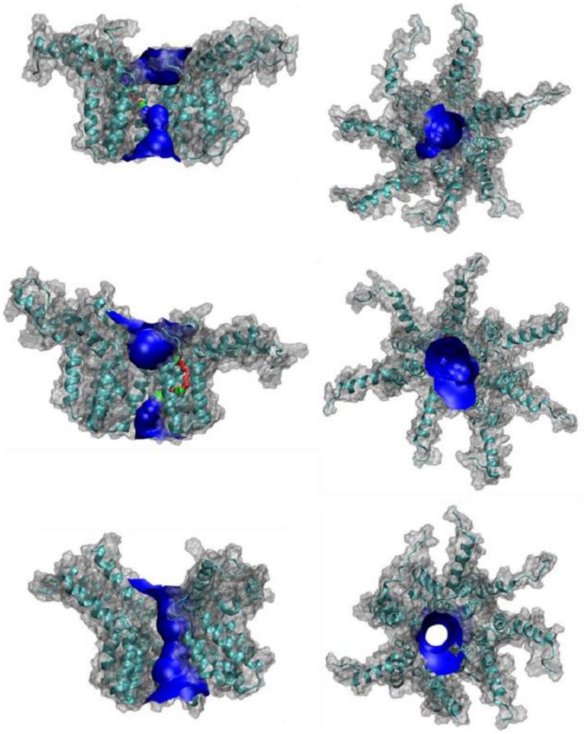

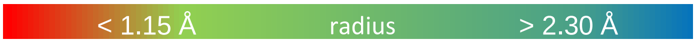


D

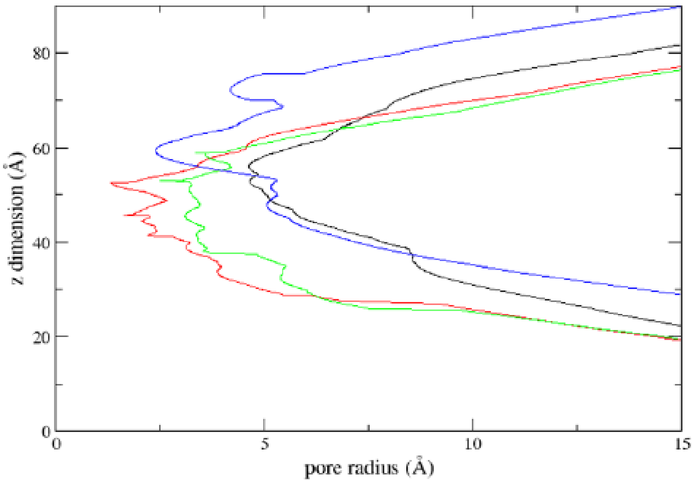


200 ns hole profile (neutral)

E

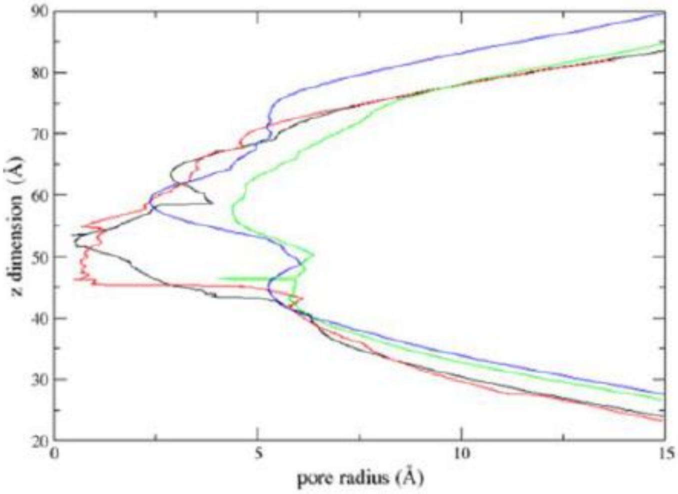


200 ns hole profile (protonated)

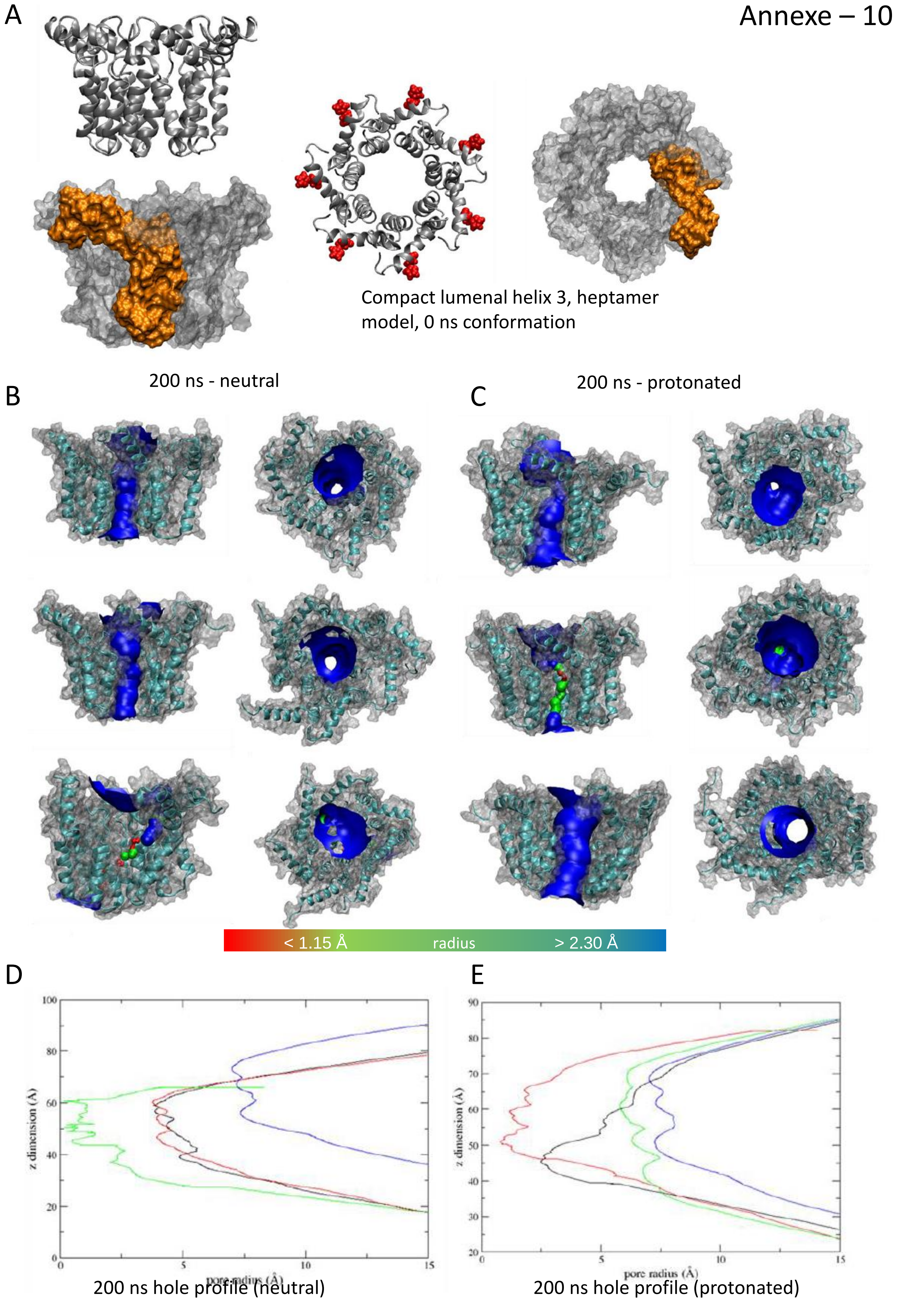
A
Compact lumenal helix 3, heptamer model, 0 ns conformation
B
200 ns - neutral
C
200 ns - protonated
< 1.15 Å
radius
> 2.30 Å
D
z dimension (Å)
pore radius (Å)
200 ns hole profile (neutral)
E
z dimension (Å)
pore radius (Å)
200 ns hole profile (protonated)
